# Manual Annotation of Genes within *Drosophila* Species: the Genomics Education Partnership protocol

**DOI:** 10.1101/2020.12.10.420521

**Authors:** Chinmay P. Rele, Katie M. Sandlin, Wilson Leung, Laura K. Reed

## Abstract

Annotating the genomes of multiple organisms allows us to study their genes as well as the evolution of those genes. While many eukaryotic genome assemblies already include computational gene predictions, these predictions can benefit from review and refinement through manual gene annotation. The Genomics Education Partnership (GEP; thegep.org) has developed an annotation protocol for protein-coding genes that enables undergraduate students and other researchers to create high-quality gene annotations that can be utilized in subsequent scientific investigations. For example, this protocol has been utilized by the GEP faculty to engage undergraduate students in the comparative annotation of genes involved in the insulin signaling pathway in 28 *Drosophila* species, using *D. melanogaster* as the informant genome. Students construct gene models using multiple lines of computational and experimental evidence including expression data (e.g., RNA-Seq), sequence similarity (e.g., BLAST, multiple sequence alignments), and computational gene predictions. For quality control, each gene is annotated by at least two students working independently, followed by reconciliation of the submitted gene models by a more experienced student. This article provides an overview of the annotation protocol and describes how discrepancies in student submitted gene models are resolved to produce a final, high-quality gene set suitable for subsequent analyses. This annotation protocol can be adapted to other scientific questions (e.g., expansion of the *Drosophila* Muller F element) and other species (e.g., parasitoid wasps) to provide additional opportunities for undergraduate students to participate in genomics research. These student annotation efforts can substantially improve the quality of gene annotations in publicly available genomic databases.

## Introduction

Genome annotation requires the assessment and integration of multiple lines of computational and experimental evidence. Several computational pipelines have been developed (e.g., BRAKER, MAKER) that can be used to construct an initial set of structural gene annotations for eukaryotic genomes (Holt and Yandell 2011; Hoff *et al.* 2019). Accuracy of the gene models produced by gene prediction algorithms depends on multiple biological factors (e.g., genome size, ploidy, repeat density, complexity of the transcriptome) and technical factors (e.g., quality of the genome assembly, evolutionary distance from the informant species, availability of transcriptome data) (Brůna *et al.* 2020). These factors can contribute to differences in the total number of gene predictions in closely related species (Figure 1).

**Figure 1:**
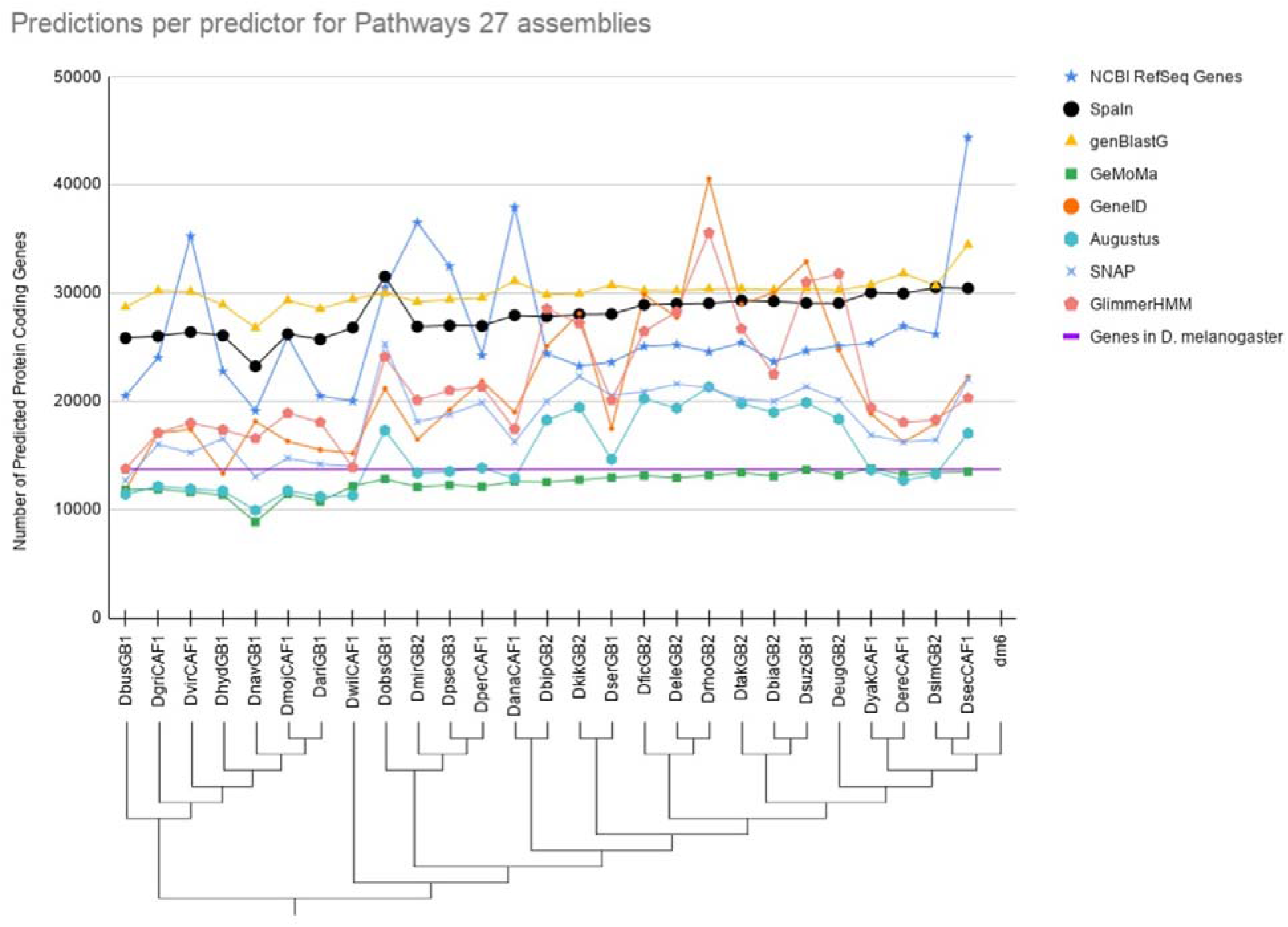
Variable Number of Gene Predictions. Variable number of gene predictions per gene predictor algorithm for each genome assembly for species in our clade (*Drosophila*). The number of genes predicted by different gene prediction algorithms differs depending on divergence from *D. melanogaster* for those predictors that use homology as their primary tool. The number of gene predictions varies drastically across algorithms and species. Some algorithms appear to consistently either underpredict or overpredict the number of genes relative to the well curated set of genes in *D. melanogaster* as maintained by FlyBase (purple line). The genome assemblies indicated in cladogram correspond to those mentioned in Table 1.

**Figure 2:**
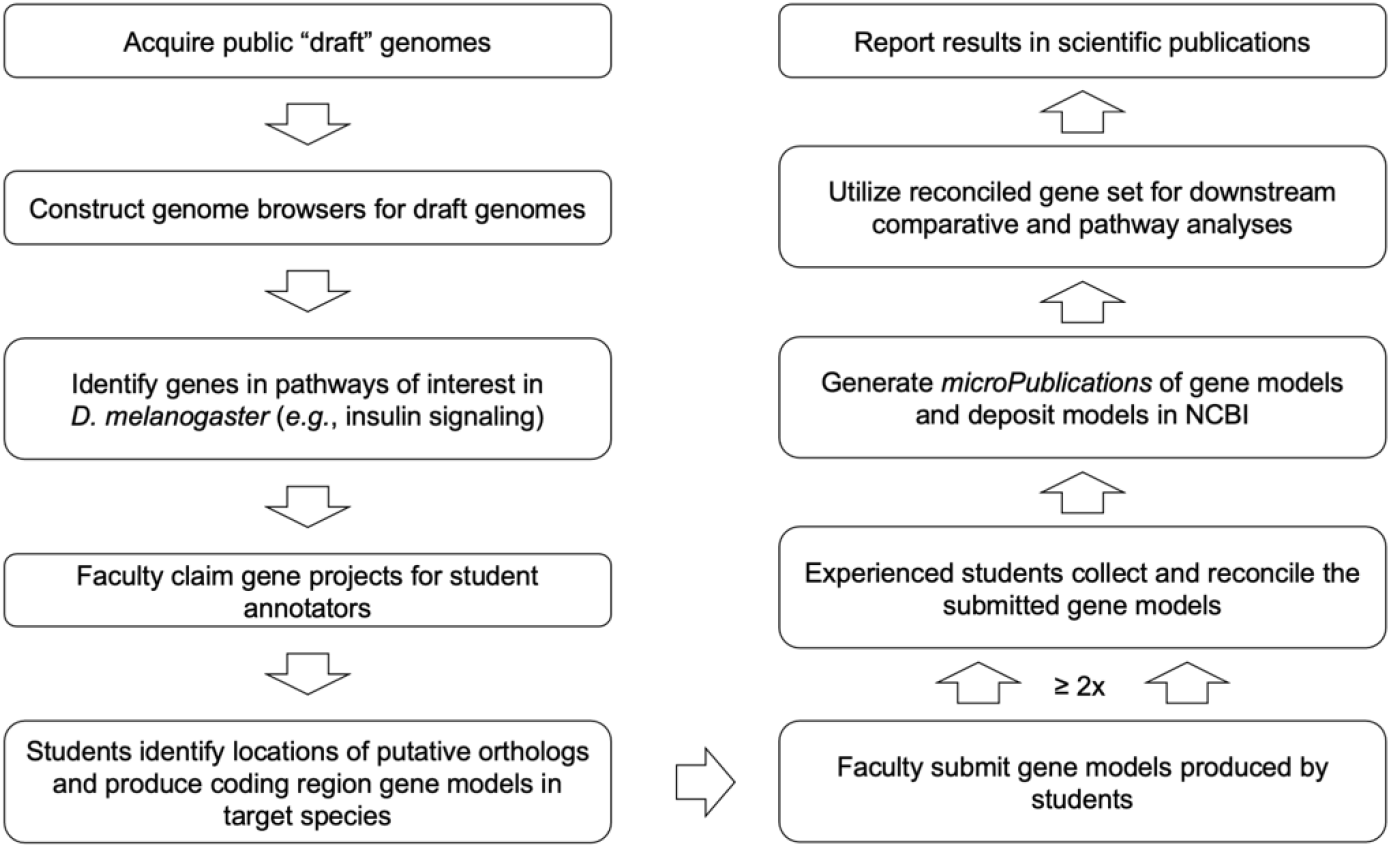
Overall Model Creation Workflow. Summarized workflow for annotation and reconciliation of a model to produce a finalized gene model

Evidence-based gene prediction algorithms (e.g., *Gnomon* (Souvorov *et al.* 2010)) use data, such as RNA-Seq derived from the same species, to predict genes within that assembly. The advent of high-throughput RNA sequencing (RNA-Seq) data has led to substantial improvement in the quality of gene annotations (Hoff *et al.* 2016; Keilwagen *et al.* 2018), particularly for species that lack high-quality gene annotations from a closely related informant species. However, the efficacy of assembling transcripts from RNA-Seq data depends on the expression level of the transcript in the developmental stages and tissues sampled by RNA-Seq (Shao and Kingsford 2017). Furthermore, long-read RNA sequencing, which is needed to resolve mRNA isoforms, has several challenges pertaining to RNA extraction methods, bias toward short transcripts, sequencing throughput, and read accuracy (reviewed in (Byrne *et al.* 2019)). Past studies have also shown that transcriptome constructed from long-read RNA-Seq data (e.g., from Iso-Seq by Pacific Biosciences and Direct RNA sequencing by Oxford Nanopore) has high sensitivity but low precision (Kovaka *et al.* 2019).

Consequently, despite recent advances in gene prediction algorithms and the increasing availability of RNA-Seq data, gene predictions produced by computational algorithms can still benefit from manual review and refinement (Mudge and Harrow 2016; Tello-Ruiz *et al.* 2019). This article describes a protocol, developed by the Genomics Education Partnership (GEP; thegep.org), to engage undergraduate students in the comparative annotation of protein-coding genes involved in the insulin signaling pathway across 28 species of *Drosophila* (including *D. melanogaster* as the reference species).

As of December 2020, GEP students from 35 institutions have used this annotation protocol to construct 1,114 gene models across 27 *Drosophila* species. Despite differing instructional settings and instructors, this protocol ensures that GEP students use a uniform standard to construct gene models that are best supported by the available evidence. As an additional level of quality control, each gene is annotated by at least two students working independently, and the submitted gene models are then reconciled by an experienced student using the *Apollo* genome annotation editor (Dunn *et al.* 2019).

These reconciled gene models will be described in *microPublication* articles (Raciti *et al.* 2018) and submitted to the NCBI Third Party Annotation (TPA) database (Clark *et al.* 2016). Researchers can utilize the high-quality, manually curated gene models constructed by GEP students to investigate gene and genome evolution.

## Methods

### Overview of the Coding Region Annotation Protocol

*Drosophila* researchers and curators at FlyBase have produced high-quality gene annotations for *D. melanogaster* based on large amounts of genetic and sequencing data (Matthews *et al.* 2015). Our protocol utilizes the high-quality and comprehensive gene annotations from *D. melanogaster* (informant species) to facilitate the annotation of the coding regions of orthologous genes in other *Drosophila* species (target species). In the absence of compelling evidence (e.g., RNA-Seq data) indicating significant differences in the gene model, the proposed gene model in the target species minimizes the number of changes compared to the ortholog in the informant species (i.e., construct the most parsimonious gene model assuming evolutionary conservation).

In order to generate manual annotations for the protein-coding sequence (CDS) of a particular gene in the target species, we need to (1) identify the ortholog of that gene from our reference species in our target species using sequence similarity and synteny, (2) determine the structure and approximate coordinates of each isoform and its coding exons, and then (3) refine those coordinates for each isoform. The key analysis steps are summarized in the Annotation Workflow for the Pathways Project (Supplement 1). A walkthrough illustrating each step of the annotation protocol on an example gene is available on the GEP website (Supplement 2).

The annotation and reconciliation protocols described below utilize multiple bioinformatics tools. The “Data (and Software) Availability” section at the end of the manuscript provides a brief overview of each tool, including version information and references.

#### CDS Annotation Procedure

##### Project Claiming

A GEP faculty member selects a *D. melanogaster* gene involved in the insulin signaling pathway (i.e., target gene in the informant genome) and one of the 27 *Drosophila* species (target species) for their students to annotate. A gene projects can be chosen by the faculty member to best suit their students and the pedagogical goals of the faculty member’s course based on the estimated difficulty of the project, as well as any specific interests in the biological function of the gene.

##### Identify the Ortholog

The ortholog assignment in the target species is based on the analysis of protein sequence similarity and synteny (i.e., the relative gene order and orientation) (Jun *et al.* 2009; Jahangiri-Tazehkand *et al.* 2017). The key analysis steps are summarized in Supplement 3.

To identify the ortholog of our target gene, we examine the genomic neighborhood surrounding the gene in the GEP UCSC Genome Browser (further described in the Software section below; https://gander.wustl.edu) in both the informant species and the target species. This includes identifying the orientation and nearest two upstream and downstream genes relative to our target gene. We also compare any nested genes to the locus surrounding our putative ortholog in the target species (Figure 3).

**Figure 3:**
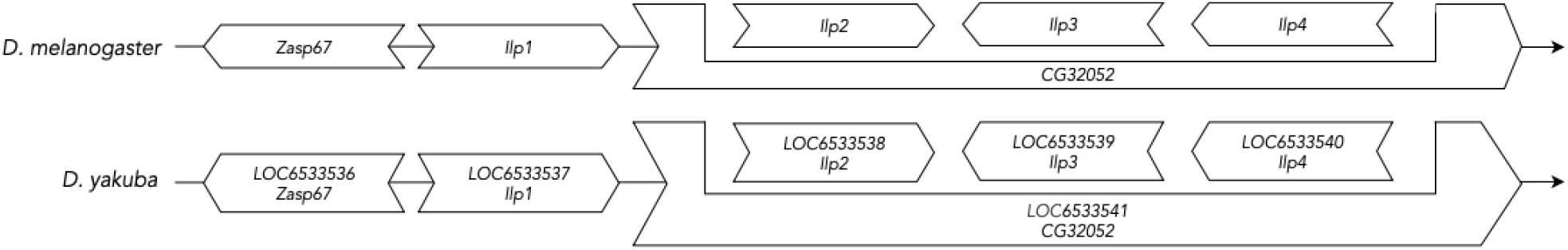
Synteny of Ilp2 in D. yakuba. Schematic of genomic neighborhood around the *Ilp2* gene in *D. melanogaster* and *D. yakuba*, showing that *Ilp2*, *Ilp3*, and *Ilp4* are nested within *CG32052.*

Locating the putative ortholog requires us to obtain the protein sequence for the target gene in our informant species from the *Gene Record Finder* and use it as the query to perform a *tblastn* search against the target species’ genome assembly. We perform *tblastn* against a specific assembly of interest on NCBI (https://www.ncbi.nlm.nih.gov/). We use *tblastn* since we are comparing our protein sequence query against a nucleotide database. Amino acid sequences are more conserved than nucleotide sequences across evolutionary time, due the degeneracy of the genetic code sequence similarity searches at the amino acid level are more sensitive than searches at the nucleotide level (States *et al.* 1991; Gonzalez *et al.* 2019). There are three possible outcomes of our *tblastn* search: (1) getting zero matches, (2) getting one match, or (3) getting more than one match.

A lack of *tblastn* matches could be caused by gaps or misassemblies in the genome assembly of the target species, or due to lack of sequence conservation between the orthologs. If this is the case, we can use the ortholog assignments for the flanking genes in the informant species and synteny (orthology of neighboring genes) to help establish the location of the ortholog for the target gene. If the location and candidate for the ortholog still cannot be established, the ortholog may not be present in the target species. The primary usage of our *tblastn* search is comparing the metrics of the best and second-best matches to confidently assign orthology. The presence of a single or multiple matches implies potential orthology that is then confirmed by comparing the local synteny of our target gene between the target and informant species. If there is no match for neighboring genes in an evolutionarily diverged species but a singular match exists for the target gene, there still may be an orthologous relationship between the reference and target sequences if they are reciprocal best hits in a BLAST search.

Figure 3 shows an example of how we can use synteny to establish the ortholog of the target gene. *Ilp2* in *D. melanogaster* and *D. yakuba* have the same genes surrounding them. A few of these genes are also nested within another gene, which further supports that the genomic neighborhood is conserved, and thus, the ortholog has been identified.

##### Identify the Approximate Coordinates of Each Coding Exon

Once the putative ortholog is identified, we begin constructing our gene model by separately mapping each coding exon to determine their approximate locations using the “Align two or more sequences” (*bl2seq*) feature provided by the NCBI BLAST web service.

For each isoform of the target gene in *D. melanogaster*, we performed *tblastn* searches of each coding exon of the isoform (query) in *D. melanogaster* against the scaffold which contains the putative ortholog of the target gene in the target species (subject). The amino acid sequence for each coding exon in the *D. melanogaster* isoform is obtained from the Gene Record Finder (further described in the Software section below). The scaffold in the target species which contains the putative ortholog of the target gene was determined from the “Identify the Ortholog” step above. For scaffolds larger than 10 Mbp, the “From” and “To” fields under “Subject subrange” in the NCBI *tblastn* search interface are used to limit the size of the search region to the approximate location of the target gene (inferred from results of the “Identify the Ortholog” step).

The default *tblastn* search parameters for NCBI Web BLAST are used in this search except for the following parameters:

1. Select the “Align two or more sequences” checkbox
2. Specify a subject subrange that corresponds to the coding span estimated at the “Identify the Ortholog” step
3. Select “No adjustment” under “Compositional adjustments”
4. Uncheck the “Low complexity regions” filter (under “Filters and Masking”)

At the end of this process, we have identified a collinear set of coordinates for most of the coding exons of the isoform. The results of these *tblastn* searches provide further supporting evidence for the ortholog assignment and provide anchors from which to narrow the size of the search region for identifying small or weakly conserved coding exons.

Since *tblastn* only aligns to complete codons, the BLAST alignments will not include partial codons adjacent to the splice junctions. In addition, *tblastn* does not take the locations of potential splice sites into account when it generates the BLAST alignment. Consequently, other lines of evidence (e.g., computational gene predictions, RNA-Seq data) must be used to refine the start and end coordinates of each coding exon.

##### Refine Coordinates

The coding exon coordinates determined by the *tblastn* searches are approximate as they only detect regions of similarity with complete codons and are only estimates of the portion of the coding exon which shows significant similarity to the coding exon in the informant genome. These coordinates do not account for intron splicing rules. We refine the collinear coordinates identified above utilizing the RNA-Seq track, other homology-based alignment algorithms, and gene predictions, as well as comparing the sequence to other species within the phylogeny (more details about the tracks used can be found in Supplement 4). If available, the RNA-Seq data can provide empirical support for the proposed gene model in the target species. Canonical splice sites (GT donor and AG acceptor) are adhered to unless there is good evidence to the contrary, such as spliced RNA-Seq reads from the target species and conservation of non-canonical splice sites across the clade. These splice sites were computed using HISAT (Kim *et al.* 2015) by aligning raw RNA-Seq read data within the Sequence Read Archive (Leinonen *et al.* 2011). We also need to ensure the donor and acceptor have compatible phases. A more detailed version of the workflow for coordinate refinement can be found within Supplement 5. Figure 4 gives an example coordinate refinement to account for the complete exon boundaries considering RNA-Seq data and splice site compatibility, and Table 2 shows the coordinates for the entire example gene along with their refined counterparts.

**Figure 4:**
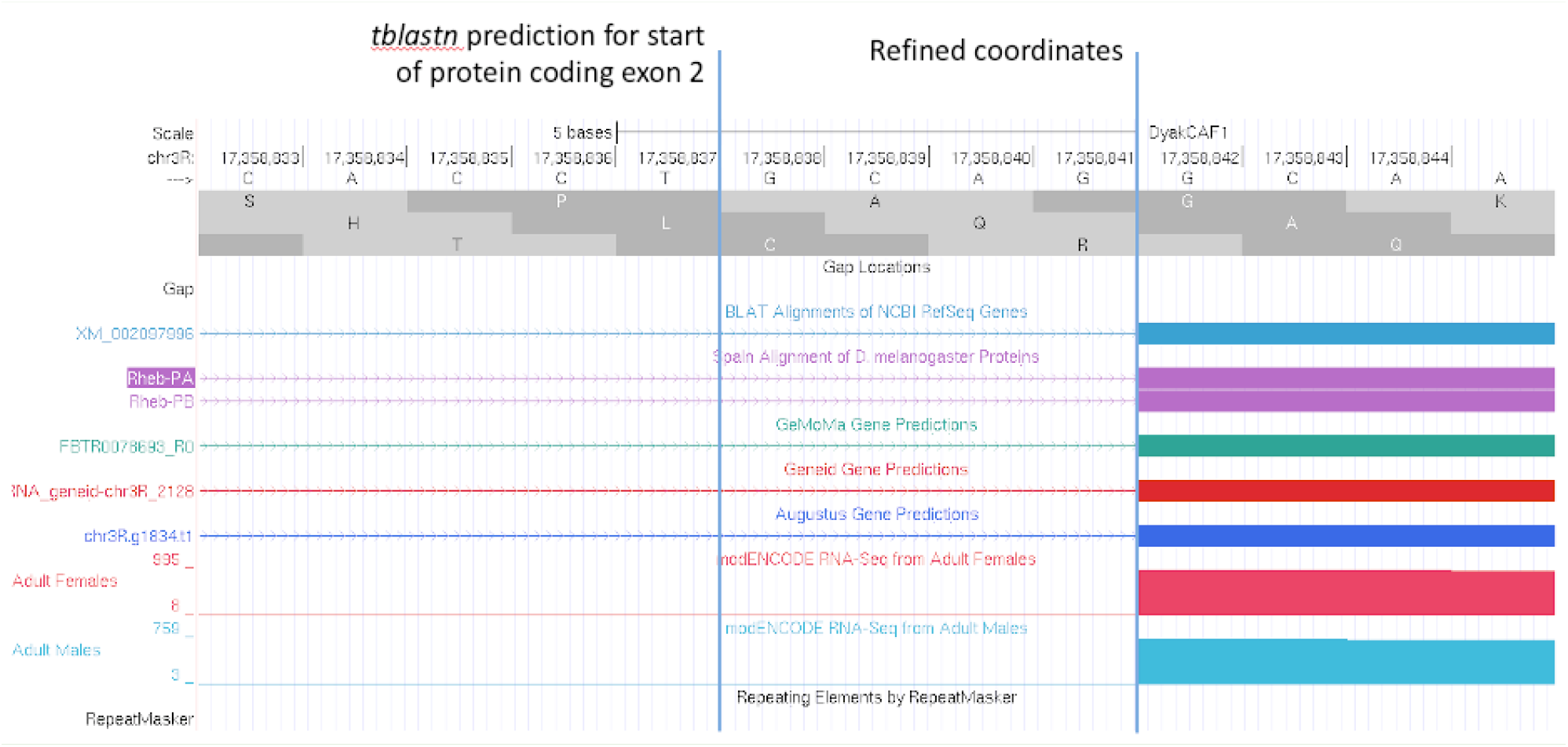
Refined Coordinates. Coordinate refinement for the second CDS of *Rheb* in *D. yakuba* since tblastn predicted the exon to be larger than it actually is based on other lines of evidence.

**Table 1:** Assemblies used for Insulin Signaling Pathway. Assemblies used for the Insulin Signaling Pathway project along with their NCBI RefSeq Accession Number and species names along with the accessions of the RNA-Seq assemblies used

| Assembly | NCBI RefSeq Accession | Species Name | RNA-Seq Assembly Accessions |
| --- | --- | --- | --- |
| DbusGB1 | GCA_001277935.1 | *D. buskii* | PRJNA274996 |
| DgriCAF1 | GCA_000005155.1 | *D. grimshawi* | SRP073087 |
| DvirCAF1 | GCA_000005245.1 | *D. virilis* | SRP006203 |
| DhydGB1 | GCA_002780465.1 | *D. hydei* | PRJNA373926 |
| DnavGB1 | GCA_001654015.1 | *D. navajoa* | NA |
| DmojCAF1 | GCA_000005175.1 | *D. mojavensis* | SRP006203 |
| DariGB1 | GCA_001654025.1 | *D. arizonae* | PRJNA395148 |
| DwilCAF1 | GCA_000005925.1 | *D. willistoni* | PRJNA388952 |
| DobsGB1 | GCA_002217835.1 | *D. obscura* | PRJDB4576 |
| DmirGB2 | GCA_000269505.2 | *D. miranda* | SRP009365 |
| DpseGB3 | GCA_000001765.2 | *D. pseudoobscura* | SRP006203 |
| DperCAF1 | GCA_000005195.1 | *D. persimilis* | PRJNA388952 |
| DanaCAF1 | GCA_000005115.1 | *D. ananassae* | SRP006203, SRP007906, PRJNA257286, PRJNA388952 |
| DbipGB2 | GCA_000236285.2 | *D. bipectinata* | PRJNA63469 |
| DkikGB2 | GCA_000224215.2 | *D. kikkawai* | PRJNA63469 |
| DserGB1 | GCA_002093755.1 | *D. serrata* | SRP096263 |
| DficGB2 | GCA_000220665.2 | *D. ficusphila* | PRJNA63469 |
| DeleGB2 | GCA_000224195.2 | *D. elegans* | PRJNA63469 |
| DrhoGB2 | GCA_000236305.2 | *D. rhopaloa* | PRJNA63469 |
| DtakGB2 | GCA_000224235.2 | *D. takahashii* | PRJNA63469 |
| DbiaGB2 | GCA_000233415.2 | *D. biarmipes* | PRJNA63469 |
| DsuzGB1 | GCA_000472105.1 | *D. suzukii* | PRJNA221549 |
| DeugGB2 | GCA_000236325.2 | *D. eugracilis* | PRJNA63469 |
| DyakCAF1 | GCA_000005975.1 | *D. yakuba* | SRP006203 |
| DereCAF1 | GCA_000005135.1 | *D. erecta* | PRJNA414017, PRJNA264407 |
| DsimGB2 | GCA_000754195.3 | *D. simulans* | SRP006203 |
| DsecCAF1 | GCA_000005215.1 | *D. sechellia* | PRJNA205470, PRJNA414017 |
| dm6 | GCA_000001215.4 | *D. melanogaster* | GSE117217 |

**Table 2:** Refining rough tblastn coordinates. Comparison between the *tblastn* and refined coordinates for the model of *Rheb* in *D. yakuba*

| | *tblastn* | | Refined | | $\Delta$ | |
| --- | --- | --- | --- | --- | --- | --- |
| Exon Number | Start | Stop | Start | Stop | Start | Stop |
| 1 | 17,358,666 | 17,358,725 | 17,358,666 | 17,358,666 | 0 | -11 |
| 2 | 17,358,838 | 17,358,927 | 17,358,842 | 17,358,913 | 4 | -14 |
| 3 | 17,359,007 | 17,359,216 | 17,359,011 | 17,359,218 | 4 | 2 |
| 4 | 17,359,279 | 17,359,407 | 17,359,407 | 17,359,407 | -1 | 0 |
| 5 | 17,359,467 | 17,359,556 | 17,359,470 | 17,359,556 | 3 | 0 |

After refining the coding exon coordinates, we use the *Gene Model Checker* to verify the coordinates for the proposed gene model satisfies the biological constraints for protein-coding genes in most eukaryotes, and reflects the structure of the *D. melanogaster* ortholog. The dot plot and protein alignment between the model and the *D. melanogaster* ortholog help us verify that our proposed gene model is the most parsimonious compared to the *D. melanogaster* ortholog. For more information on the *Gene Model Checker*, refer to the “Data (and Software) Availability” section.

We repeat the “Identify the Approximate Coordinates of Each Coding Exon” and the “Refine Coordinates” steps to construct gene models for each protein coding isoform of our target gene.

##### Final Submission

We then submit a GFF and FASTA (FAA and FNA) of each of the isoforms (see *Annotation Files Merger* tool below) as well as a complete Annotation Report (Supplement 6) to document the evidence supporting the proposed gene model.

#### Exceptions to the Standard Annotation Workflow

There are many challenges that can require the use of different annotation strategies than the basic annotation protocol outlined above. Gene models with these exceptions require support from multiple lines of evidence (e.g., RNA-Seq data, congruence with multiple gene predictors, conservation with *D. melanogaster* and other species in the clade). Below we describe the process for two such more common challenges:

##### Non-canonical Splice Site

GT and AG are the most commonly used splice donor and acceptor sites, respectively, and variant splice sites are termed non-canonical. For example, the GC splice donor site appears in ~1% (604 / 71,909) of the unique introns in *D. melanogaster* (FlyBase release 6.34). The use of a non-canonical splice site in a gene model will typically be supported by splice junction predictions derived from spliced RNA-Seq reads. The presence of a non-canonical splice site in the orthologous intron in the informant species (*D. melanogaster*) or in multiple *Drosophila* species more closely-related to the target species could also be used to support the annotation of a non-canonical splice site.

##### Assembly Errors

Each sequencing platform (e.g., Sanger, Illumina, PacBio, Nanopore) has a distinct error profile (Huang *et al.* 2012; Yang *et al.* 2017) that could introduce errors (e.g., base substitutions, insertions, and deletions) into the consensus sequence of an assembly. Transposons and other repetitive sequences (e.g., tandem repeats) in eukaryotic genomes can also lead to gaps and misassemblies (Mikheenko *et al.* 2018). These assembly issues can have substantial impacts on gene annotations — leading to frame shifts within coding exons, partial genes, missing genes or isoforms, and errors in ortholog/paralog assignments (e.g., due to “duplications” caused by misassemblies).

The publicly available genome assemblies our projects utilize are constructed using different sequencing technologies and assembly protocols. Since these genome assemblies have not been manually improved, they might contain assembly errors that could interfere with coding region annotations. Consequently, in cases where the proposed gene models for the target species include changes in gene structure compared to the *D. melanogaster* ortholog (e.g., novel/missing isoforms, exons, and/or introns), further investigations are needed to ascertain if the difference is caused by an assembly error, or reflects true divergence. As part of this assessment, we evaluate multiple lines of evidence such as: (1) conservation across other species in the *Drosophila* phylogeny, (2) consistency with genomic reads and RNA-Seq reads in the NCBI Sequence Read Archive (SRA) database, or (3) comparison with other genome assemblies for the same species (Miller *et al.* 2018).

##### Other Exceptions

Other potential challenges include whether the orthology of the gene in the target species has been correctly assigned in instances such as changes in gene structure relative to the reference species, absence of an isoform in the target species, or presence of a novel paralog are observed. We might also have cases where we have a non-canonical start codon (e.g., *Akt1*), stop codon readthrough (e.g., *jim*), or trans-splicing (e.g., *mod(mdg4)*), none of which conform to a standardized genetic ruleset. Challenges caused by assembly errors could be partial genes due to either gaps in the assembly, or the gene being near the end of a scaffold, as well as potential frameshifts within CDSs caused by errors in the consensus sequence.

#### Common Issues in Gene Models Prior to Submission

##### Multiple Fails in the Gene Model Checker Checklist

Gene models sometimes produce multiple “fails” for specific biological rules that were violated in the *Gene Model Checker*. Typically, fails are attributed to errors in upstream regions of the gene model that propagate downstream.

One common cause of multiple failures is frameshifts caused by selecting incompatible donor and acceptor splice sites. When a coding exon ends in an incomplete codon, the 3’ end of that coding exon and the 5’ beginning of the nearest downstream coding exon must include nucleotides that form a complete codon once the intron has been spliced out. The number of nucleotides between the end of the last complete codon and the splice donor site is defined as the phase of the splice donor site. Similarly, the number of nucleotides between the splice acceptor site and the start of the first complete codon is defined as the phase of the splice acceptor site. In order to maintain the open reading frame after the intron has been spliced out, the sum of the donor and acceptor phases for adjacent coding exons must either be zero (no extra codon) or three (one extra codon). Selecting incompatible donor and acceptor splice sites causes a frameshift that changes the reading frame of any coding exons downstream of that splice junction, thereby triggering multiple fails in the *Gene Model Checker*.

To resolve multiple fails, start troubleshooting at the beginning of the gene. In many cases, correcting errors in the upstream portion of the gene model resolves the fails reported downstream.

### Reconciliation

While most gene models produced by students using the annotation protocol described above are congruent with each other, incongruent models require further examination by a student reconciler. Reconciliation is carried out by experienced students who have received additional training on reconciliation under the guidance of a faculty and/or senior-scientist mentor.

The strategy of collecting at least two student models for each gene of interest in a target species and then comparing them with each other to reconcile a final model is essential for quality control. Student reconcilers look for differences in the submitted gene models, paying special attention to the three most common errors that might invalidate a model (described below), and investigate any large-scale anomalies (e.g., proposed novel isoform, missing specific exons or isoforms). It is relatively common for one student to make a single error but less common for another student, working independently of the first, to make the same error.

#### Reconciliation Process

Reconciliation is performed using Apollo (Dunn *et al.* 2019), a web-based collaborative gene annotation editor that allows reconcilers to view student-generated models alongside the evidence tracks (Figure 5) used for annotating those models.

**Figure 5:**
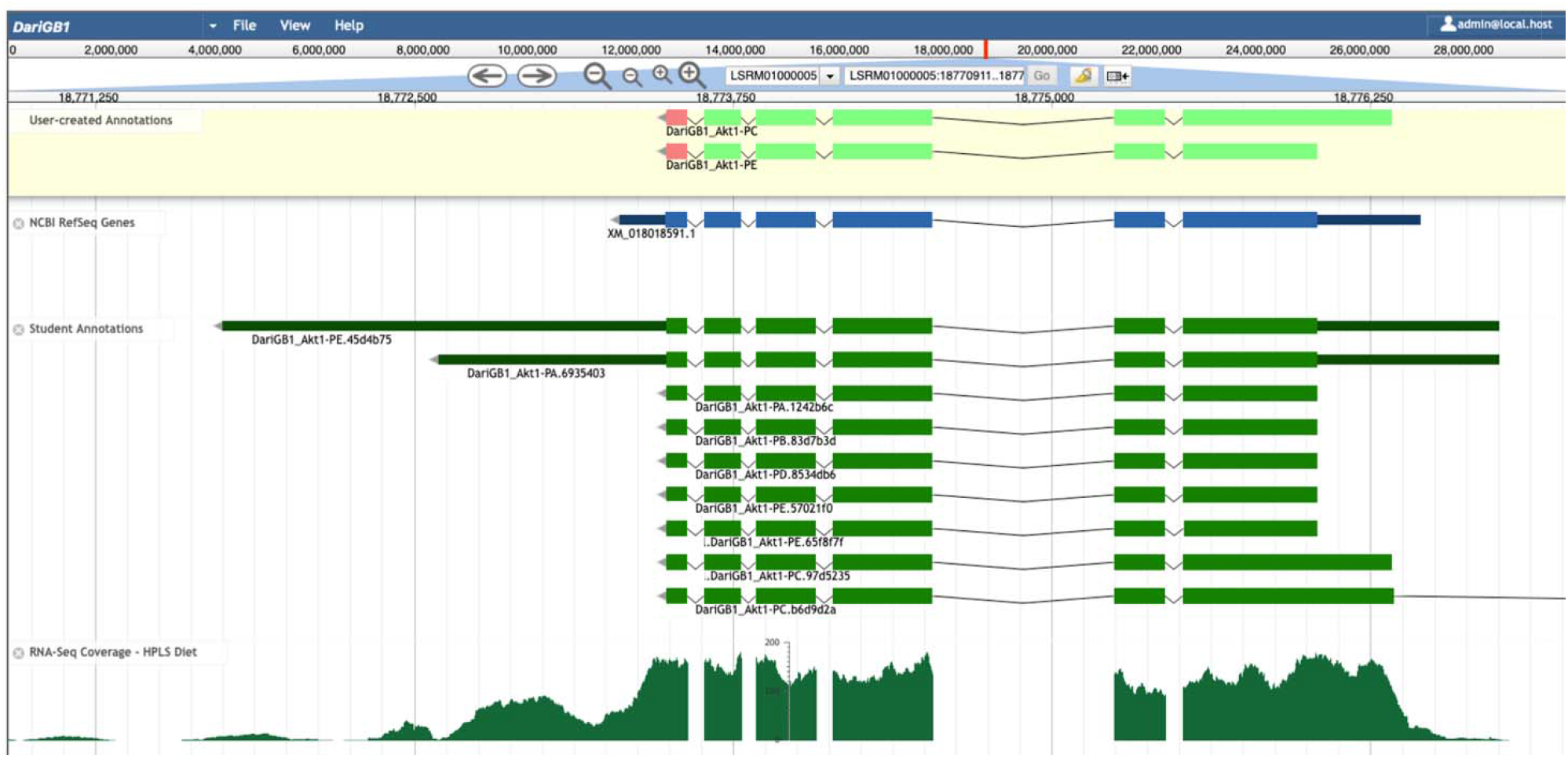
Apollo Screenshot for the Akt1 gene model in D. arizonae. Final gene model for *Akt1* in *D. arizonae*, along with the submitted student models, and RNA-Seq data aligning to the region. The final model shows that despite there being only a single isoform prediction for a protein coding gene by RefSeq, there are likely two protein coding isoforms for this gene, which were annotated using multiple lines of evidence. The second isoform has a larger coding region in the reconciled gene model that is missed by the RefSeq genome predictor.

Reconcilers evaluate the available student annotations for each isoform in conjunction with the other evidence tracks (e.g., sequence similarity, RNA-Seq data, gene predictions) to construct the final gene model that is best supported by the available evidence. After creating a reconciled model, reconcilers draft a *microPublication* describing the supporting evidence for the final gene model (Raciti *et al.* 2018). The student annotators and their faculty mentors then review and approve the article draft. Reconciled models are also used in downstream meta-analysis and deposited into the NCBI Third Party Annotation (TPA) database.

#### Most Common Annotation Errors

Reconciliation consists of reviewing two or more models created by student annotators. The main advantage of manually curated gene models relative to computational predictions is the ability for the curator (in our case, the student annotator) to evaluate and integrate across non-conforming pieces of evidence. The reconciler also pays close attention to each of the putative idiosyncrasies in the proposed model. Some of the most common idiosyncrasies/errors are listed below.

##### Selection of Incorrect Splice Sites

The use of a non-canonical splice site in the orthologous intron in other *Drosophila* species provides supporting evidence for the use of the non-canonical splice site in the target species (Sheth *et al.* 2006; Parada *et al.* 2014; Sibley *et al.* 2016). There are cases where annotators place too high of a priority on the use of canonical GT and AG donor and acceptor splice sites, when there is strong evidence from spliced RNA-Seq reads and conservation in other *Drosophila* species that support the use of non-canonical splice sites.

##### Missing/Extra Exons

There may be cases where the student model is missing exons or has predicted an additional exon relative to the *D. melanogaster* ortholog. Sometimes these predictions are well supported by the data, but often the mismatch in exon number is due to the student annotator failing to fully account for all lines of evidence.

##### Incorrect Ortholog Assignment

Correctly identifying an ortholog depends on an appropriate BLAST search, which provides matches to the gene of interest, and properly evaluating the conservation of synteny. If a student annotates a gene that is not the ortholog, a reconciler must evaluate based on BLAST and synteny what the correct position of the ortholog is in the target species, and if the particular model does not have two congruent annotations, it must be returned to the claim list for annotation by other students.

## Discussion

This annotation protocol has been used by GEP faculty to engage undergraduate students in the comparative annotation of insulin signaling pathway genes in 27 *Drosophila* species. Similar protocols have been used by the GEP on other scientific projects in the past (Slawson *et al.* 2006; Leung *et al.* 2010, 2015, 2017). Collectively, in the last two years, GEP faculty and students across the country have helped us generate 1114 gene models (783 unique models) for the Insulin Pathways project as of December 2020.

To ensure that the gene models produced by GEP students are of high quality, each gene is annotated by at least two students working independently and then reconciled by a more experienced student. This manual curation process ensures the availability of an accurate set of gene models from multiple *Drosophila* species for the comparative study of network architecture and the evolution of genes involved in signaling pathways (e.g., the insulin signaling pathway).

For example, GEP students were able to annotate a new isoform for *Akt1* in *D. arizonae*. This isoform was missed by the NCBI RefSeq gene predictor due to a non-canonical start codon that is conserved in *Akt1* in *D. melanogaster*.

The annotation and reconciliation protocols described in this article have been adapted to two additional scientific projects within the GEP: investigation of the expansion of the Muller F element in four *Drosophila* species and investigation of the evolution of venom proteins in four parasitoid wasp species. The annotation protocol described here is focused only on the annotation coding regions. Once additional experimental data (e.g., CAGE, RAMPAGE, long-read RNA-Seq) becomes available in more species, future analyses will focus on the annotation of untranslated regions and transcription start sites. Faculty can also use this protocol to create new Course-based Undergraduate Research Experiences (CUREs) that engage students in the comparative analysis of genes involved in other metabolic and signaling pathways.

## Data and Software Availability

### Data

The accession number of the assemblies used can be found in Table 1. More information about the tracks used to construct each of the tracks can be found in the track information page for each assembly from the browser of the assembly.

### Software

All custom software and tools generated by the GEP can be accessed from the GEP website (thegep.org). The tools are all web-based applications, thus requiring no special installation on the part of the user. The versions of the tools used can be found in Table 3.

**Table 3:**
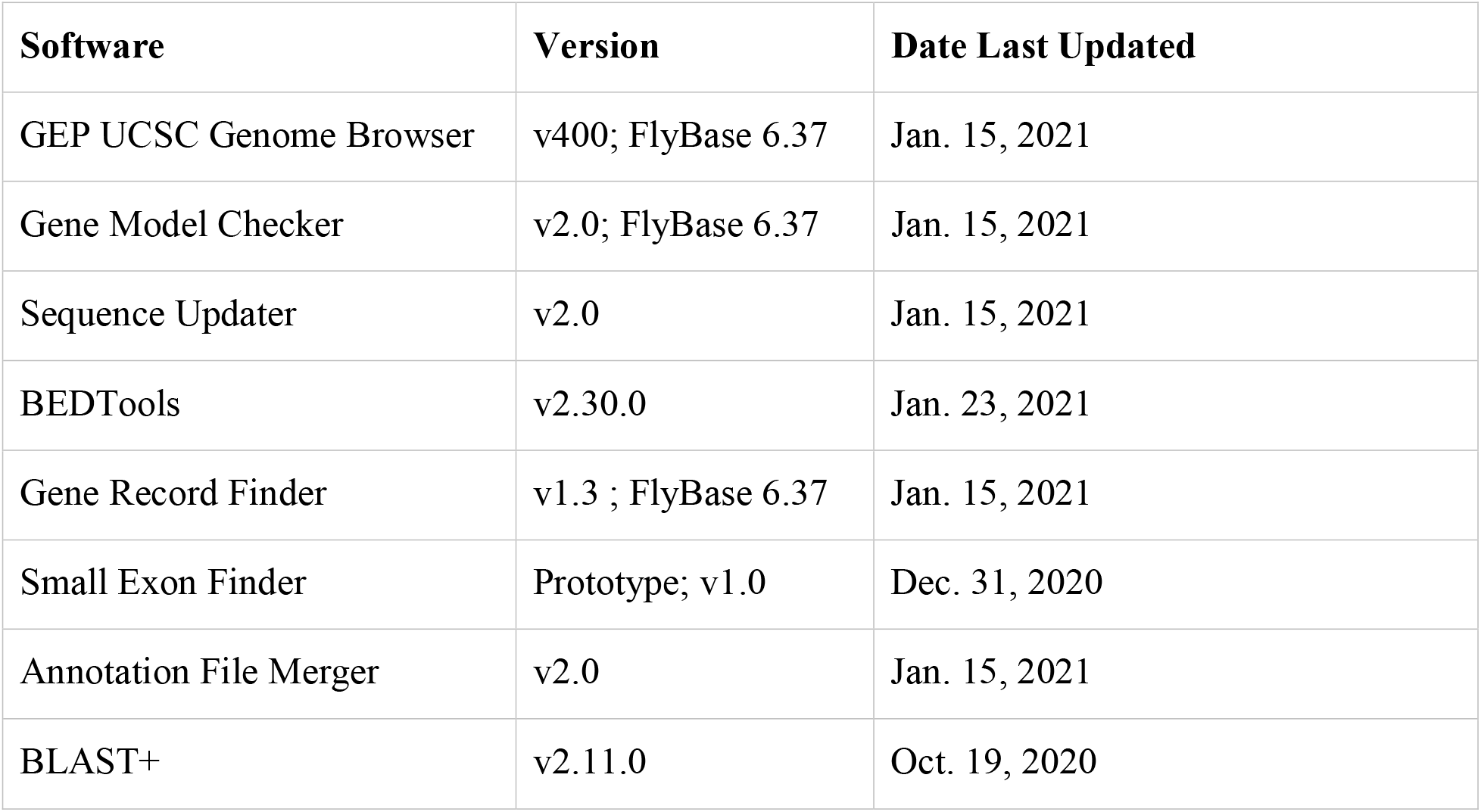
Software Version Information. Version information of the software used for manual gene annotation along with the release date

| Software | Version | Date Last Updated |
| --- | --- | --- |
| GEP UCSC Genome Browser | v400; FlyBase 6.37 | Jan. 15, 2021 |
| Gene Model Checker | v2.0; FlyBase 6.37 | Jan. 15, 2021 |
| Sequence Updater | v2.0 | Jan. 15, 2021 |
| BEDTools | v2.30.0 | Jan. 23, 2021 |
| Gene Record Finder | v1.3 ; FlyBase 6.37 | Jan. 15, 2021 |
| Small Exon Finder | Prototype; v1.0 | Dec. 31, 2020 |
| Annotation File Merger | v2.0 | Jan. 15, 2021 |
| BLAST+ | v2.11.0 | Oct. 19, 2020 |

#### GEP UCSC Genome Browser

Since GEP materials are catered towards undergraduates that may have limited or no experience with programming and command-line interfaces, we endeavor to make all data easily accessible, thereby reducing the barrier to engagement in genomics research. The GEP maintains a mirror of the UCSC Genome Browser with *Drosophila* genomes (https://gander.wustl.edu). We created a these genome browsers (Shaffer *et al.* 2014) with multiple evidence tracks using data generated by various algorithms (Supplement 2).

#### Gene Record Finder

The *Gene Record Finder* summarizes the FlyBase annotations for protein-coding genes in *D. melanogaster*. It provides information about the structure of each *D. melanogaster* gene, such as the number of isoforms (and number of isoforms with unique coding regions), as well as the amino acid sequence for each coding exon and the nucleotide sequence for each transcribed exon. The *Gene Record Finder* also provides exon usage maps that demarcate the exons used by each isoform. This information is used in step two of the annotation protocol (“Identify the Approximate Coordinates of Each Coding Exon”). While this information can be obtained from FlyBase directly, it is easier for annotators to retrieve the gene structure information from a single page instead of through multiple FlyBase Transcript and Polypeptide Reports.

#### Gene Model Checker

The *Gene Model Checker* provides a way for annotators to check their own work when constructing gene models. It verifies that the proposed gene model satisfies basic biological constraints (e.g., maintains an open reading frame, uses canonical start and stop codons, and canonical splice donor and acceptor sites). It also indicates whether the predicted exon number matches that in the orthologous isoform in *D. melanogaster.* It does not inform us whether our proposed model is otherwise accurate. If there is empirical evidence indicating the gene has unusual characteristics (e.g., the use of a non-canonical start codon, stop codon read-through, polycistronic transcripts (Crosby *et al.* 2015)), we provide the supporting evidence for our claims in the annotation report form. The “Dot Plot” section of the *Gene Model Checker* output compares the proposed gene model against the putative ortholog in *D. melanogaster* using protein alignment algorithms. Large gaps in the dot plot or protein alignment might indicate the selection of an incorrect splice site, missing exons, or extra exons in the proposed gene model. Note that this tool does not compare the proposed gene model against other lines of evidence, such as RNA-Seq data or computational gene predictions. The *Gene Model Checker* also produces the three annotation files for the proposed gene model that are required for project submission (i.e., the GFF and FASTA files for the transcript and protein sequences (FAA and FNA)).

The *Gene Model Checker* requires input of the species, assembly, and the scaffold where the putative ortholog is located. The gene model specific information requested by the *Gene Model Checker* includes the name of the ortholog in *D. melanogaster,* the set of coding exon coordinates for the gene in the target species, and the orientation of the gene. Further model specific information requested includes noting whether or not the untranslated regions (UTRs) are included, the gene model is complete (i.e., encompasses all CDSs/UTRs), and whether the genomic region containing the gene in the target species has consensus errors (Figure 6).

**Figure 6:**
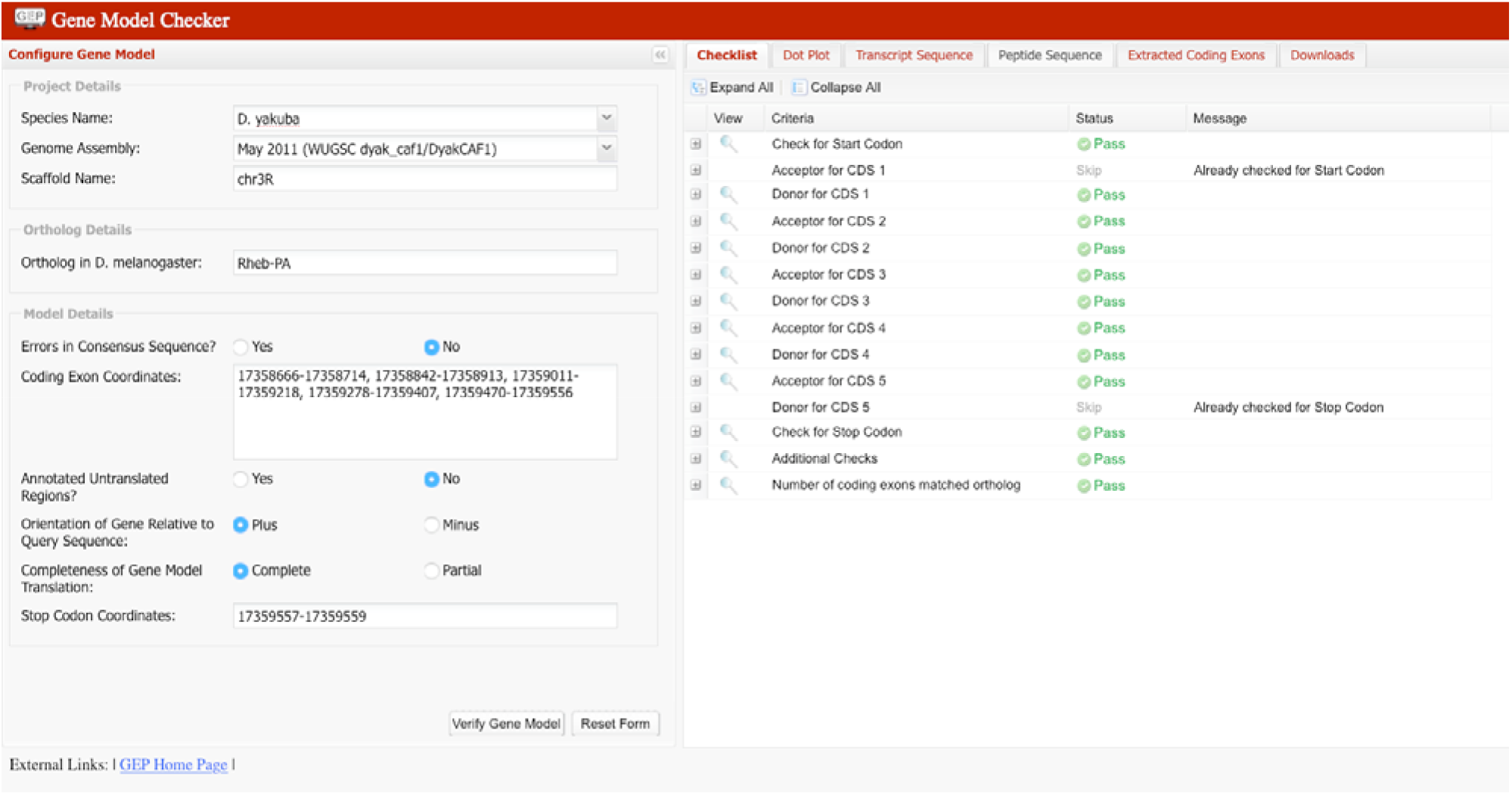
Gene Model Checker. Gene Model Checker with information for *Rheb*-PA in *D. yakuba*: visualizing how we establish biological sanity for one of the two proposed isoform models.

Please note that, at present, this tool only supports the 28 *Drosophila* species that are currently in the GEP UCSC Genome Browser.

#### Small Exons Finder

The typical use case for the *Small Exons Finder* tool is for identifying CDSs that are too small or too weakly conserved to be detected by BLAST. The tool is designed to look for open reading frames (ORFs) that satisfy a set of biological constraints. These constraints include the type of CDS (i.e., initial, internal, or terminal CDS), the phase of the donor or acceptor site, and the expected CDS size according to the *D. melanogaster* model. The *Small Exons Finder* then looks for ORFs in the provided sequence that conform to the aforementioned constraints. Compared to the *ORFfinder* tool developed by NCBI (Rombel *et al.* 2002), the *Small Exons Finder* allows users to search for open reading frames that are less than 30bp in size, and it can search for initial, internal, and terminal coding exons with constraints on the phases of the donor and acceptor sites.

#### Sequence Updater

The *Sequence Updater* tool is primarily used to create a VCF (Danecek *et al.* 2011) file to correct assembly errors in the consensus sequence. It provides an updated assembly sequence with the base(s) hypothesized to be more accurate within a specific region in the assembly. This tool is used to correct errors (i.e., base substitutions, insertions, deletions) in the genomic sequence of the project region. The VCF file produced by the *Sequence Updater* can be used with the *Gene Model Checker* to validate a gene model with consensus errors.

#### Annotation Files Merger

The *Gene Model Checker* produces GFF, FNA, and FAA files for each isoform. The submission pipeline requires a single GFF, FNA, and FAA file that includes all of the isoforms in a project. For each type of annotation file, the *Annotation Files Merger* is used to combine the annotations for all of the isoforms in a project into a single file. The *Annotation Files Merger* also enables the user to view the combined GFF file as a custom track on the GEP UCSC Genome Browser.

#### BEDTools

BEDTools (Quinlan and Hall 2010) was used to perform genomic arithmetic and compare locations of genomic features in multiple tracks of the GEP UCSC Genome Browser.

#### BLAST+

The entire BLAST suite (Camacho et al. 2009) was used for multiple aspects of the protocol, including, but not limited to sequence annotation and synteny assignment.

## Acknowledgements

The work done by GEP Faculty and their students has generated student models that have benefited us in refining our protocol. We would also like to thank the reconcilers who have generated models that can be used for downstream analysis.

