## Supplementary figures and images for "Manual Annotation of Genes within *Drosophila* Species: the Genomics Education Partnership protocol"

### Supplement 1

#### Pathways Annotation Workflow

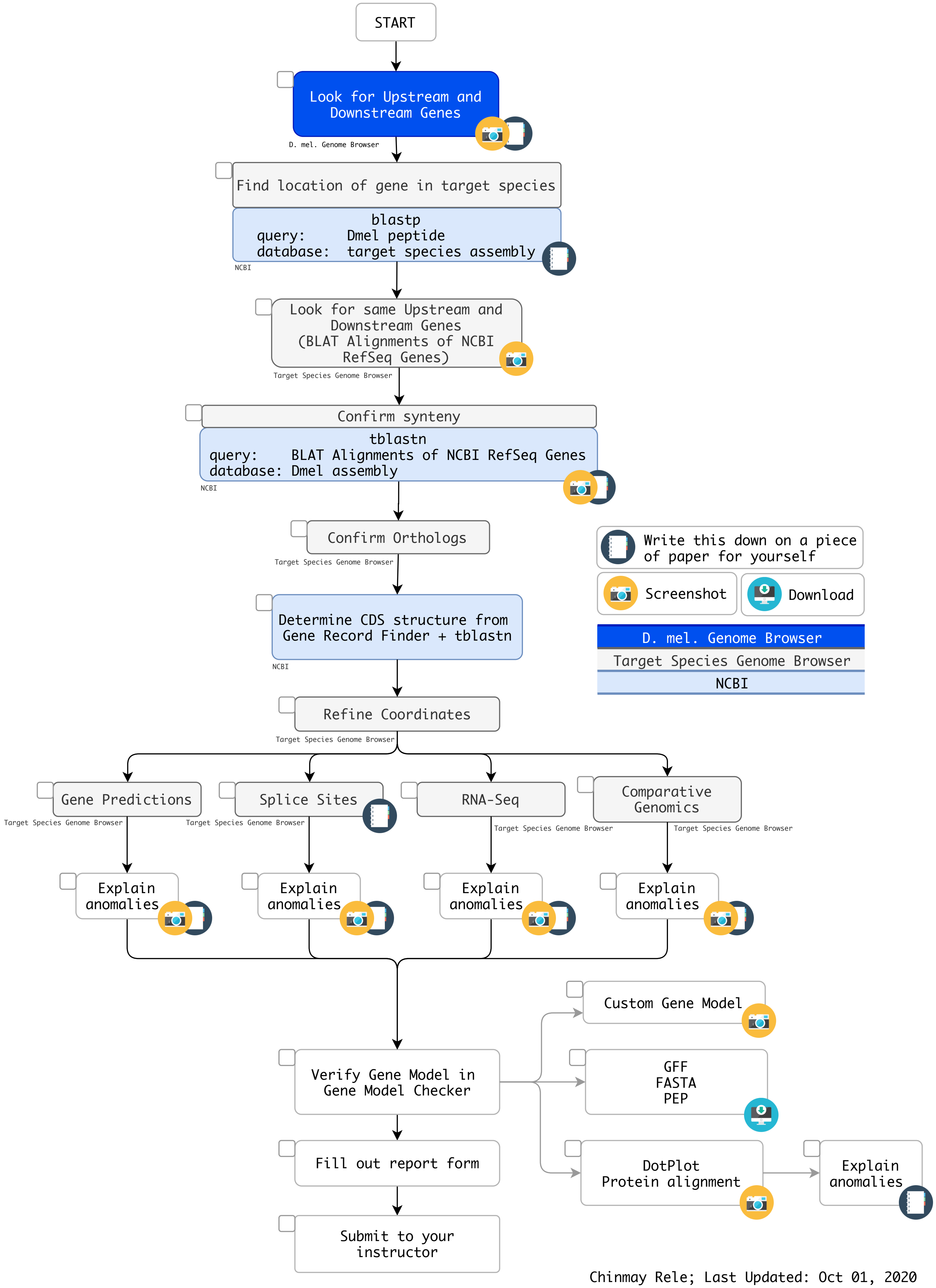


Chinmay Rele; Last Updated: Oct 01, 2020

### Supplement 5

#### Determining CDS Coordinates

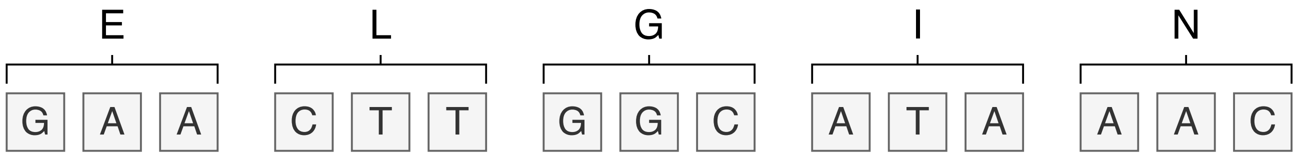


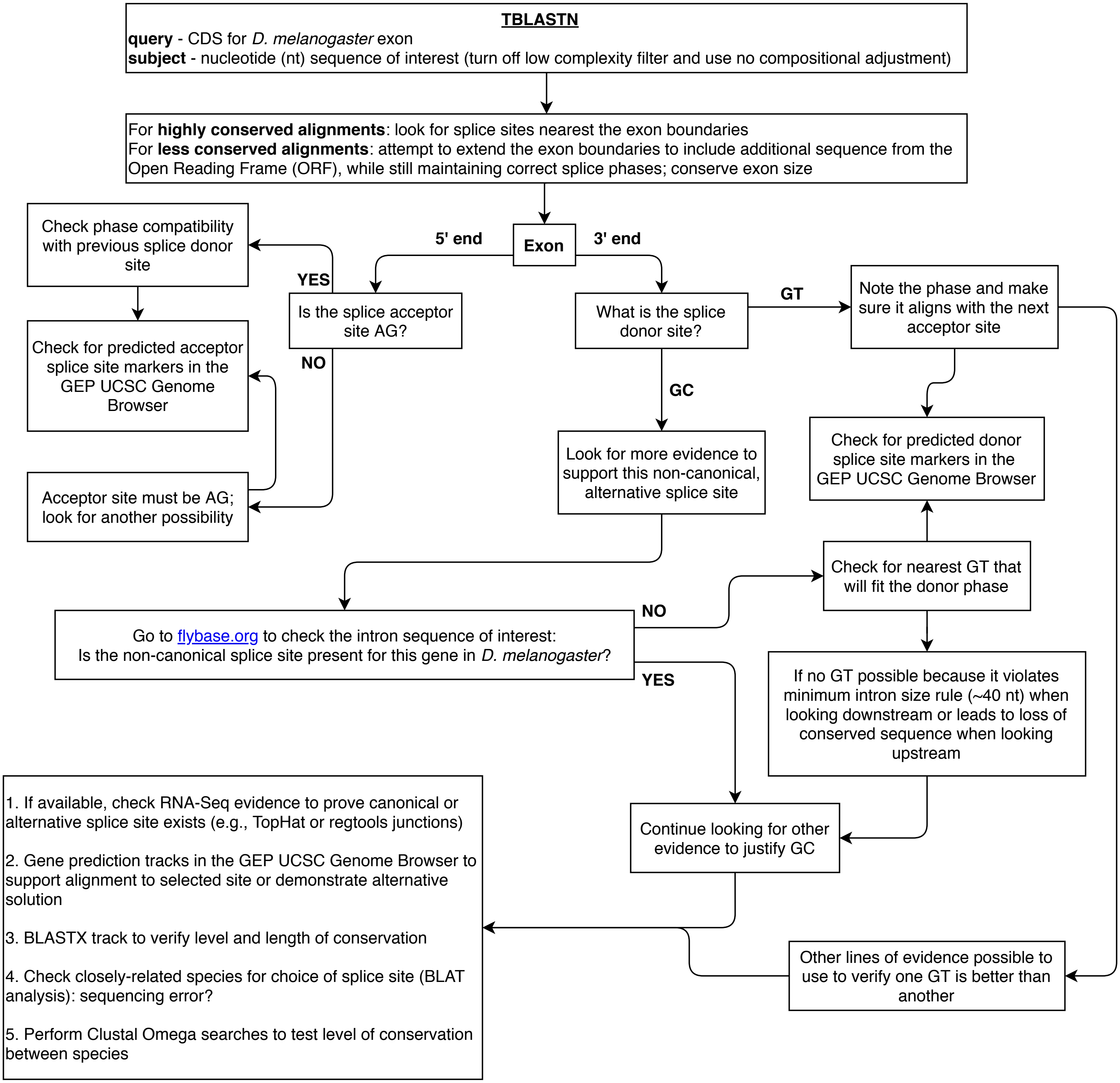


Adapted from Jeannette Wong 08/05/2010
Last Update: 08/24/2020
