## Supplement 2 for "Manual Annotation of Genes within *Drosophila* Species: the Genomics Education Partnership protocol"

### Pathways Project: Annotation Walkthrough

#### Prerequisites

- Understanding Eukaryotic Genes (modules 1, 4, 5, and 6)
- RNA-Seq Primer (automatic download)
- An Introduction to NCBI BLAST

#### Resources & Tools

- All links for this walkthrough can be found on the Pathways Project page of the GEP website (thegep.org/pathways).

### Introduction

The Pathways Project is focused on annotating genes found in well characterized signaling and metabolic pathways across the *Drosophila* genus. The current focus is on the insulin signaling pathway which is well conserved across animals and critical to growth and metabolic homeostasis. The long-term goal of the Pathways Project is to analyze how the regulatory regions of genes evolve in the context of their positions within a network.

This walkthrough illustrates how to apply the GEP annotation strategy for the Pathways Project to construct a gene model for the Ras homolog enriched in brain (*Rheb*) gene in *Drosophila yakuba*. This walkthrough focuses on annotation of the coding regions so we will not annotate the untranslated regions (UTRs) or the transcription start site (TSS).

It is important to note that the *Rheb* gene in *D. yakuba* is relatively straightforward in comparison to what your own project might be. Some of the steps in this walkthrough might appear to be excessive, but keep in mind that you will potentially have a more complex project, so it is important to follow this protocol as we have tried to equip you with most of what you will need should you encounter complexities.

We recommend you follow the parts in the order they are presented, and then refer back to particular parts when needed for your own project. Note that the figures have been configured to fit this document while still maintaining readability; therefore, your screen may differ slightly. Commas have been included with all coordinates to improve readability, but you do not have to enter them in the Genome Browser (navigation will work the same with or without commas).

| | | | |
|---|---|---|---|
| FYI | (For Your Information): further details and explanations or useful tips | | Reminder |
| | Review of content listed in "Prerequisites" | | Caution |

**Throughout this walkthrough, text boxes with icons are used to assist you.**

### Part 1: Examine the genomic neighborhood surrounding the gene of interest in *D. melanogaster*

1. Navigate to the GEP UCSC Genome Browser Gateway Page.
2. Click on "D. melanogaster" in the "REPRESENTED SPECIES" table.
3. Ensure "Aug. 2014 (BDGP Release 6 + ISO1 MT/dm6)" under the "D. melanogaster Assembly" field is selected.
4. Enter "**Rheb**" under the "Position/Search Term" field.
5. Click on the "Go" button (Figure 1).

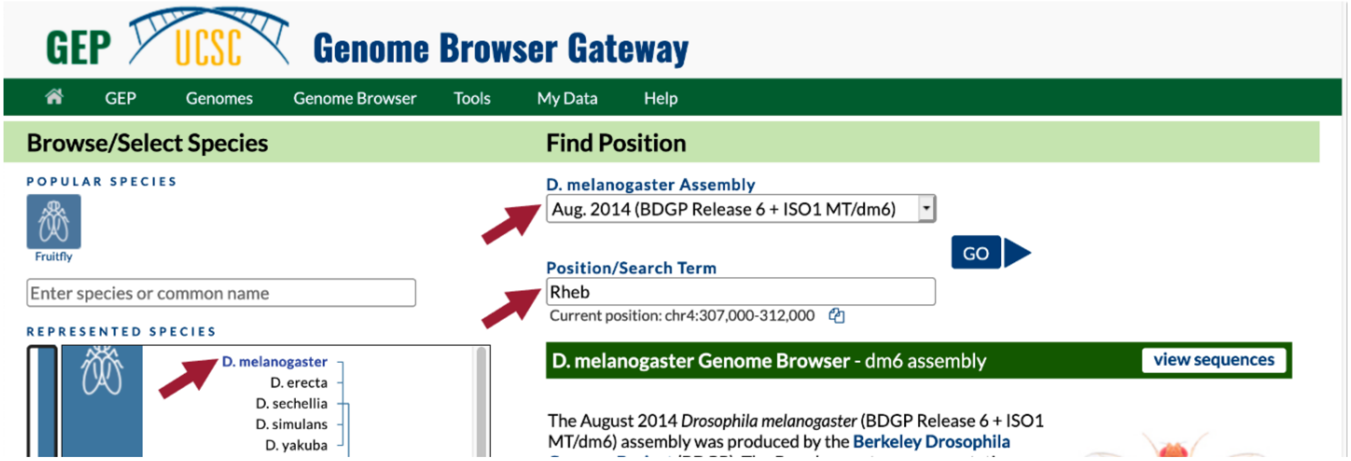

**Figure 1 Navigate to the *Rheb* gene in the *D. melanogaster* Aug. 2014 (BDGP Release 6 + ISO1 MT/dm6) assembly.**

6. On the following page, "FlyBase Protein-Coding Genes," click on "Rheb-RA at chr3R:5568921-5570491" (Figure 2).

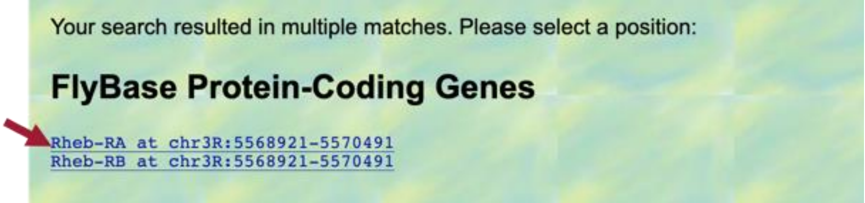

**Figure 2 Navigate to *Rheb-RA* on the right arm of chromosome 3 (chr3R) in *D. melanogaster*.**

FYI

As shown in Figure 2, *Rheb* has two isoforms in *D. melanogaster*, and they both span the same coordinates (i.e., both Rheb-RA and Rheb-RB are located on the scaffold of the right arm of chromosome 3 (chr3R) at 5,568,921 – 5,570,491).

Therefore, it doesn't matter which isoform we selected here, but **if the coordinates of the isoforms in your own project are different, you should choose the longest isoform** for this step.

7. Because the Genome Browser remembers our previous track display settings, click on the “default tracks” button in the list of buttons below the Genome Browser image (Figure 3).

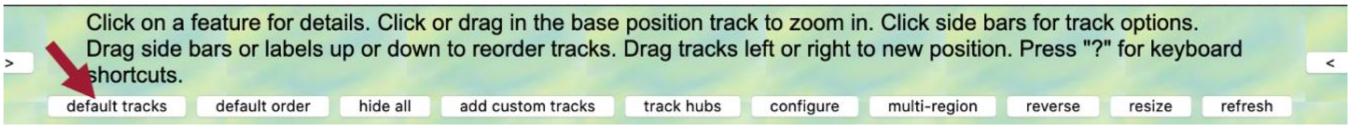

**Figure 3 Click on the “default tracks” button.**

In the “FlyBase Protein-Coding Genes” track, we should now see the gene structure for the two isoforms of the *Rheb* gene (i.e., Rheb-RA and Rheb-RB). Notice that *Rheb* is on the “chr3R” scaffold (Figure 4).

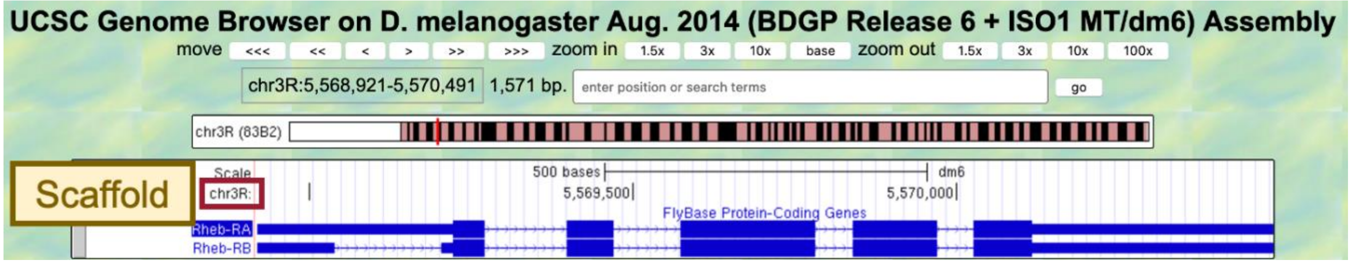

**Figure 4 There are two isoforms of *Rheb* in *D. melanogaster* (Rheb-RA and Rheb-RB). *Rheb* is located on the “chr3R” scaffold of the *D. melanogaster* genome assembly.**

8. Zoom out until you can see two genes on either side of *Rheb*.

We are now viewing the genomic neighborhood of the *Rheb* gene in *D. melanogaster* (i.e., region of the chr3R scaffold containing the *Rheb* gene and two adjacent upstream and downstream genes) (Figure 5).

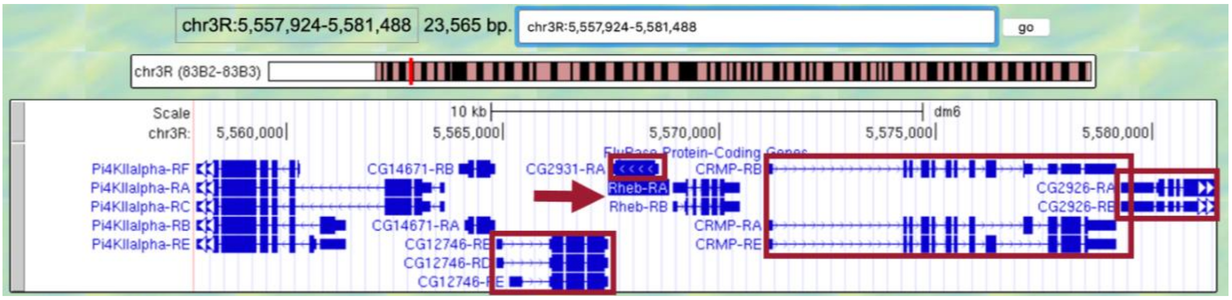

**Figure 5 The genomic neighborhood of *Rheb* (arrow) includes *CG12746, CG2931, CRMP*, and *CG2926*.**

FYI

- **Upstream**: located on the 5’ side of the reference (target) gene
- **Downstream**: located on the 3’ side of the reference (target) gene

As we learned in Understanding Eukaryotic Genes: Module 1, some genes on the same DNA molecule may be transcribed in opposite directions (i.e., genes on the positive (top) strand of the DNA are transcribed from left to right and genes on the negative (bottom) strand of the DNA are transcribed from right to left). Since genes have directionality, the upstream and downstream areas of the DNA may change depending on which gene is used as the reference. In this walkthrough, our reference (target) gene is *Rheb*.

Now we need to draw a sketch of the genomic neighborhood of *Rheb* (Figure 6). In order to do so, we must identify the direction of transcription for each gene by zooming in on an intron of each gene.

9. Start by zooming in on an intron of *Rheb*.
10. Since the arrows within the introns of *Rheb* point to the right, in our sketch we need to draw an arrow pointing to the right and label it *Rheb*.
11. Repeat Steps 9-10 for the two nearest genes on either side of *Rheb*.
    - Note: If a gene only has one exon, and therefore no introns, zoom into the exon to identify the direction of transcription.

The arrows within the introns of *Rheb* point to the right; therefore, we know *Rheb* is transcribed from left to right and located on the positive (top) strand of DNA. Since *Rheb* is our reference (target) gene (i.e., gene we are annotating), the two nearest genes on its 5' side are considered upstream and the two nearest genes on its 3' side are considered downstream.

- The two nearest upstream genes to *Rheb* are *CG2931* and *CG12746*.
- The two nearest downstream genes to *Rheb* are *CRMP* and *CG2926*.

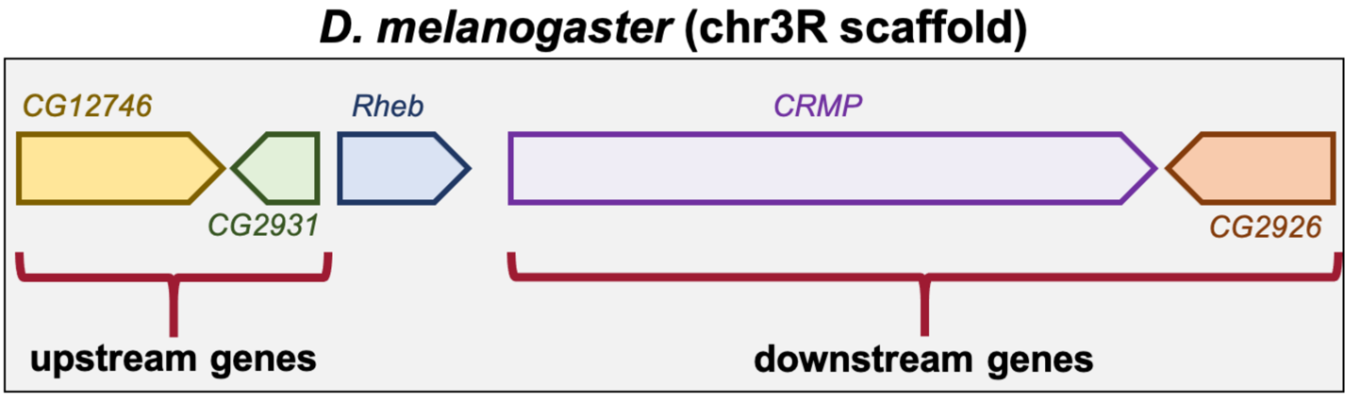

**Figure 6 Sketch of the genomic neighborhood of *Rheb* in *D. melanogaster*. Since the direction of the arrows within their introns point to the right, *CG12746* and *CRMP* are on the positive strand. *CG2931* and *CG2926* are on the negative strand since the direction of the arrows within their introns point to the left.**

FYI

Each protein-coding gene annotated by FlyBase in *D. melanogaster* has an annotation symbol that begins with the prefix "**CG**" (i.e., Computed Gene). Unless genes are characterized experimentally and formally named, they are referred to by this symbol. For example, the two neighboring upstream genes (i.e., *CG2931* and *CG12746*) and one downstream gene (i.e., *CG2926*) have not yet been named. However, the Ras homolog enriched in brain gene has been experimentally characterized and so it is referred to by its gene symbol *Rheb*.

### Part 2: Identify the genomic location of the ortholog in *D. yakuba*

Now that we've examined the genomic neighborhood of *Rheb* in *D. melanogaster*, we need to identify the location of *Rheb* in *D. yakuba*.

#### Part 2.1: Retrieve the protein sequence for Rheb-PA in *D. melanogaster*

1. In the "FlyBase Protein-Coding Genes" track, click on "Rheb-RA" (Figure 7).

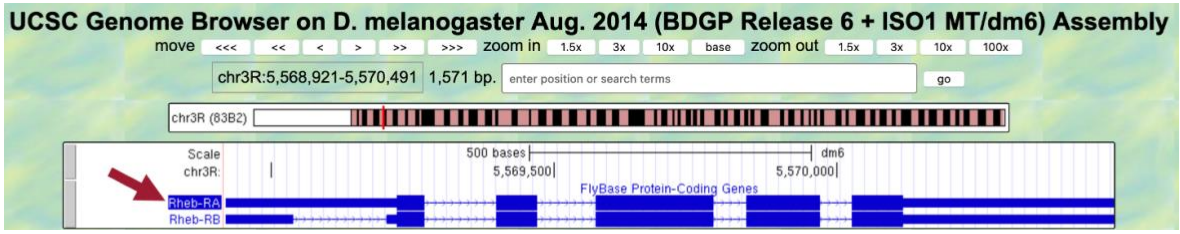

**Figure 7 Click on "Rheb-RA" to view details regarding this protein-coding gene that was annotated by FlyBase.**

2. Under the "Links to sequence" heading, click on the "Translated Protein" link (Figure 8, left).

We are now viewing the sequence for the 182 amino acids in the translated protein of Rheb-RA in *D. melanogaster* (Figure 8, right).

3. Copy the entire sequence (including the header) so we can use it in our *tblastn* search.

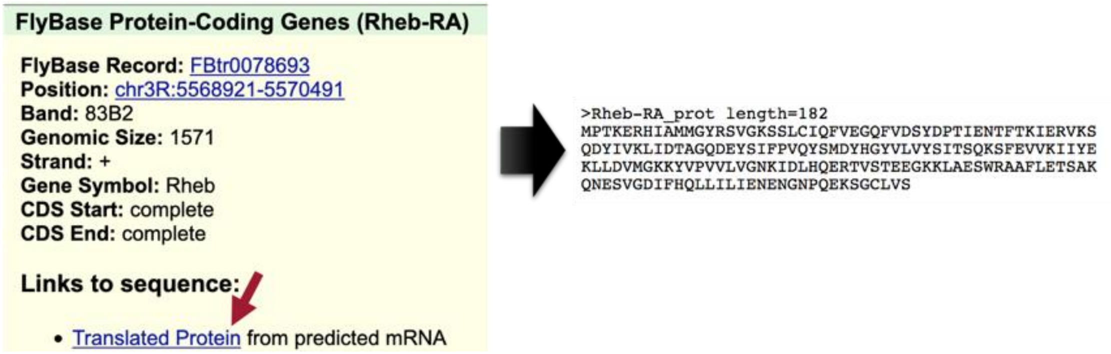

**Figure 8 Click on the "Translated Protein" link for the Rheb-RA feature to obtain the sequence of the translated protein.**

FYI

What is the difference between Rheb-**RA** and Rheb-**PA**?

The prefix "Rheb" corresponds to the gene symbol. The "R" or "P" in the suffix designates the associated transcript (mRNA) or protein-product of the gene, respectively. The "A" in the suffix refers to the A isoform of the gene.

Gene symbols (e.g., *Rheb*) are italicized while their mRNA and protein products are not (e.g., Rheb-RA and Rheb-PA).

#### Part 2.2: Perform a *BLAST* search of Rheb-PA against the *D. yakuba* genome assembly

The Basic Local Alignment Search Tool (*BLAST*) finds regions of local similarity between biological sequences by comparing nucleotide or protein sequences to sequence databases (or to an individual nucleotide or protein sequence) and calculates the statistical significance of each match. The statistical values we will focus on are:

- **Expect Value (E-value)**: describes the number of hits one can "expect" to see by chance when searching a database of a particular size
  - The lower the E-value (i.e., the closer it is to zero), the more significant the match.
- **Percent Identity**: describes how similar the query sequence is to the database (or subject) sequence (i.e., how many characters in each sequence are identical)
  - The higher the Percent Identity (i.e., the closer it is to 100), the more significant the match.

The five traditional *BLAST* programs:

- *blastn* program searches nucleotide databases using a nucleotide query
- *blastp* program searches protein databases using a protein query
- *blastx* program searches protein databases using a translated nucleotide query
- *tblastn* program searches translated nucleotide databases using a protein query
- *tblastx* program searches translated nucleotide databases using a translated nucleotide query

| BLAST Program | Query (sequence to match) | Database/Subject (searching for match) | Function | Common Use Cases |
|---|---|---|---|---|
| blastn (nucleotide BLAST) | nucleotide | nucleotide | searching with shorter queries, cross-species comparison | map mRNAs against genomic assemblies |
| blastp (protein BLAST) | protein | protein | general sequence identification and similarity searches | search for proteins similar to predicted genes |
| blastx | nucleotide → protein | protein | identifying potential protein products encoded by a nucleotide query | map proteins/CDS against genomic sequence |
| tblastn | protein | nucleotide → protein | identifying database sequences encoding proteins similar to query | map proteins against genomic assemblies |
| tblastx | nucleotide → protein | nucleotide → protein | identifying nucleotide sequences similar to the query based on their coding potential | identify genes in unannotated sequences |

Arrows indicate the *BLAST* program translates the nucleotide sequence before performing the search.

**See An Introduction to NCBI BLAST for additional details.**

In Part 2.1, we retrieved the protein sequence for Rheb-PA in *D. melanogaster*. Now we need to perform a *BLAST* search of Rheb-PA against the *D. yakuba* genome assembly.

Since we are looking for the orthologous region of *Rheb* in *D. yakuba*, we want *BLAST* to search the entire genome of *D. yakuba* to identify regions of local similarity with the protein sequence of *Rheb* in *D. melanogaster* we obtained in Part 2.1. In other words, we need to *BLAST* our *Rheb* protein sequence from *D. melanogaster* against the entire genome of *D. yakuba* to narrow down the possible regions where *Rheb* could be located in *D. yakuba*.

For this *BLAST* search, we will use the *tblastn* program to search the translated nucleotide database of our target species, *D. yakuba*, using the protein sequence of *Rheb* in *D. melanogaster* as our query.

- **Query**: sequence we are looking to match (protein sequence of *Rheb* in *D. melanogaster*)
- **Database (Subject)**: collection of sequences we are searching for matches (*BLAST* will translate the entire genome of *D. yakuba* before searching for a match to the Rheb-PA sequence from *D. melanogaster*)

1. Navigate to the Pathways Project Genome Assemblies page.
2. Click on the "Genome *BLAST*" link for *D. yakuba* (Figure 9).
   - Note: Here we are selecting the *D. yakuba* genome database to search.

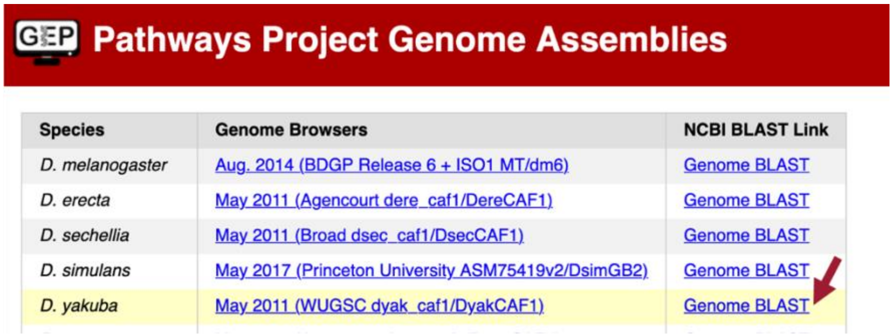

| Species | Genome Browsers | NCBI BLAST Link |
|---|---|---|
| D. melanogaster | Aug. 2014 (BDGP Release 6 + ISO1 MT/dm6) | Genome BLAST |
| D. erecta | May 2011 (Agencourt dere_caf1/DereCAF1) | Genome BLAST |
| D. sechellia | May 2011 (Broad dsec_caf1/DsecCAF1) | Genome BLAST |
| D. simulans | May 2017 (Princeton University ASM75419v2/DsimGB2) | Genome BLAST |
| D. yakuba | May 2011 (WUGSC dyak_caf1/DyakCAF1) | Genome BLAST |

**Figure 9 Click on the "Genome *BLAST*" link for *D. yakuba* in the "NCBI *BLAST* Link" column.**

3. Make sure the "*tblastn*" tab is selected (Figure 10).
4. Paste the Rheb-PA sequence we copied from Part 2.1 into the "Enter Query Sequence" text box.
5. Click within the "Job Title" text box and it should automatically populate.
6. Make sure the check box next to "Show results in a new window" is selected.
7. Click on the "*BLAST*" button.

FYI: The National Center for Biotechnology Information (NCBI) periodically updates the genome assemblies used for BLAST searches that can cause a host of issues when they occur in the middle of a semester. Using the "Genome BLAST" links on the "Pathways Project Genome Assemblies" page ensures student annotators are navigating to the correct genome assembly database when performing their BLAST search against the whole genome of their target species. However, this circumvention will only be needed in Part 2 of the Pathways Project protocol. In Parts 3 and 5, we will navigate directly to the NCBI BLAST home page.

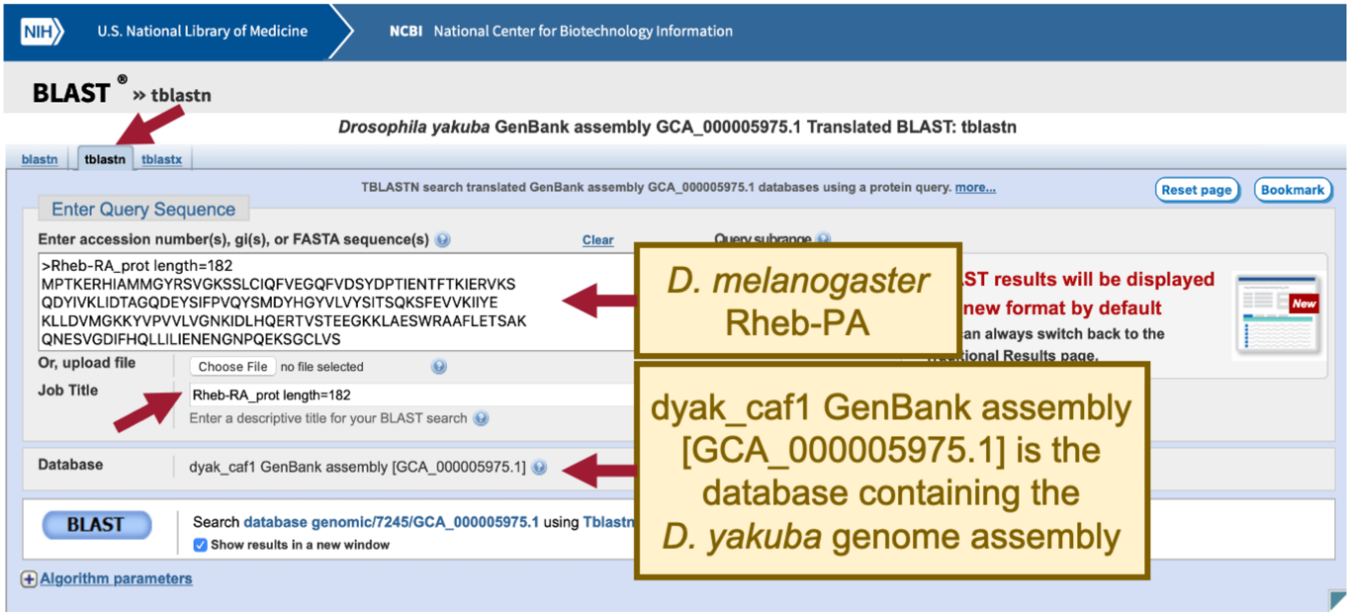

**Figure 10 Configure *tblastn* to compare the *D. melanogaster* protein Rheb-PA (query) against the *D. yakuba* whole genome assembly (database).**

When performing a search, *BLAST* may return any number of matches (often referred to as "hits") for regions of local similarity between our query sequence and database; however, each hit is not necessarily statistically significant. *BLAST* provides statistical scores to help us determine which alignments between the two sequences are statistically significant and which are spurious (i.e., likely occurred by chance alone and, therefore, are not evidence of real biological conservation). If *BLAST* returns multiple good hits (i.e., more than one match with a low E-value and a high sequence identity), we will need to investigate them all further to determine if we have the best ortholog.

Our *tblastn* search found six regions within the *D. yakuba* genome that show similarities with the protein sequence of *Rheb* in *D. melanogaster* (Figure 11); however, only one of these is a good hit (*Drosophila yakuba* strain Tai18E2 chromosome 3R, whole genome shotgun sequence; sequence identity: 97.14% and E-value: 2e-78[1]). The second hit has a much higher E-value (8e-37) and much lower percent identity (43.75%), and this pattern of increasingly lower quality matches continues throughout the other four matches as well. Therefore, we will continue our analysis based on the hypothesis that the putative ortholog of Rheb-PA in *D. yakuba* is located somewhere in the scaffold of chromosome 3R (chr3R).

Notice that each of the genome regions of our six hits has a unique accession number. The accession number for the chromosome 3R scaffold in *D. yakuba* is CM000160.2 (Figure 11).

FYI Similar to how humans have unique fingerprints that can be used to identify them at crime scenes, each biological sequence is assigned a unique accession number to allow scientists to identify it from the millions of other sequences stored in a database.

[1] 2e-78 = 2 x $10^{-78}$

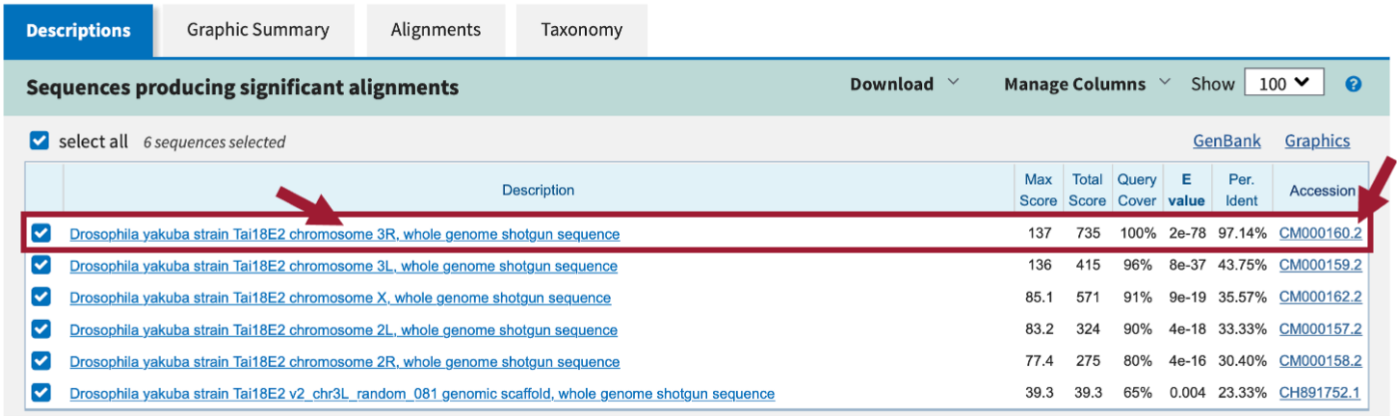

**Figure 11 Our *tblastn* search with the *D. melanogaster* protein Rheb-PA (query) against the *D. yakuba* whole genome assembly (database) found six regions of similarity. The best match is located on the "chromosome 3R" scaffold (accession number: CM000160.2) of *D. yakuba*.**

#### Part 2.3: Summarize the *tblastn* results for Rheb-PA on *D. yakuba* chr3R

1. Under the "Descriptions" tab, uncheck the box next to "select all" (Figure 12).
   - Note: This will hide the alignments from chromosomes 3L, X, 2L, and 2R since we only want to look at the best match (i.e., chromosome 3R).
2. Click on the "*Drosophila yakuba* strain Tai18E2 chromosome 3R, whole genome shotgun sequence" link in the "Description" column to navigate to the alignment.

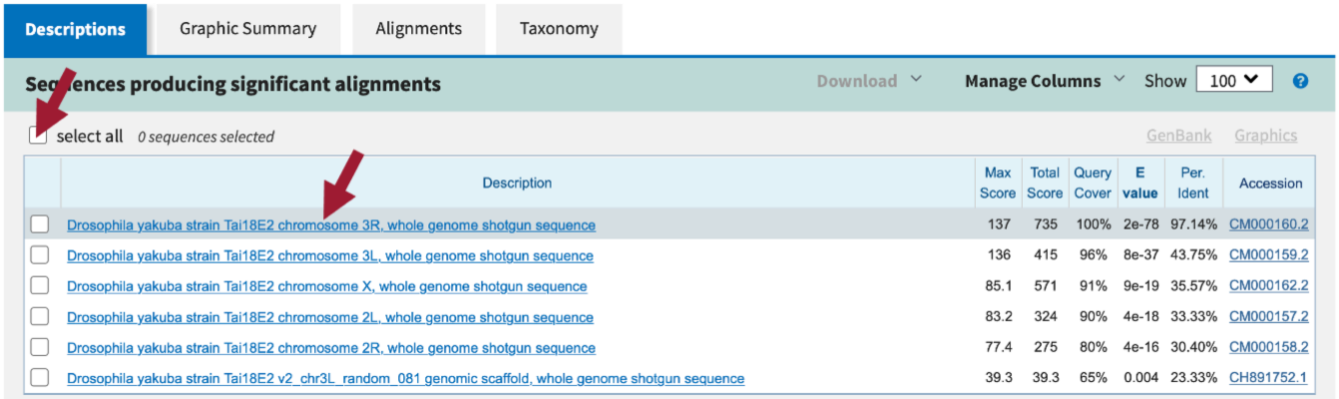

**Figure 12 Uncheck "select all" and then click on the link for the best match to navigate to the alignment.**

3. In the blue toolbar for the *BLAST* hit, select the "Subject start position" option from the drop-down menu of the "Sort by" field to order the matches based on the start coordinates on *D. yakuba* chromosome 3R in ascending order (Figure 13).

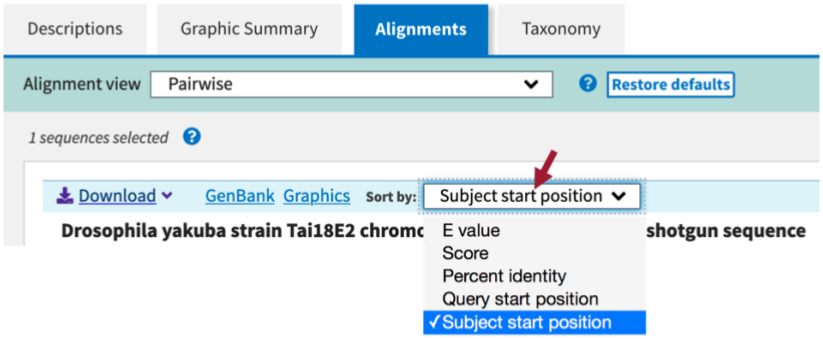

**Figure 13 In the blue toolbar for the *BLAST* hit, select the "Subject start position" option from the drop-down menu of the "Sort by" field to order the matches based on the start coordinates of the *D. yakuba* chromosome 3R scaffold in ascending order.**

We should now see 12 ranges listed in ascending order. Each range corresponds to a contiguous portion of the subject sequence that shows significant sequence similarity (i.e., matches) with a portion of the query sequence. Here, the 12 ranges show matches between Rheb-PA in *D. melanogaster* (Query) and a range of coordinates from chromosome 3R in *D. yakuba* (Sbjct). For example, Range 1 shows a match between Rheb-PA in *D. melanogaster* (55 – 163) and coordinates 5,727,469 – 5,727,792 of chromosome 3R in *D. yakuba* when translated in frame +1. Range 1 has an E-value of 2e-10 and a sequence identity of 36% (Figure 14).

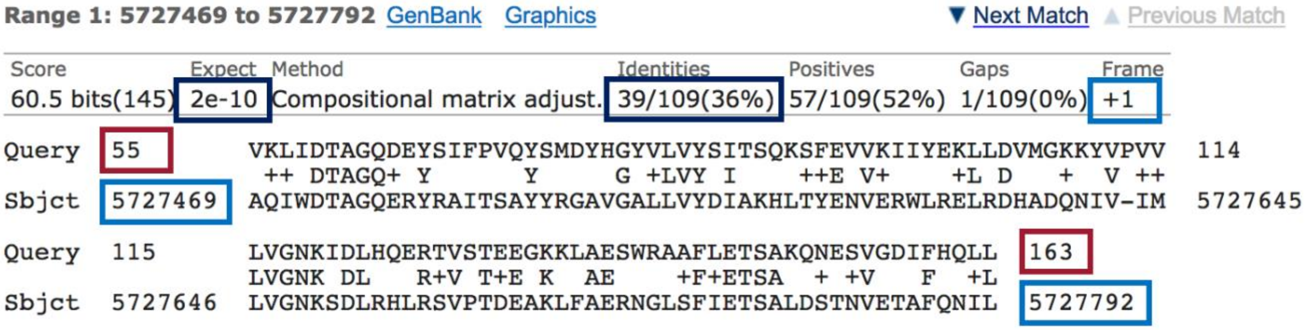

**Figure 14 The query coordinates (55 – 163) and subject coordinates (5,727,469 – 5,727,792) are shown in red and blue, respectively.**

Since chromosome 3R in *D. yakuba* is 28,832,112 bp long, it is likely we will have some ranges (i.e., alignment matches or hits) that do not correspond to our ortholog; therefore, we need to examine each match more closely.

Remember that we are looking for matches with low E-values and high sequence identities, and there are five matches (ranges 7-11) that fit these criteria (E-value of 2e-78 and sequence identities that range from 83% to 97%). These five alignment matches are also collinear and appear on the same strand of DNA (+ Frame) (Figure 15).

The best collinear set of alignments to Rheb-PA is located at 17,358,666 – 17,359,556 on the chromosome 3R scaffold of the *D. yakuba* genome assembly and the five alignment matches cover all 182 amino acids of Rheb-PA (Figure 15). Therefore, we will continue our analysis based on the hypothesis that the putative ortholog of Rheb-PA is located at approximately 17,358,666-17,359,556 on the chromosome 3R scaffold of the *D. yakuba* genome assembly. See Appendix A for details on investigating the other *tblastn* alignments to *D. yakuba* chromosome 3R scaffold.

| Range | *D. melanogaster* | | *D. yakuba* | | E-Value | Identities (%) | Subject Frame |
|---|---|---|---|---|---|---|---|
| | Query Start | Query End | Subject Start | Subject End | | | |
| 1 | 55 | 163 | 5,727,469 | 5,727,792 | 2e-10 | 36 | +1 |
| 2 | 59 | 128 | 5,883,589 | 5,883,383 | 1e-04 | 37 | -3 |
| 3 | 8 | 131 | 9,373,502 | 9,373,062 | 1e-09 | 30 | -2 |
| 4 | 54 | 130 | 9,387,874 | 9,387,623 | 4e-11 | 40 | -3 |
| 5 | 6 | 44 | 9,388,199 | 9,388,062 | 0.002 | 46 | -2 |
| 6 | 18 | 108 | 9,658,240 | 9,658,575 | 5e-06 | 28 | +1 |
| 7 | 1 | 20 | 17,358,666 | 17,358,725 | 2e-78 | 90 | +3 |
| 8 | 16 | 45 | 17,358,838 | 17,358,927 | 2e-78 | 83 | +1 |
| 9 | 40 | 109 | 17,359,007 | 17,359,216 | 2e-78 | 97 | +2 |
| 10 | 111 | 153 | 17,359,279 | 17,359,407 | 2e-78 | 93 | +1 |
| 11 | 153 | 182 | 17,359,467 | 17,359,556 | 2e-78 | 93 | +3 |
| 12 | 53 | 173 | 26,681,803 | 26,681,411 | 1e-05 | 30 | -3 |

best collinear set of alignments to Rheb-PA

**Figure 15 Summary of the *tblastn* search results for the 12 matches to Rheb-PA within chromosome 3R of *D. yakuba*. The best collinear set of alignments to Rheb-PA is located at 17,358,666-17,359,556.**

#### Part 3: Examine the genomic neighborhood of the putative ortholog in *D. yakuba*

In Part 1, we sketched the genomic neighborhood of *Rheb* in *D. melanogaster*. Here we will examine the genomic neighborhood of *Rheb* in *D. yakuba* and then compare the order and orientation of these genes to what we found in *D. melanogaster*.

Based on parsimony, the genes surrounding *Rheb* in *D. yakuba* should be identical or very similar in function to the genes in the genomic neighborhood of *Rheb* in *D. melanogaster*. Additionally, the neighboring genes should also match in orientation (look at the direction of transcription of the neighboring genes; this is called synteny). Since *Rheb* is on the positive strand in *D. melanogaster* (i.e., transcribed left to right) and since *CG2926* is on the negative strand in *D. melanogaster* (i.e., transcribed right to left), these two genes should also be on different strands (i.e., transcribed in different directions) in *D. yakuba*.

FYI

**Navigating the GEP UCSC Genome Browser**

Each annotation track has an associated description page that contains a discussion of the track, the methods used to create the annotation, and, in some cases, filter and configuration options to fine-tune the information displayed in the track (e.g., RNA-Seq tracks).

To view the description page, click on the gray mini-button to the left of a displayed track (left) or on the label for the track in the Track Controls section (right).

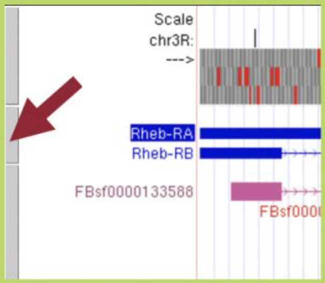

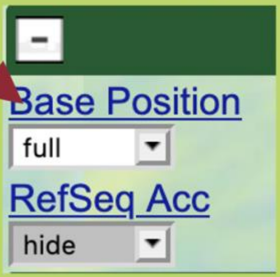

- Click or drag in the base position track to zoom in.
- Drag tracks left or right to a new position.
- Drag gray mini-button up or down to reorder tracks.
- Type "?" for keyboard shortcuts.

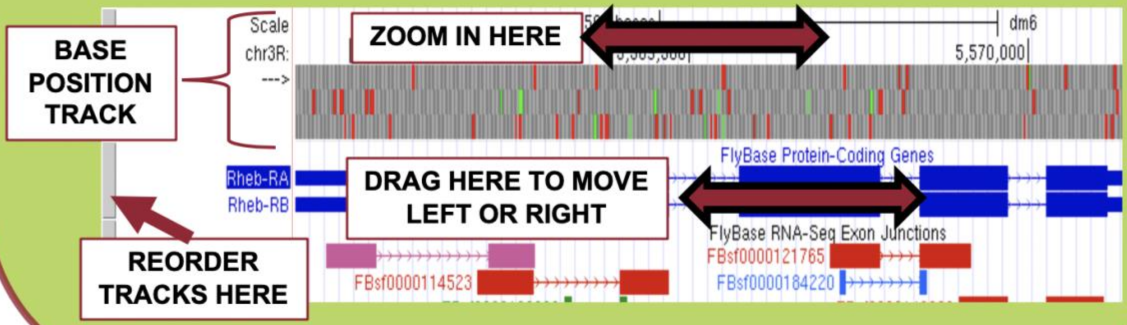

#### Part 3.1: Examine the evidence for a protein-coding gene in the *D. yakuba* region surrounding the *tblastn* alignment

FYI The accession number for the *D. yakuba* chromosome 3R scaffold (CM000160.2) and scaffold name (chr3R) are all synonymous (i.e., all are different ways to refer to the same region of the *D. yakuba* genome).

In the Genome Browser, either CM000160.2:17,358,666-17,359,556 or chr3R:17,358,666-17,359,556 could be entered in the "Position/Search Term" field text box and we would still navigate to the same genomic region.

1. Navigate to the Genome Browser.
2. Click on "D. yakuba" in the "REPRESENTED SPECIES" table.
3. Under the "D. yakuba Assembly" field, confirm that "May 2011 (WUGSC dyak_caf1/DyakCAF1)" is selected.
4. In Part 2, we determined the putative ortholog of Rheb-PA is located at approximately 17,358,666 – 17,359,556 on the chromosome 3R (chr3R) scaffold of the *D. yakuba* genome assembly. Enter "**chr3R:17,358,666-17,359,556**" under the "Position/Search Term" field to examine this region.
5. Click on the "Go" button (Figure 16).

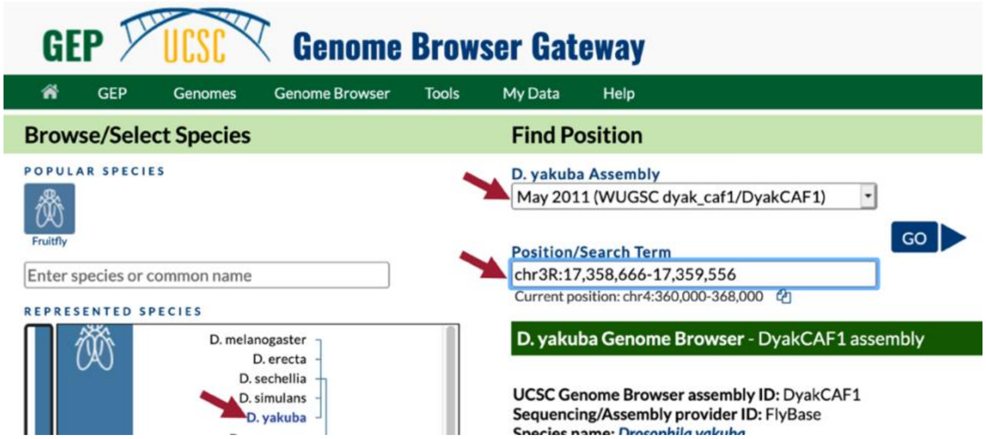

**Figure 16 Navigate to the region surrounding the best collinear set of alignments to the *D. melanogaster* Rheb-PA protein in the *D. yakuba* May 2011 (WUGSC dyak_caf1/DyakCAF1) assembly (i.e., chr3R:17,358,666-17,359,556).**

6. In the list of buttons below the Genome Browser image, click on "default tracks" (Figure 3).
7. Zoom out 3x.

In the Genome Browser image, we should now see the following tracks (Figure 17):

- *BLAT* Alignments of NCBI RefSeq Genes
- *Spaln* Alignment of *D. melanogaster* Proteins
- Gene Prediction Tracks:
  - *GeMoMa* Gene Predictions
  - *Geneid* Gene Predictions
  - *Augustus* Gene Predictions
- modENCODE RNA-Seq from Adult Females
- modENCODE RNA-Seq from Adult Males

FYI: **B**LAST-**L**ike **A**lignment **T**ool (BLAT) maps nucleotide or protein sequences against an assembly. BLAT is faster, but less sensitive, than BLAST.

The *Spaln* Alignment is similar to *blastx*.

Notice that the *BLAT* alignment for the coding regions of the *D. yakuba RefSeq* transcript XM_002097996 against chr3R line up with the *Spaln* alignment to the *D. melanogaster* proteins Rheb-PA and Rheb-PB. The coding portions of the first four alignment blocks are in congruence with the placements of the first four coding exons (CDS's) predicted by *GeMoMa*, *Geneid*, and *Augustus*. The coding portions of the last alignment block are in congruence with the predictions by *GeMoMa* and *Augustus*.

According to the *BLAT* alignment for the *RefSeq* transcript XM_002097996, the putative (probable) ortholog of Rheb-PA is located at chr3R:17,358,347-17,359,906 in *D. yakuba* and the region spanning chr3R:17,358,666-17,359,559, within the putative ortholog, corresponds to the alignment to the coding region of the *RefSeq* transcript.

Before we examine the genomic neighborhood of the putative ortholog, we need to identify the direction of transcription for the putative ortholog of Rheb-PA.

8. Zoom in on an intron of one of the gene predictions (e.g., *GeMoMa*) for chr3R:17,358,666-17,359,559.

Since the arrows within the introns of the gene prediction point to the right, we know the feature in this region is on the positive (top) strand of DNA.

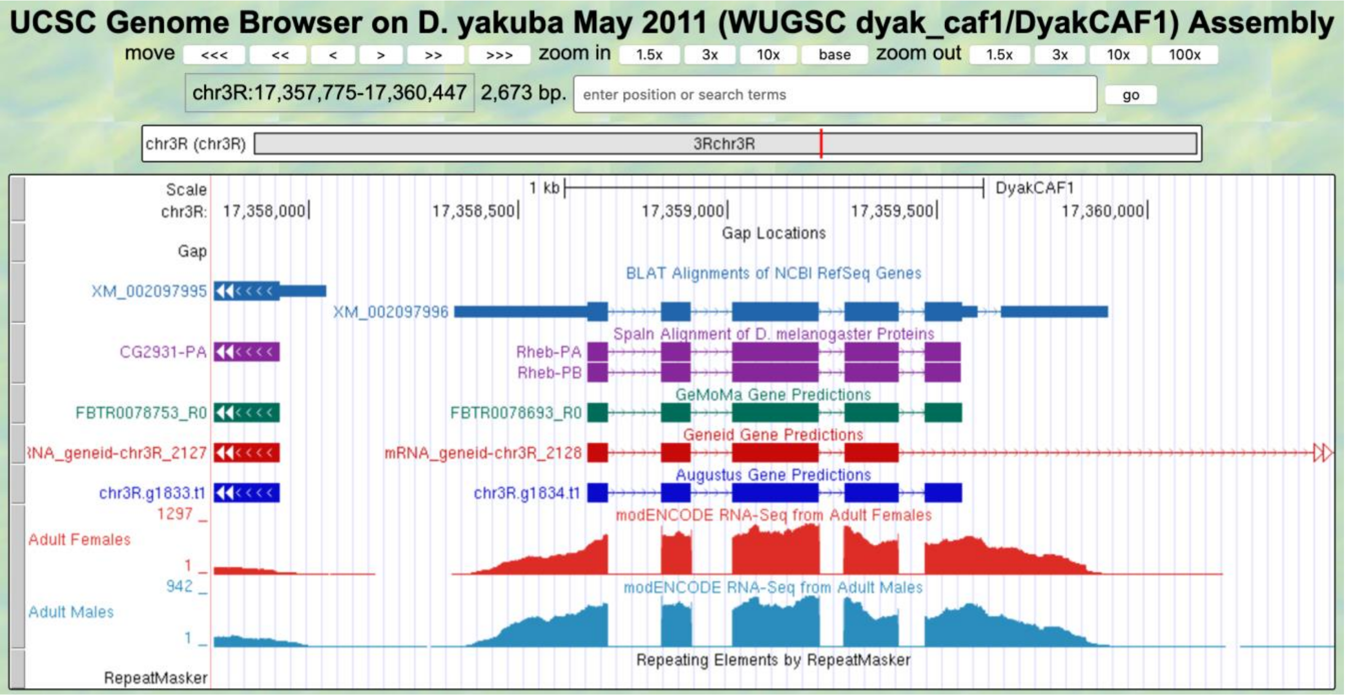

**Figure 17** Based on the *BLAT* Alignments of NCBI RefSeq Genes, *Spaln* protein alignments to both isoforms of *Rheb*, and the *GeMoMa*, *Geneid*, and *Augustus* gene predictions, we can infer that the feature in this region is on the positive strand.

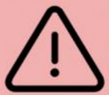
We cannot always trust that what we see in the Genome Browser is accurate, particularly for *Drosophila* species that are more distantly related to *D. melanogaster*. The gene prediction tracks, like their name implies, are predictions; thus, your role, as a researcher in the Pathways Project, is to help scientists studying these genes be confident in the specific model for the gene. Your brain is far superior to a computer algorithm in weighing conflicting evidence, thus your model will be more reliable than what a computer can produce alone.

For example, in your own project, there might be a situation where a gene predictor(s) does not show a gene in an area that has an alignment to *D. melanogaster* proteins (or vice versa); therefore, you'll need to investigate that further.

#### Part 3.2: Use synteny to gather additional evidence for the ortholog assignment

1. Zoom out another 3x (you should now be viewing chr3R:17,355,102-17,363,120).
2. In the "*BLAT* Alignments of NCBI RefSeq Genes" track shown in the Genome Browser image, click on "XM_002097995" (Figure 18).
   - Note: XM_002097995 is the accession number for the *D. yakuba RefSeq* transcript that is aligned to this region of chr3R by *BLAT*.

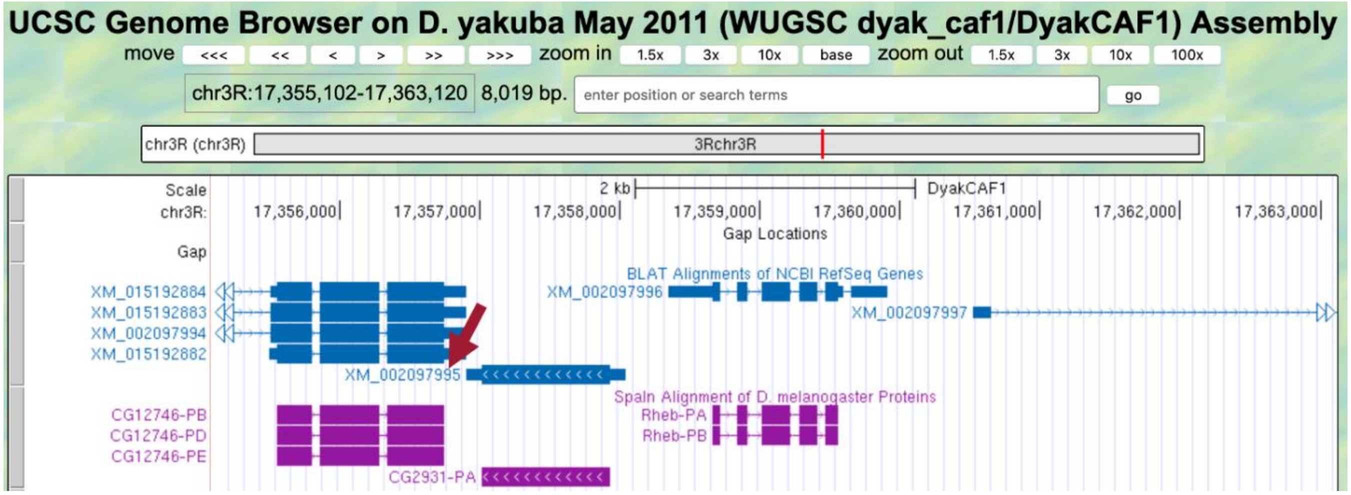

**Figure 18 Click on the *BLAT* Alignments of NCBI RefSeq Genes track for the nearest upstream neighbor.**

Notice the position of XM_002097995 is "chr3R:17,356,905-17,358,043" (Figure 19).

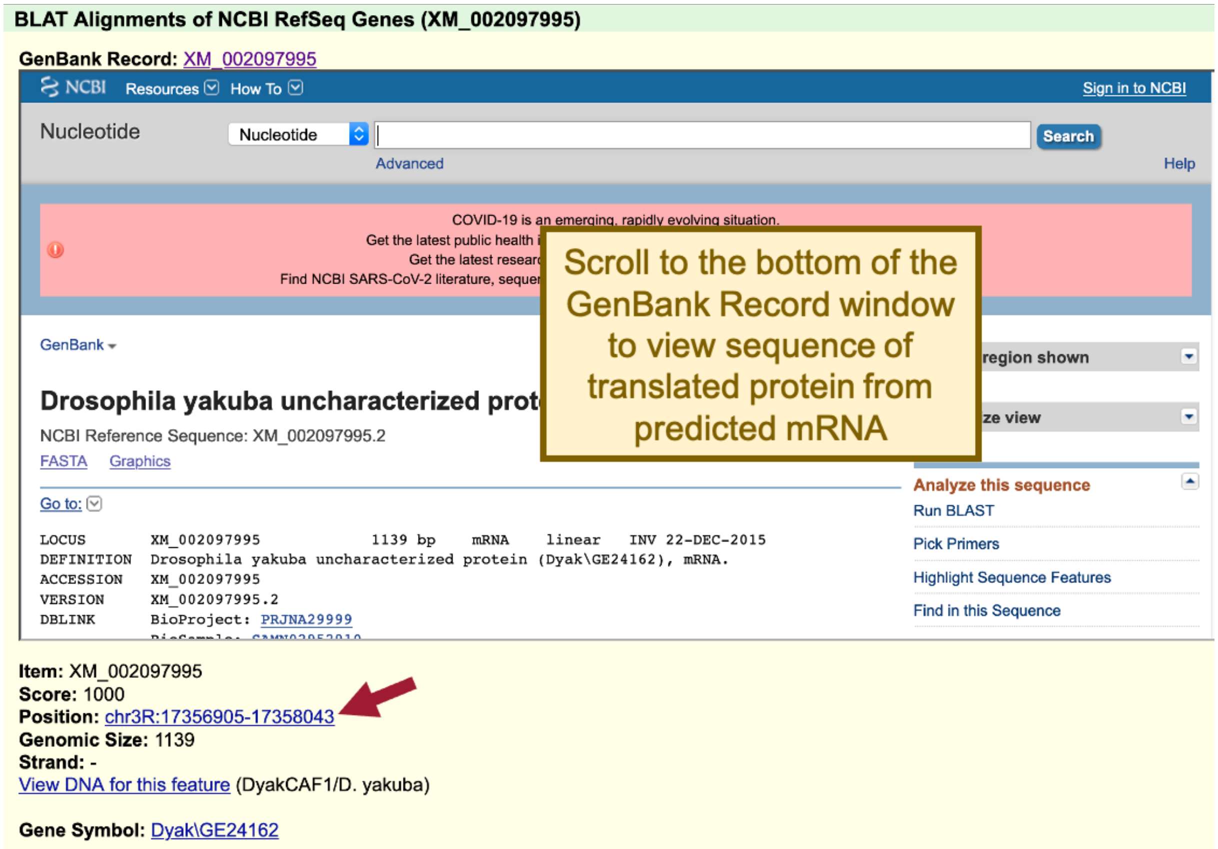

**Figure 19 *RefSeq* Gene feature XM_002097995 is located at chr3R:17,356,905-17,358,043.**

3. Scroll to the bottom of the GenBank Record window to the "translation" sequence within the "CDS" section (Figure 20).

We are now viewing the computationally predicted protein sequence of the nearest predicted upstream neighbor to Rheb-PA in *D. yakuba*.

4. Copy the accession number for the translated protein sequence (see arrow in Figure 20).

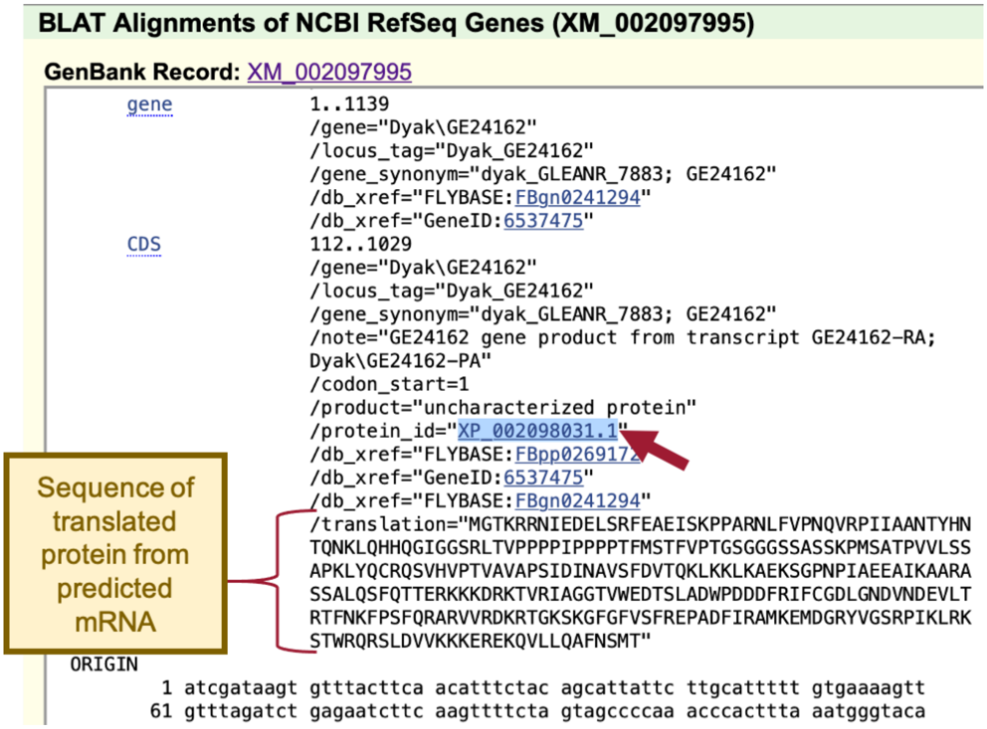

**Figure 20 Scroll to the bottom of the GenBank Record window for the *RefSeq* Genes feature (XM_002097995) to view the sequence of the translated protein. Copy the accession number for the translated protein sequence labeled "protein_id" (see arrow).**

When we run BLAST for this protein, we can enter the accession number "XP_002098031.1" and the program will use the translated protein sequence shown in Figure 20.

FYI The NCBI database of reference sequences (*RefSeq*) is a curated, non-redundant collection of naturally occurring DNA, RNA, and protein sequences. The RefSeq database includes both known (manually reviewed by NCBI staff or collaborators) and computationally predicted sequences.

The two-letter prefix (followed by an underscore) of RefSeq accession numbers has an implied meaning with respect to the type of molecule it represents (e.g., known or predicted model, genomic scaffold, mRNA).

| Accession Prefix | Description |
|---|---|
| XM_ | mRNA (predicted model) |
| XP_ | protein (predicted model) |
| NP_ | protein (known) |

**A complete list of RefSeq accession numbers is available in the NCBI Handbook.**

5. Navigate to NCBI *BLAST*.
6. Click on the "Protein *BLAST*" button (Figure 21).
   - Note: This is a *blastp* search (Part 2.2, review box).

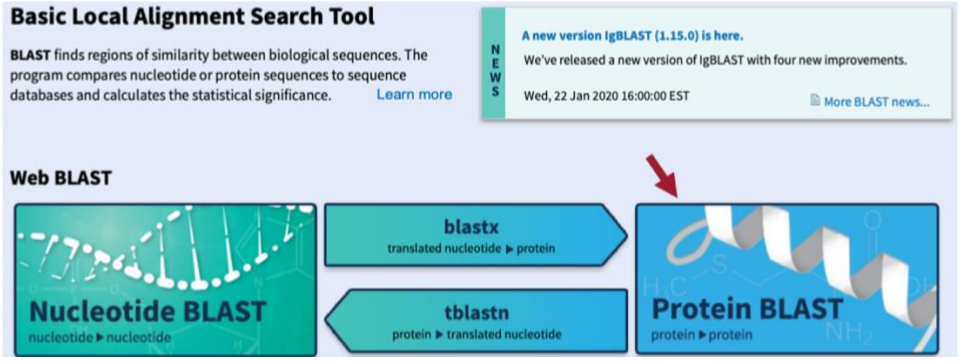

**Figure 21 Navigate to the NCBI *BLAST* website and then click on the "Protein *BLAST*" button.**

7. Paste the accession number for the translated protein sequence we copied from the GenBank Record for XM_002097995 into the "Enter Query Sequence" text box (Figure 22).
8. Click within the "Job Title" text box and it should automatically populate.
9. Under "Choose Search Set," select "Reference proteins (refseq_protein)" as the "Database" to search.
10. In the "Organism" text box, enter "**Drosophila melanogaster (taxid:7227)**."
11. Make sure the check box next to "Show results in a new window" is selected.
12. Click on the "*BLAST*" button.

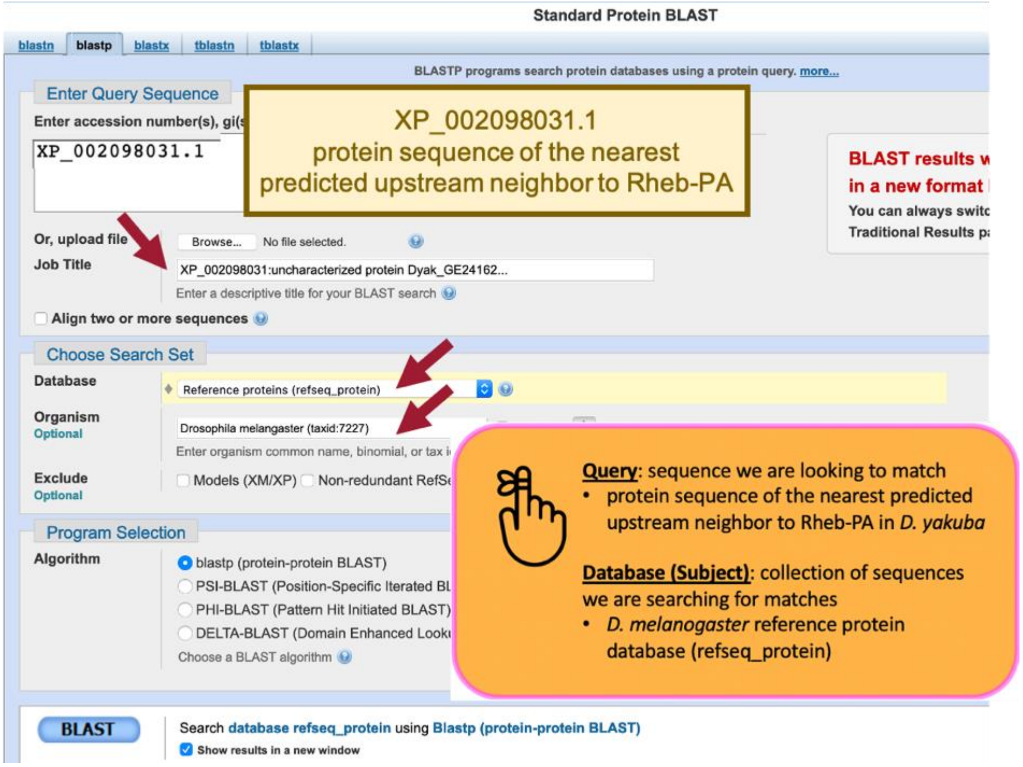

**Figure 22 Configure *blastp* to compare the protein sequence of the nearest predicted upstream neighbor to Rheb-PA in *D. yakuba* (query; XP_002098031.1) against the *D. melanogaster* reference protein database (database/subject).**

Our *blastp* search found 76 proteins within the *D. melanogaster* reference protein database that show similarities with the protein sequence of the nearest predicted upstream neighbor of Rheb-PA in *D. yakuba*; however, only one of these is a good hit (uncharacterized protein Dmel_CG2931 [Drosophila melanogaster]; accession: NP_649552.1, sequence identity of 96.72%, and an E-value of 0.0). The remaining hits had much higher E-values (2e-13 to 0.038) and much lower percent identities (50% to 22.69%) (Figure 23). Therefore, we can conclude that *CG2931* is the nearest upstream neighbor of *Rheb* in both *D. yakuba* and *D. melanogaster*. Furthermore, *CG2931* is located at approximately chr3R:17,356,905-17,358,043 in *D. yakuba*.

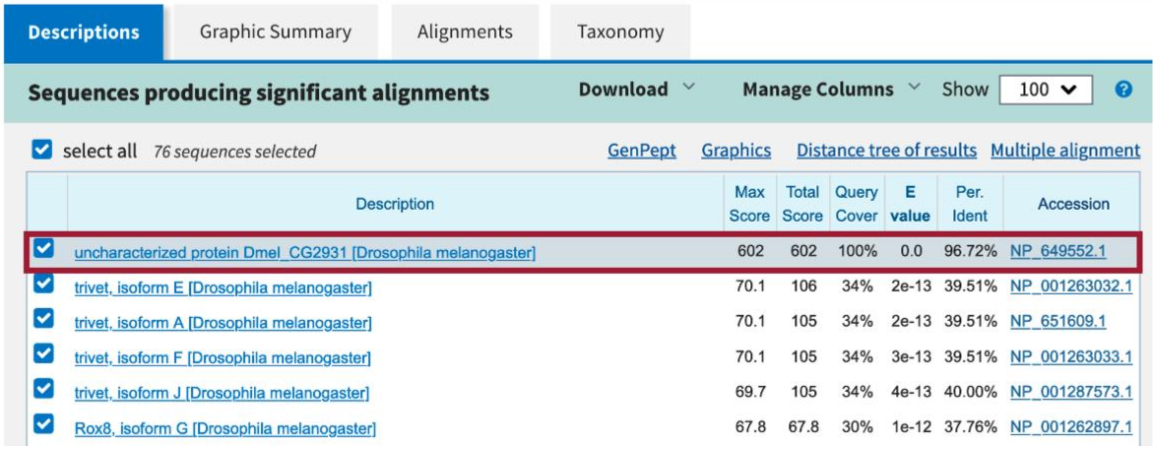

| Description | Max Score | Total Score | Query Cover | E value | Per. Ident | Accession |
|---|---|---|---|---|---|---|
| uncharacterized protein Dmel_CG2931 [Drosophila melanogaster] | 602 | 602 | 100% | 0.0 | 96.72% | NP_649552.1 |
| trivet, isoform E [Drosophila melanogaster] | 70.1 | 106 | 34% | 2e-13 | 39.51% | NP_001263032.1 |
| trivet, isoform A [Drosophila melanogaster] | 70.1 | 105 | 34% | 2e-13 | 39.51% | NP_651609.1 |
| trivet, isoform F [Drosophila melanogaster] | 70.1 | 105 | 34% | 3e-13 | 39.51% | NP_001263033.1 |
| trivet, isoform J [Drosophila melanogaster] | 69.7 | 105 | 34% | 4e-13 | 40.00% | NP_001287573.1 |
| Rox8, isoform G [Drosophila melanogaster] | 67.8 | 67.8 | 30% | 1e-12 | 37.76% | NP_001262897.1 |

**Figure 23 The best *blastp* match to the nearest predicted upstream neighbor of Rheb-PA in *D. yakuba* is "uncharacterized protein Dmel_CG2931 [Drosophila melanogaster]" (accession: NP_649552.1) with an E-value of 0.0 and a sequence identity of 96.72%.**

Now we need to repeat the *blastp* search for the other upstream neighboring gene (i.e., XM_015192884) and both downstream neighbors (i.e., XM_002097997 and XM_002097998) (Figure 24).

FYI For genes with more than one isoform, use one of the longer *BLAT* alignment for your *blastp*. However, if your BLAST results are ambiguous (i.e., not what you expected), try repeating the search with one of the other isoforms.

| | | Most Upstream | Nearest Upstream | Nearest Downstream | Most Downstream |
|---|---|---|---|---|---|
| **Target Species NCBI RefSeq Genes Accession (Protein Accession)** | | XM_015192884 (XP_015048370.1) | XM_002097995 (XP_002098031.1) | XM_002097997 (XP_002098033.2) | XM_002097998 (XP_002098034.1) |
| **BEST HIT** | **Description** | uncharacterized protein Dmel_CG12746, isoform E [Drosophila melanogaster] | uncharacterized protein Dmel_CG2931 [Drosophila melanogaster] | collapsin response mediator protein, isoform E [Drosophila melanogaster] | uncharacterized protein Dmel_CG2926, isoform A [Drosophila melanogaster] |
| | **Accession** | NP_649551.4 | NP_649552.1 | NP_730954.2 | NP_649554.1 |
| | **E-Value** | 0.0 | 0.0 | 0.0 | 0.0 |
| | **Percent Identity** | 84.30% | 96.72% | 99.66% | 86.18% |
| **2ND BEST HIT NOT an isoform of best hit** | **Description** | wunen, isoform B [Drosophila melanogaster] | trivet, isoform E [Drosophila melanogaster] | uncharacterized protein Dmel_CG6106, isoform B [Drosophila melanogaster] | toutatis, isoform E [Drosophila melanogaster] |
| | **E-Value** | 1e-15 | 2e-13 | 5e-13 | 2e-12 |
| | **Percent Identity** | 30.18% | 39.51% | 24.81% | 48% |

*The second-best hit reported in this table should not be an isoform of the best hit. For example, the best hit for the most downstream neighbor of Rheb-PA is "uncharacterized protein Dmel_CG2926, isoform A," the second-best hit is "uncharacterized protein Dmel_CG2926, isoform B," and the third best hit is "toutatis, isoform E." Since the second-best hit was an isoform of the best hit, "toutatis, isoform E" should be reported as the next best hit.

**Figure 24 Summary of the *blastp* search results for the protein sequences of the two nearest predicted upstream and downstream neighbors to Rheb-PA in *D. yakuba* against the *D. melanogaster* reference protein database.**

Now we need to draw a sketch of the genomic neighborhood of *Rheb* in *D. yakuba*. In order to do so, we need to repeat the process we followed in Part 1 for each of the neighboring genes in *D. yakuba* identified in our *blastp* search (i.e., zoom into an intron of each of the four neighboring genes' predictions and draw arrows in the correct directions on our sketch).

13. Zoom out far enough to examine the genomic region surrounding the putative *Rheb* ortholog.
14. Use the strategy described in Steps 10 – 11 of Part 1 to determine the orientations of the two nearest genes upstream and downstream of the putative *Rheb* ortholog (Figure 25).

| | Most Upstream | Nearest Upstream | Target Gene | Nearest Downstream | Most Downstream |
|---|---|---|---|---|---|
| ***D. melanogaster*** **Gene Symbol** | *CG12746* | *CG2931* | *Rheb* | *CRMP* | *CG2926* |
| **Strand in *D. melanogaster* (+/-)** | + | - | + | + | - |
| **Target Species Gene Symbol** | *CG12746* | *CG2931* | *Rheb* | *CRMP* | *CG2926* |
| **Strand in Target Species (+/-)** | + | - | + | + | - |

**Figure 25 The putative orthologs of *CG12746* and *CG2931* are located upstream of *Rheb*, and the putative orthologs of *CRMP* and *CG2926* are located downstream of *Rheb* in the *D. yakuba* chr3R scaffold, and each gene is on the +, -, +, and - strands, respectively.**

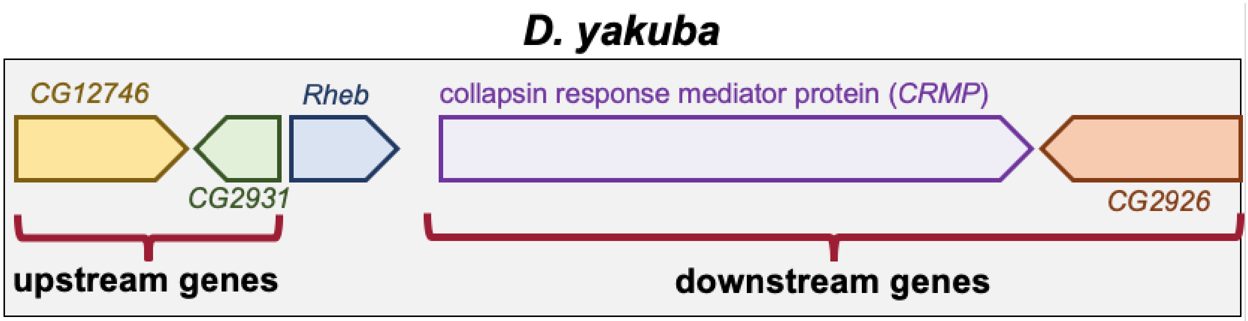

**Figure 26 Sketch of the genomic neighborhood of *Rheb* in *D. yakuba*.**

If the genomic neighborhood looks similar between *Rheb* in *D. melanogaster* (Figure 6) and the putative ortholog in *D. yakuba* (Figure 26), we can be confident we have found the true ortholog. However, if any of the information is inconsistent with this being a syntenic region, we should inspect our other hits in the *tblastn* search to see if a different genomic region is a better match overall.

Examination of the genomic regions surrounding the *Rheb* gene in *D. melanogaster* (Figure 27; top) and the putative *Rheb* ortholog in *D. yakuba* (Figure 27; bottom) shows that the relative gene order (i.e., *CG12746*, *CG2931*, *Rheb*, *CRMP*, and *CG2926*) and orientations (+, -, +, + , -) are the same in the two species. Hence the synteny analysis supports the assignment of the *D. yakuba* feature at chr3R:17,358,666-17,359,556 as an ortholog of *Rheb*.

- Note: If your target species' assembly happened to have been numbered from the opposite end of the relevant scaffold, the orientation (+ or – strand) of the orthologs could be the opposite (i.e., -, +, -, -, +) of what you see in *D. melanogaster* but still be syntenic.

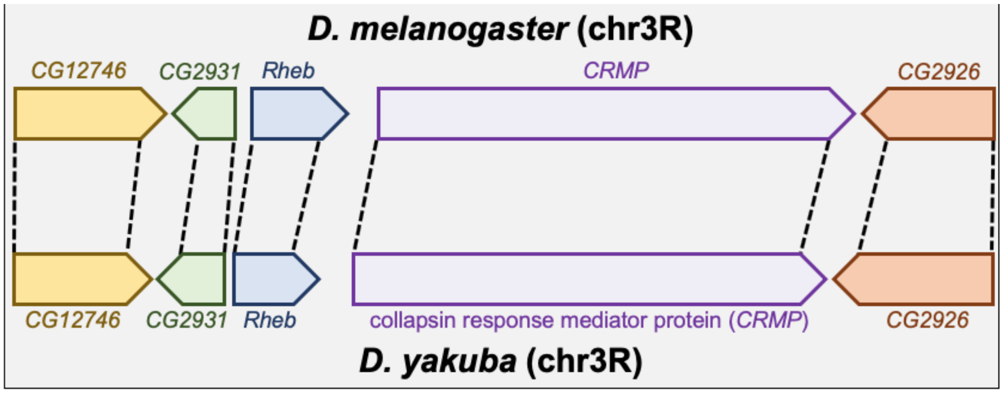

**Figure 27 Comparison of the relative order and orientation of the genomic neighborhoods of *Rheb* in *D. melanogaster* (top) and *D. yakuba* (bottom).**

#### Part 4: Determine the gene structure in *D. melanogaster*

In Part 4 we will use the Gene Record Finder, which is a web tool that enables us to quickly identify a unique set of exons for a given gene and to retrieve their Coding DNA Sequences (CDS's), also referred to as coding exons.

The Gene Record Finder will also provide details, such as number of isoforms, exon-intron structure and their coordinates, and transcript and protein information of the gene in question (in this case *Rheb*). **It is important to remember that the details provided by the Gene Record Finder are for the gene in the reference organism, *D. melanogaster*.** We will use the details from *Rheb* in *D. melanogaster* to assist us with creating a gene model for *Rheb* in *D. yakuba*.

Before we can construct the orthologous gene model, we need to ascertain the gene structure (e.g., number of isoforms and CDS's) of the *D. melanogaster Rheb* gene using the Gene Record Finder.

1. Open a new web browser tab and navigate to the Gene Record Finder.
2. Enter "**Rheb**" into the text box.
3. Click on the "Find Record" button (Figure 28).

> If we enter "rheb" in the Gene Record Finder, we will get the following error: *Cannot complete the request because of the following error: Cannot find the FlyBase gene record: rheb.*
>
> This is because gene symbols in *Drosophila* are **case-sensitive** (e.g., *Tor* and *tor* correspond to two different genes in *D. melanogaster*).

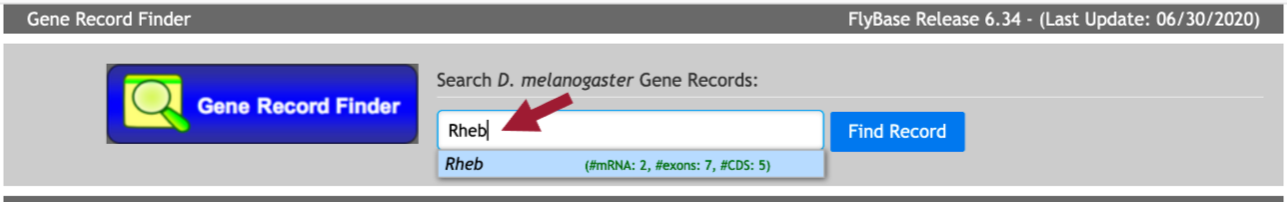

**Figure 28 Use the Gene Record Finder to retrieve the gene record for the *Rheb* gene in *D. melanogaster.***

The Gene Record Finder shows that *Rheb* has two isoforms (A and B) in *D. melanogaster*. A graphical overview of the two isoforms is shown in the "mRNA Details" panel. The "CDS usage map" (under the "Polypeptide Details" tab) shows that both isoforms have the same set of coding exons (CDS's) (i.e., 1_9855_0, 2_9855_2, 3_9855_2, 4_9855_1, and 5_9855_0). (The coding exons are ordered from 5' to 3' from left to right in the CDS usage map.) Hence the differences between these two isoforms are limited to the untranslated regions (UTRs) (Figure 29).

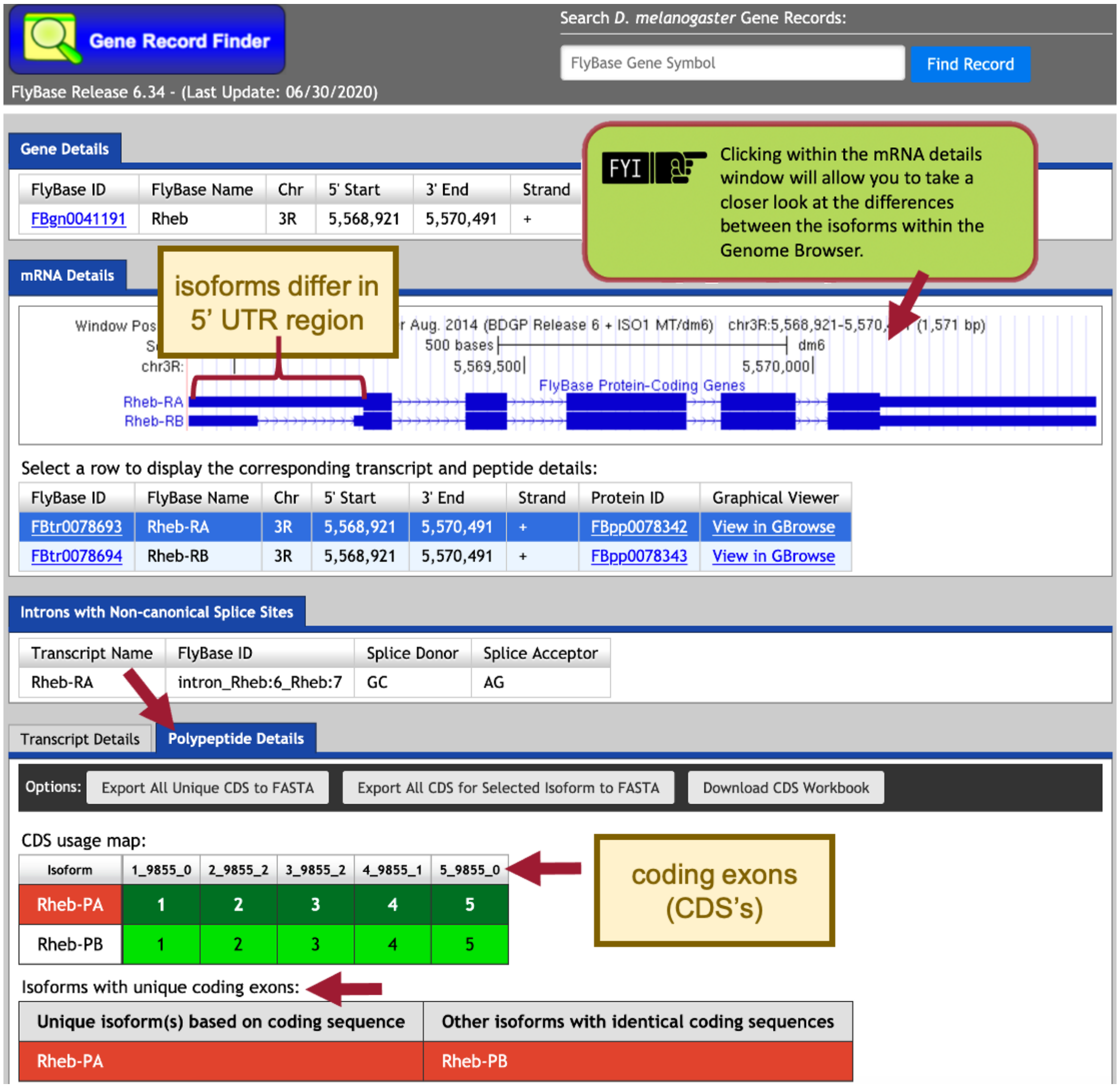

**Figure 29 The "mRNA Details" panel of the Gene Record Finder shows that the *Rheb* gene has two isoforms in *D. melanogaster* (i.e., Rheb-RA and Rheb-RB). Under the "Polypeptide Details" tab, the "CDS usage map" indicates that both isoforms have five coding exons (CDS's), and the "Isoforms with unique coding exons" section shows that both isoforms have identical coding sequences.**

Based on parsimony (i.e., minimizing the number of changes compared to *D. melanogaster*), we expect to find both the A and B isoforms of *Rheb* in our *D. yakuba* genome sequence. For this walkthrough, we will only focus on the annotation of the coding exons (CDS's), which do not include the UTRs. Consequently, we only need to determine the coordinates of the five coding exons (CDS's) for one of the isoforms (e.g., isoform A) because the set of coding exons for both the A and B isoforms are the same.

### Part 5: Determine the approximate location of the coding exons (CDS's) in *D. yakuba*

The initial *tblastn* search we performed in Part 2 helped define the search region for the putative ortholog within the genomic scaffold of the target genome, *D. yakuba*. The approximate coordinates of each CDS can now be determined by aligning each CDS of the gene in *D. melanogaster* against this search region of the target genome.

The next step in our analysis is to determine the approximate coordinates of each coding exon (CDS) of Rheb-PA in *D. yakuba*. Because the *BLAST* algorithm does not take the positions of potential splice sites within a complete protein sequence into account when it generates the alignment, *BLAST* often extends the alignment beyond the coding exon boundary and into the intron. To ameliorate this issue, the GEP annotation protocol recommends mapping each coding exon separately to determine their approximate locations and then further refine the exon boundaries by searching for compatible splice donor and acceptor sites through visual inspection using the Genome Browser.

In addition to comparing a query sequence against a collection of subject sequences in a database (e.g., Part 2 of this walkthrough), the NCBI *BLAST* web service also allows us to compare two or more sequences against each other (using the program *bl2seq* (*BLAST* 2 sequences)).

In order to map the amino acid sequences of each *D. melanogaster* CDS against the *D. yakuba* scaffold, we must translate the entire *D. yakuba* scaffold sequence in all six reading frames (i.e., three reading frames in the plus and minus strands, respectively) and then compare each conceptual translation against each CDS sequence from *D. melanogaster*. This means that we can use either *tblastn* or *blastx* to perform this search (depending on whether we treat the CDS sequence as the query or the subject sequence, respectively).

In this walkthrough, we will perform five *tblastn* searches— using each *D. melanogaster Rheb* CDS sequence as the query and the *D. yakuba* Accession Number (CM000160.2) we identified for the chr3R scaffold in Part 2.2 as the database (subject) sequence.

1. Navigate to the Gene Record Finder.
2. Enter "**Rheb**" into the text box.
3. Click on the "Find Record" button.

**Details provided by the Gene Record Finder are for the gene in *D. melanogaster*.**

Scroll down to the CDS usage map (under the "Polypeptide Details" tab). Since *Rheb* has 5 coding exons (CDS's), we will need to run five different *tblastn* searches, one for each of our CDS's. Let's start with CDS-1.

4. To view the protein sequence for CDS-1, select row 1 (FlyBase ID: 1_9855_0).
5. Copy the protein sequence (including the header) shown in the pop-up window (Figure 30).

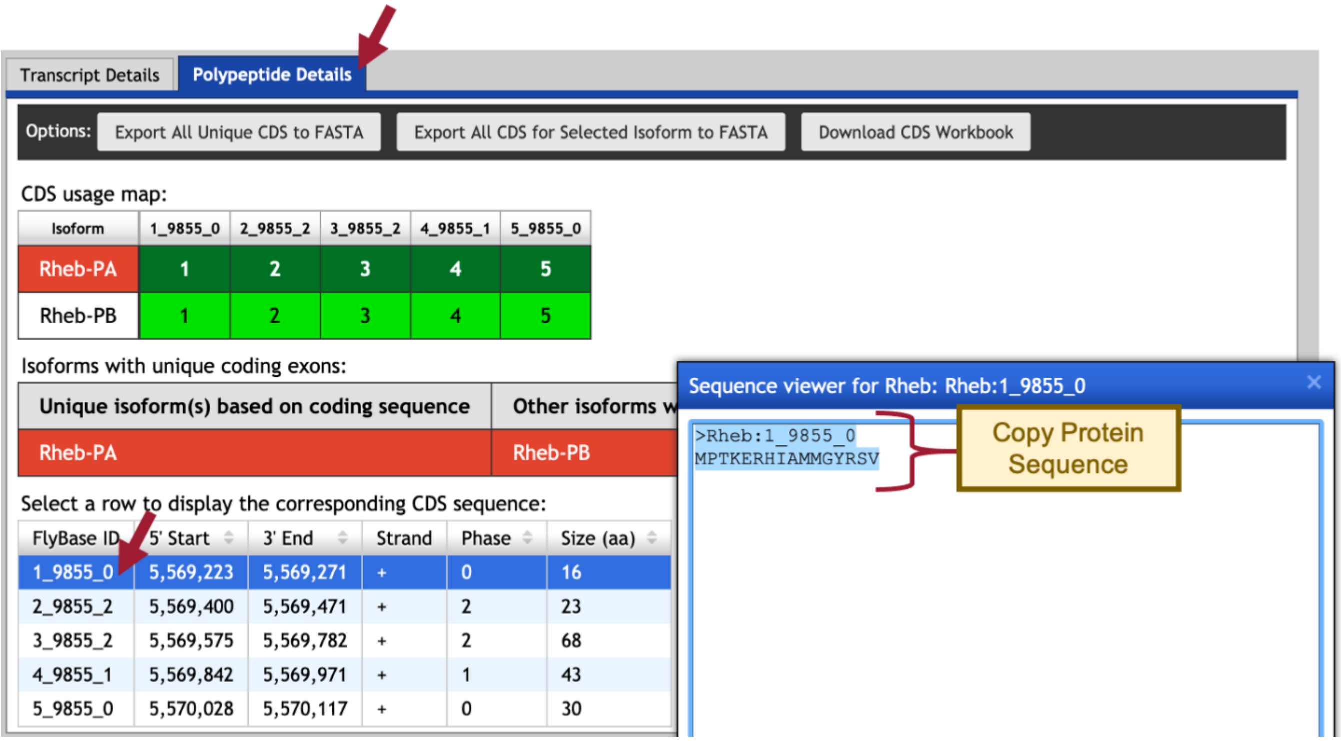

**Figure 30 Use the Gene Record Finder to retrieve the amino acid sequence for CDS-1 (FlyBase ID: 1_9855_0). To obtain the sequence for CDS-1, select row 1 (left) and then copy the protein sequence shown in the pop-up window (right).**

6. To setup the *tblastn* search, navigate to the NCBI *BLAST* website.
7. Click on the "*tblastn*" image under the "Web BLAST" section (Figure 31).

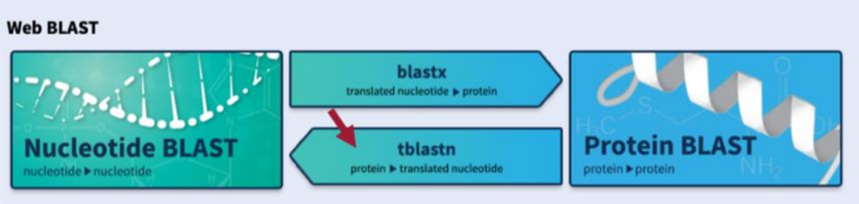

**Figure 31 Navigate to the NCBI BLAST website, and then click on the "*tblastn*" image.**

8. Select the "Align two or more sequences" checkbox (Figure 32).
9. Paste the sequence for CDS-1 into the "Enter Query Sequence" text box.
10. Click within the "Job Title" text box and it should automatically populate with the sequence header.
11. In the "Enter Subject Sequence" text box, enter the Accession Number for the chr3R sequence "**CM000160.2**" we identified in Part 2.2.
12. Based on our analysis in Part 2.3, limit the "Subject subrange" by entering from "**17300000**" to "**17400000**."
    - Note: Do NOT include commas or BLAST will search outside the subject subrange.

**Query**: sequence we are looking to match
- each individual CDS of *Rheb* in *D. melanogaster*

**Database (Subject)**: collection of sequences we are searching for matches
- *BLAST* will *translate* entire chr3R (CM000160.2) sequence of *D. yakuba*

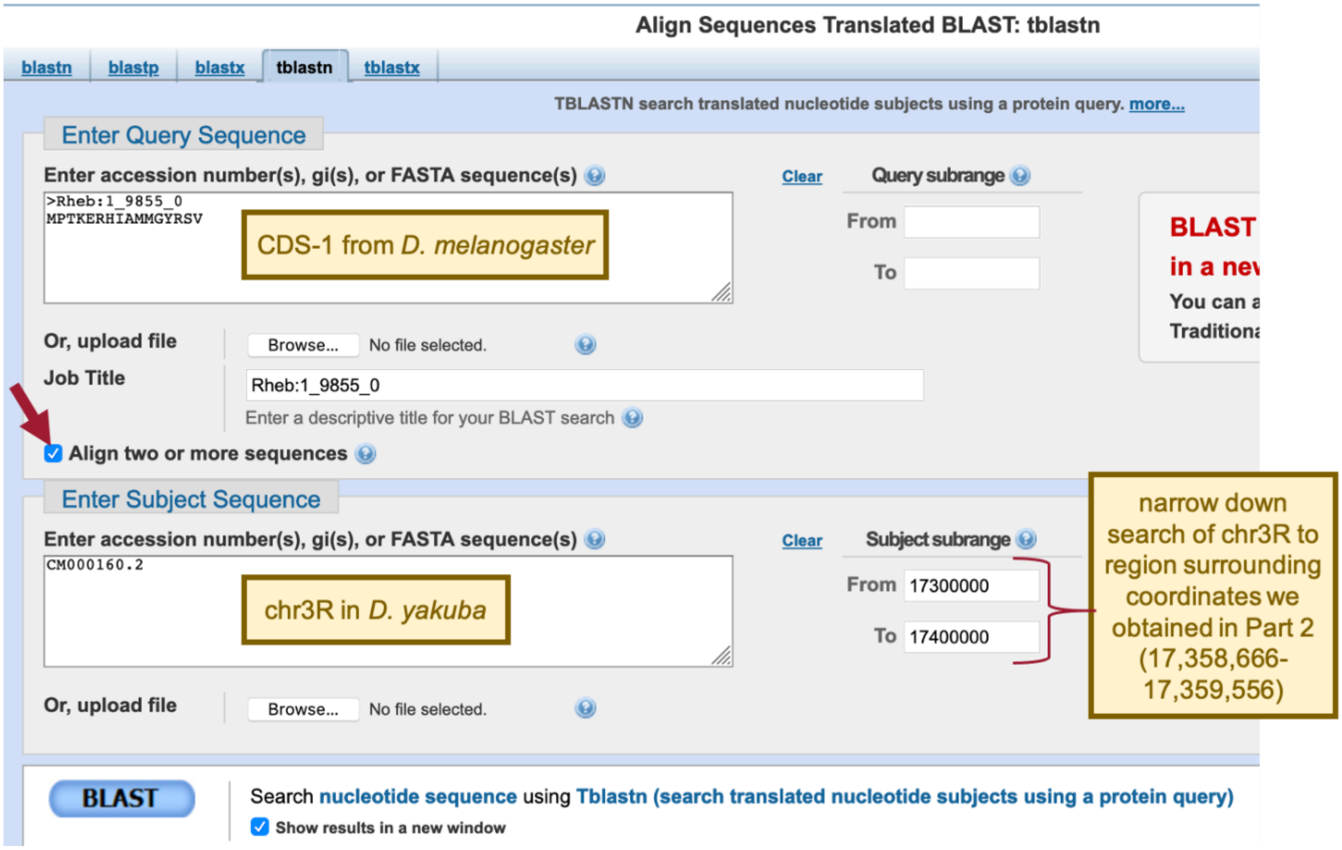

**Figure 32 Configure *tblastn* to compare the *D. melanogaster* CDS-1 (query) against the *D. yakuba* chr3R scaffold (accession: CM000160.2; subject). Since the *tblastn* search of Rheb-PA against the *D. yakuba* assembly placed the putative *Rheb* ortholog at approximately 17,358,666-17,359,556 on scaffold chr3R, the "Subject subrange" is used to limit the search region from 17,300,000 to 17,400,000. The smaller search region will increase the statistical power of the search and reduce the number of spurious matches.**

The default NCBI *BLAST* parameters are optimized for searching the query sequence against a large collection of sequences in a database. When we are using *BLAST* to compare only two sequences against each other, we need to change some of these alignment parameters because the default parameters could potentially mask the conserved regions of the coding exon.

13. Click on the "+" icon next to "Algorithm parameters" to expand the section (Figure 33).
14. In the "Scoring Parameters" section, change the "Compositional adjustments" field to "No adjustment."
15. In the "Filters and Masking" section, uncheck the "Low complexity regions" checkbox in the "Filter" field.
16. Make sure the check box next to "Show results in a new window" is selected.
17. Click on the "*BLAST*" button.

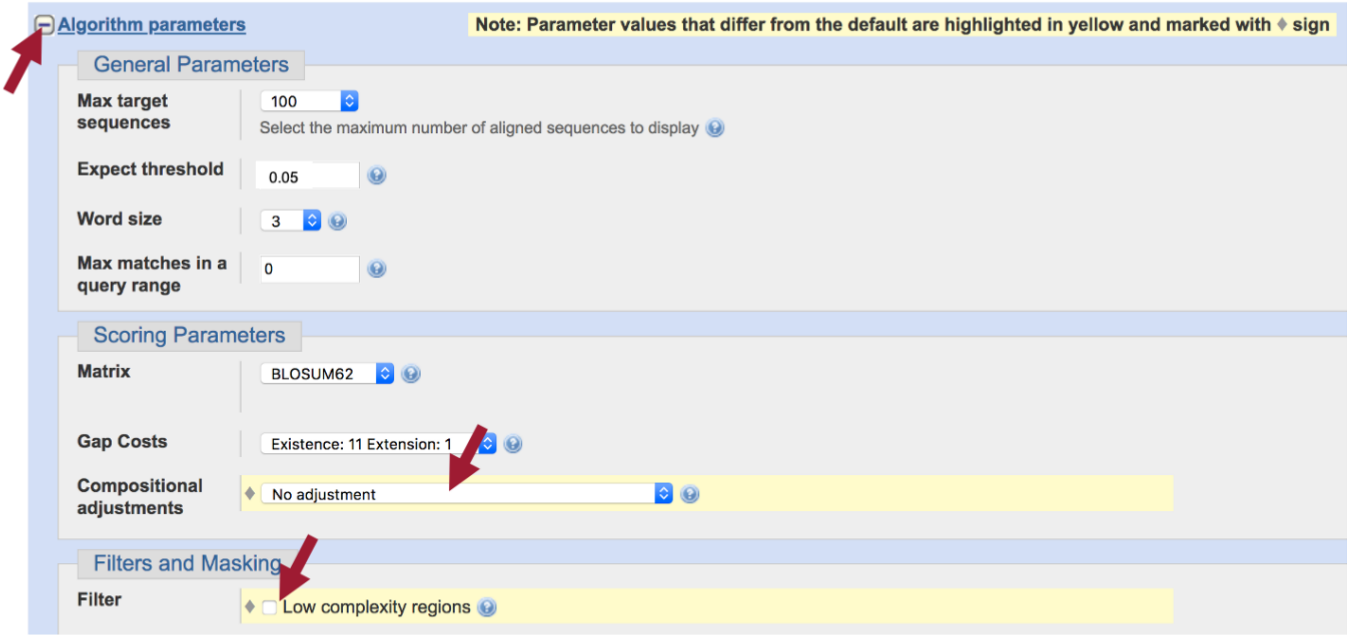

Figure 33 Expand the "Algorithm parameters" section, and then turn off "Compositional adjustments" and the filter for "Low complexity regions" to increase the sensitivity of the *tblastn* search.

The *tblastn* results show a single match (E-value: 5e-07; 94% sequence identity) to CDS-1.

18. Click on the "Alignments" tab to view the corresponding *tblastn* alignment (Figure 34).

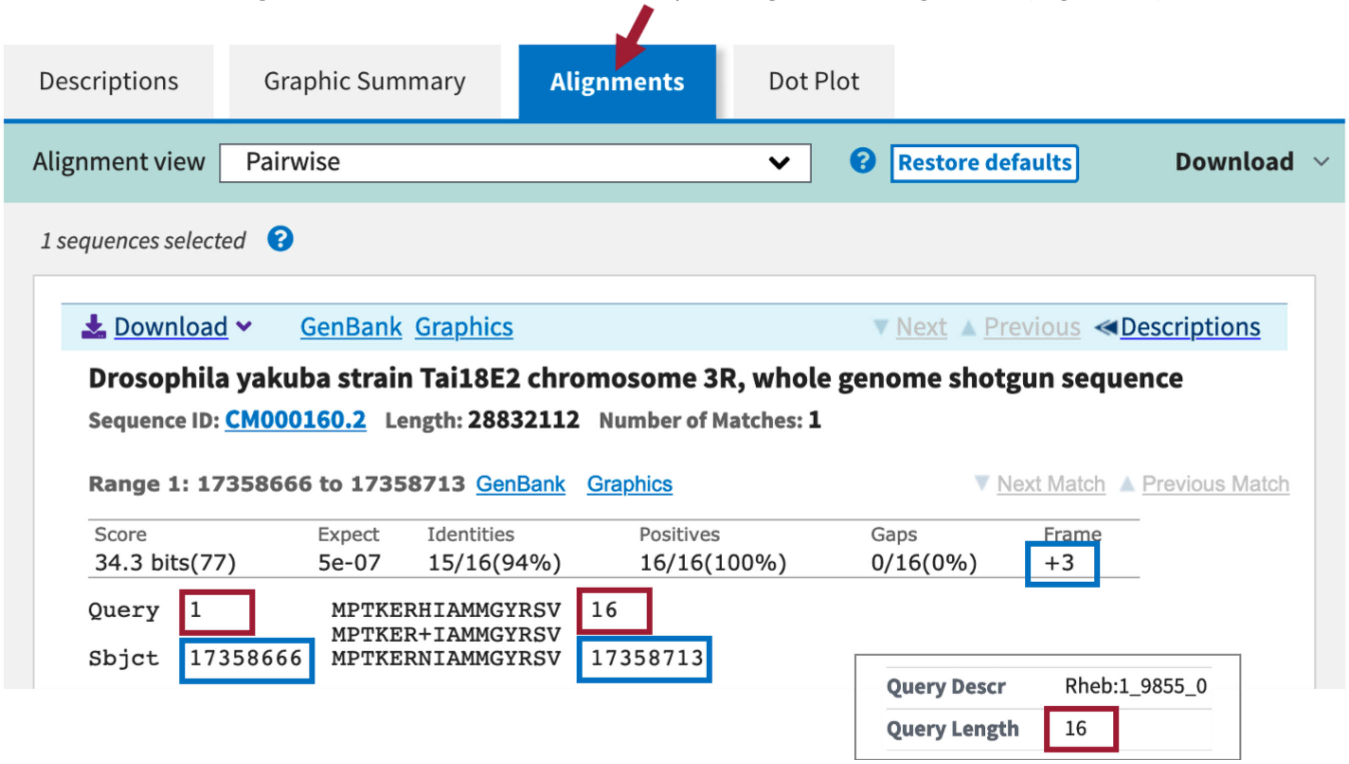

Figure 34 The *tblastn* alignment between the *Rheb* CDS-1 (query) and the *D. yakuba* scaffold (within the subject subrange of 17,300,000-17,400,000) is located at 17,358,666-17,358,713 when the sequence is translated in frame +3. The *tblastn* alignment has an E-value of 5e-07 and 94% sequence identity. This alignment covers all 16 aa of CDS-1.

The “Query” coordinates show that the alignment covers all 16 amino acids (aa) of CDS-1 (Figure 34).

- NOTE: We can find the length (in aa) of CDS-1 using the Gene Record Finder (CDS usage map under the “Polypeptide Details” tab) or at the top of the *tblastn* search results.

The “Subject” coordinates correspond to the region within chr3R (i.e., 17,358,666 – 17,358,713) that shows sequence similarity to CDS-1 when it is translated in the third reading frame in the positive strand (i.e., frame +3). Hence, we can place CDS-1 at 17,358,666 – 17,358,713 on chr3R.

We can apply this same procedure to place the other four CDS’s on the chr3R scaffold.

19. Copy the CDS-2 sequence (along with the header) from the Gene Record Finder.
20. Return to the *tblastn* search web browser tab and delete the CDS-1 sequence from the “Enter Query Sequence” textbox.
21. Paste the CDS-2 sequence in the textbox.
22. Click within the “Job Title” text box and it should automatically repopulate with the CDS-2 sequence header.
    - **Leave everything else the same as we had it for CDS-1.**
23. Click on the “*BLAST*” button to run the *tblastn* search.
24. Click on the “Alignments” tab to view the corresponding *tblastn* alignment.
25. Repeat this process to *BLAST* the remaining three CDS’s (Figure 35).

| CDS | Query Length (CDS Size) (aa) | *D. melanogaster* | | *D. yakuba* | | Subject Frame |
|---|---|---|---|---|---|---|
| | | Query Start | Query End | Subject Start | Subject End | |
| 1 | 16 | 1 | 16 | 17,358,666 | 17,358,713 | +3 |
| 2 | 23 | 1 | 23 | 17,358,844 | 17,358,912 | +1 |
| 3 | 68 | 1 | 68 | 17,359,013 | 17,359,216 | +2 |
| 4 | 43 | 1 | 43 | 17,359,279 | 17,359,407 | +1 |
| 5 | 30 | 1 | 30 | 17,359,470 | 17,359,559 | +3 |

**Figure 35 Summary of the *tblastn* CDS-by-CDS search results.**

Examination of the subject ranges for the *tblastn* alignments of the five CDS’s of *Rheb* shows that they are collinear—CDS’s 1-5 are placed on the positive strand and the subject ranges for the CDS's are in ascending order. Consequently, the CDS-by-CDS search results support the hypothesis that the putative (probable) ortholog of Rheb-PA is located at approximately 17,358,666-17,359,559 on chr3R of the *D. yakuba* genome assembly.

### Part 6: Refine coordinates of coding exons (CDS's)

Now that we've mapped each CDS separately to determine their approximate locations (Figure 35), we will now further refine the CDS boundaries by searching for compatible splice donor and acceptor sites by visual inspection using the Genome Browser.

As part of the modENCODE project, RNA-Seq data for *D. yakuba* was generated using samples from adult females and males. These RNA-Seq reads (100–125 bp in length) are derived primarily from processed mRNA (i.e., after the introns have been removed). Hence, genomic regions with RNA-Seq read coverage usually correspond to transcribed exons, which include both the translated and untranslated regions.

The "modENCODE RNA-Seq" tracks correspond to the samples from adult females (red) and males (blue) where RNA-Seq data is available (Figure 36). The height of the histograms within each track corresponds to the number of RNA-Seq reads that have been mapped to each position of the *D. yakuba* scaffold. By default, the scale of the "RNA-Seq Coverage" track will change automatically based on the minimum and maximum read depth within the genomic region being viewed.

If the number of RNA-Seq reads differs between samples (e.g., adult females vs. adult males), that would indicate alternative splicing.

**See Understanding Eukaryotic Genes Module 6 to review alternative splicing.**
**See RNA-Seq Primer (automatic download) for an overview of the different types of RNA-Seq data available on the Genome Browser.**

#### Part 6.1: Verify the start codon coordinates for Rheb-PA

Introns are removed from the pre-mRNA prior to translation. The processed mRNA also includes a 5' cap at the 5' end and a poly-A tail at the 3' end. This walkthrough focuses on annotation of the coding regions so we will not annotate the untranslated regions (UTRs) or the transcription start site (TSS).

- **start codon**: first codon of a coding sequence; signals the ribosome to begin translation of the mRNA; ATG (AUG) codes for Methionine (M)
- **stop codon**: marks the termination of translation; sometimes called "nonsense codons," since they do not specify any amino acid

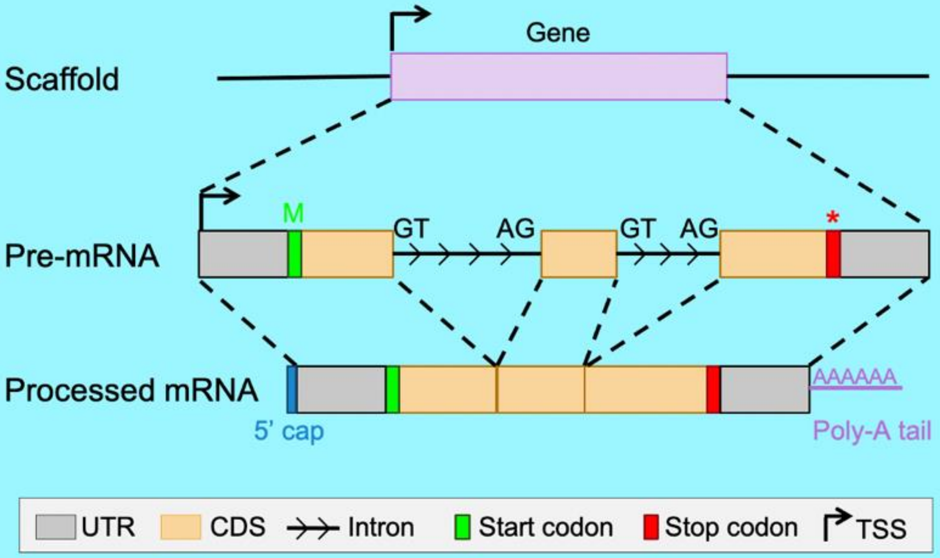

We need to ascertain whether the *tblastn* alignment for CDS-1 at 17,358,666-17,358,713 (Figure 34) is supported by the RNA-Seq data and determine the location of the start codon.

1. Return to the Genome Browser for *D. yakuba*.
2. Under the "Mapping and Sequencing Tracks," change the "Base Position" track to "full."
3. Click on the "refresh" button.
4. To examine the region of the *tblastn* alignment for CDS-1, enter "**chr3R:17,358,666-17,358,713**" into the "enter position or search terms" text box.

The RNA-Seq tracks for both samples show high RNA-Seq read depth within the *tblastn* alignment block (17,358,666-17,358,713), consistent with the hypothesis that this region is being transcribed in *D. yakuba*.

5. To examine the region surrounding the start of the *tblastn* alignment to CDS-1, enter "**chr3R:17,358,666**" into the "enter position or search terms" text box.
6. Click on the "go" button.
7. Zoom out 3x and another 10x.

In Part 5, we found our *tblastn* alignment for CDS-1 begins at 17,358,666 when translated in frame +3 (Figure 34). Examination of this region using the Genome Browser shows us that a start codon at this location is supported by multiple evidence tracks—the *BLAT* Alignments of NCBI RefSeq Genes and *Spaln* alignment of *D. melanogaster* proteins, as well as the *GeMoMa*, *Geneid*, and *Augustus* gene predictions (Figure 36).

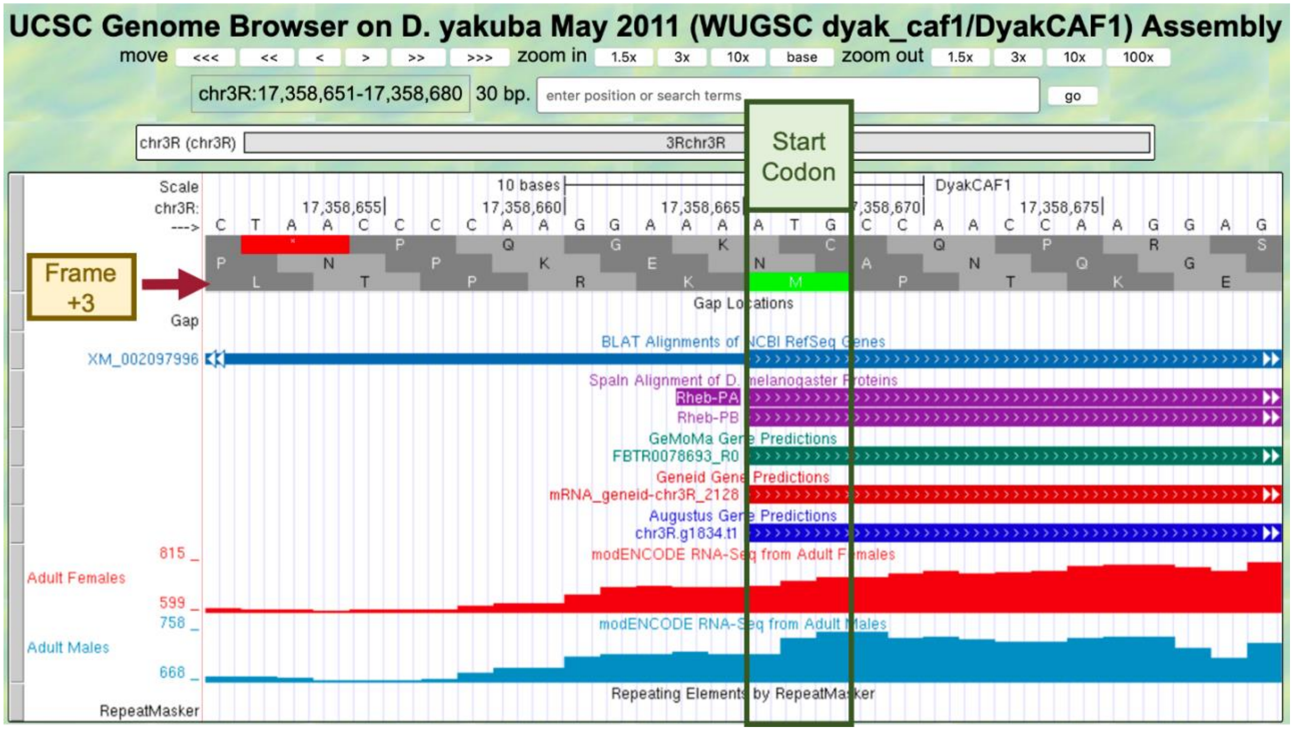

**Figure 36 The start codon at chr3R:17,358,666-17,358,668 is supported by the *BLAT* Alignments of NCBI RefSeq Genes, *Spaln* alignment of *D. melanogaster* proteins, and the GeMoMa, Geneid, and Augustus gene predictions.**

Since the RNA-Seq read coverage from the adult females and adult males samples extend upstream beyond the start codon, we need to determine if there are alternative start codon candidates available that could serve as the translation start site for Rheb-PA.

8. Zoom out 3x and another 10x.

The contiguous RNA-Seq read coverage extends upstream of the start codon at 17,358,666-17,358,668 to position 17,358,293 in the adult males' sample (Figure 37). Examination of the "Base Position" track shows that the first stop codon upstream from 17,358,666-17,358,668 occurs at 17,358,477-17,358,479 (remember that we are looking at frame +3 and that a codon consists of three amino acids). Since there are no alternative start codons in frame +3 between this stop codon and the start codon at 17,358,666-17,358,668, the most likely translation start site for the Rheb-PA ortholog in *D. yakuba* is assigned to the position 17,358,666-17,358,668 on chr3R. Consistent with the gene model for Rheb-RA in *D. melanogaster*, the RNA-Seq read coverage extending upstream of the start codon likely corresponds to the 5' untranslated region (UTRs) of the initial exon of *Rheb*.

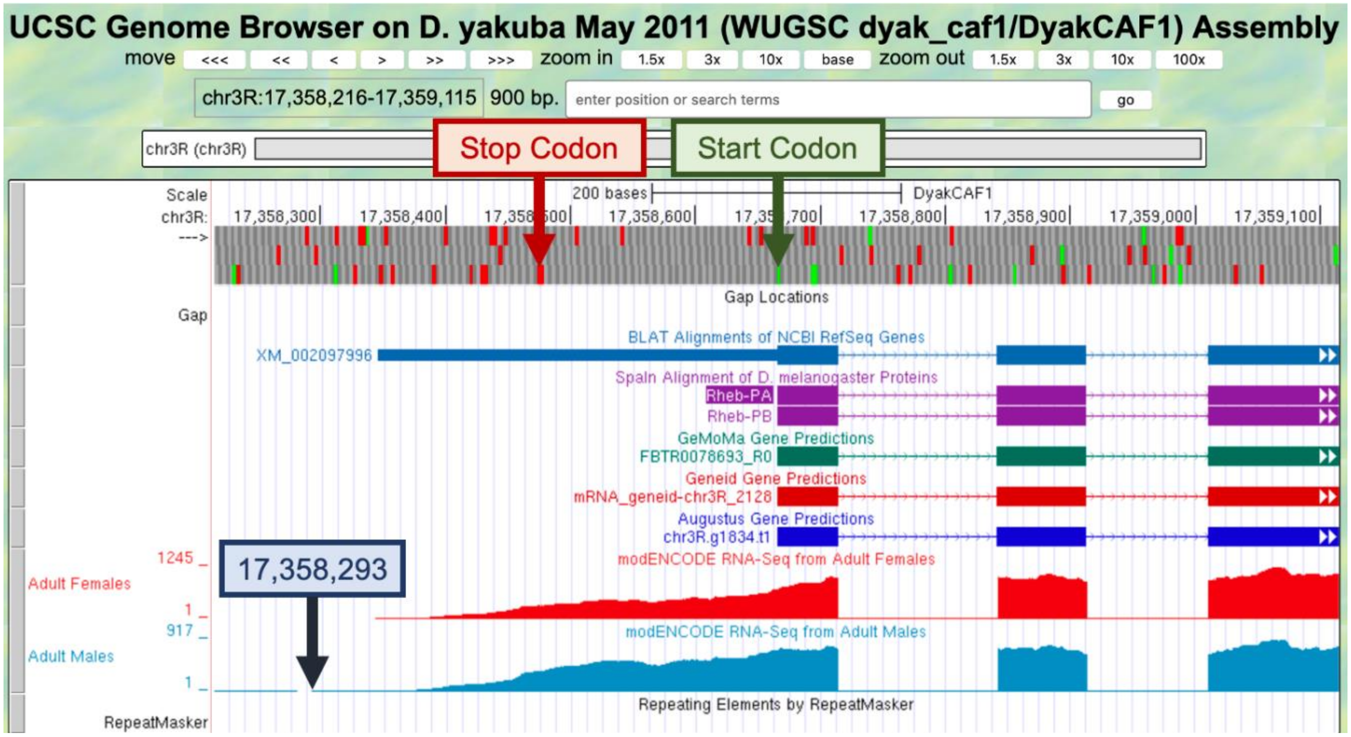

**Figure 37 The contiguous RNA-Seq read coverage extends upstream of the start codon at chr3R:17,358,666-17,358,668 to position 17,358,293 in the adult males' sample (blue box). Within the region from the start of the RNA-Seq read coverage in the adult males' sample to just before the start codon (i.e., 17,358,293-17,358,665), the first in-frame stop codon is located at 17,358,477-17,358,479 (red box). Since there are no alternative start codons in frame +3 between this stop codon and the start codon at 17,358,666-17,358,668, the translation start site for the Rheb-PA ortholog in *D. yakuba* is assigned to the position 17,358,666-17,358,668 on chr3R.**

#### Part 6.2: Verify the stop codon coordinates for Rheb-PA

In Part 5, we found our *tblastn* alignment for CDS-5 ends at 17,359,559 when translated in frame +3. The alignment covers all 30 amino acids of CDS-5 and ends with a stop codon (Figure 38).

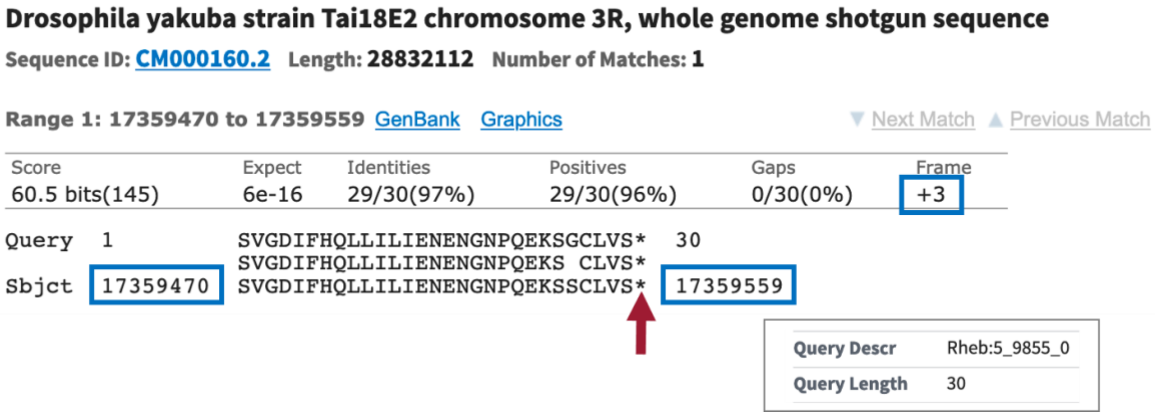

**Figure 38 The *tblastn* alignment of CDS-5 of Rheb-PA (query) against the *D. yakuba* scaffold chr3R (subject) placed the CDS at 17,359,470 – 17,359,559 when the sequence is translated in frame +3. The *tblastn* alignment covers all 30 amino acids of CDS-5, and it ends with a stop codon ('*'; red arrow).**

1. To examine the genomic region surrounding the end of the *tblastn* alignment to CDS-5, enter "**chr3R:17,359,559**" into the "enter position or search terms" text box.
2. Click on the "go" button.
3. Zoom out 3x and another 10x (Figure 39).

The stop codon at 17,359,557-17,359,559 is consistent with the *BLAT* Alignments of NCBI *RefSeq* Genes, *Spaln* alignment of *D. melanogaster* proteins, and the *GeMoMa* and *Augustus* gene predictions; however, the stop codon is inconsistent with the *Geneid* prediction mRNA_geneid-chr3R_2128, which predicted an intron in this region. However, the coding span for this *Geneid* prediction extends from 17,358,666-17,366,011, which encompasses regions with sequence similarity to the coding exons of the *CRMP* gene. Based on the *tblastn* alignment for CDS-5 and the available evidence on the Genome Browser, the stop codon for the Rheb-PA ortholog is placed at 17,359,557-17,359,559, and the last codon (S; Serine) before the stop codon ends at 17,359,556 (Figure 39).

Note that the RNA-Seq read coverage tracks for both samples indicate that transcription extends beyond the stop codon. Based on the gene structure of Rheb-RA in *D. melanogaster*, the region with RNA-Seq read coverage that extends beyond the stop codon likely corresponds to the 3' untranslated region (UTR) of the last exon in *Rheb*.

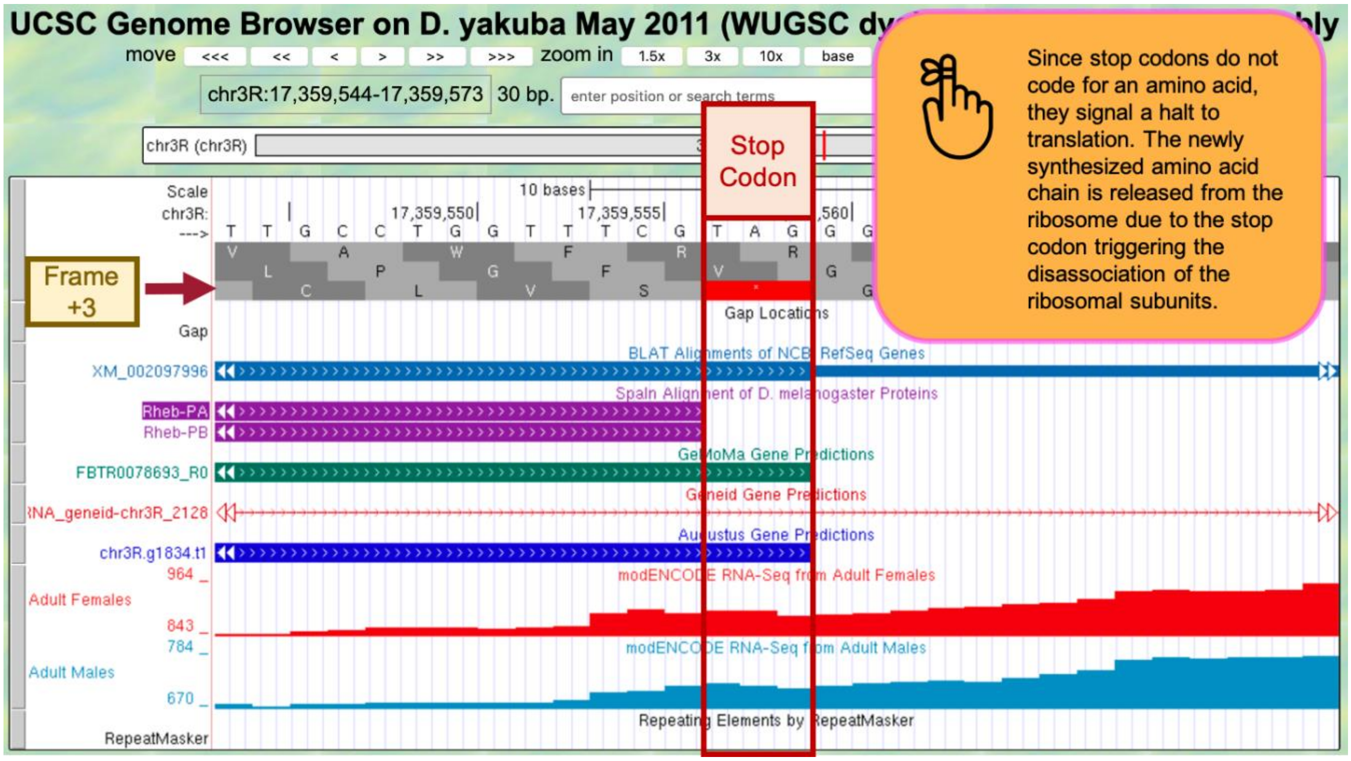

**Figure 39 Based on the *tblastn* alignment for CDS-5 and the available evidence on the Genome Browser, the stop codon for the Rheb-PA ortholog is placed at 17,359,557 – 17,359,559, and the last codon (S; Serine) before the stop codon ends at 17,359,556.**

#### Part 6.3: Determine the phases of the donor and acceptor splice sites

During splicing, introns are spliced out (removed) from the pre-mRNA so that adjacent exons are placed next to each other. This means that the ends of an exon do not necessarily correspond to the ends of the complete codon. The number of nucleotides between the last complete codon and the splice donor site is known as the **phase** of the splice donor site. Similarly, the number of nucleotides between the splice acceptor site and the first complete codon is known as the phase of splice acceptor site. Because the phases of the splice sites depend on the placement of the complete codon, the phases of the donor and acceptor sites are based on the reading frame of each CDS.

In addition, in order to maintain the open reading frame (ORF) across adjacent CDS's, the phases of the donor and acceptor sites of adjacent CDS's must be **compatible** with each other. Specifically, the **sum of the donor and acceptor phases of adjacent CDS's must either be 0 (i.e., no additional codon) or 3 (i.e., a complete codon)**. The use of incompatible splice donor and acceptor sites will introduce a frame shift into the translation of the CDS following the splice acceptor site.

In *D. melanogaster*, approximately 99% of introns have a **GT** splice donor site and 1% have a **GC** non-canonical splice donor site. Almost all introns have an **AG** splice acceptor site. The GEP comparative annotation protocol posits that all introns have a GT splice donor site and an AG splice acceptor site unless the *D. melanogaster* gene model uses a non-canonical splice site, or the non-canonical splice site is supported by RNA-Seq data.

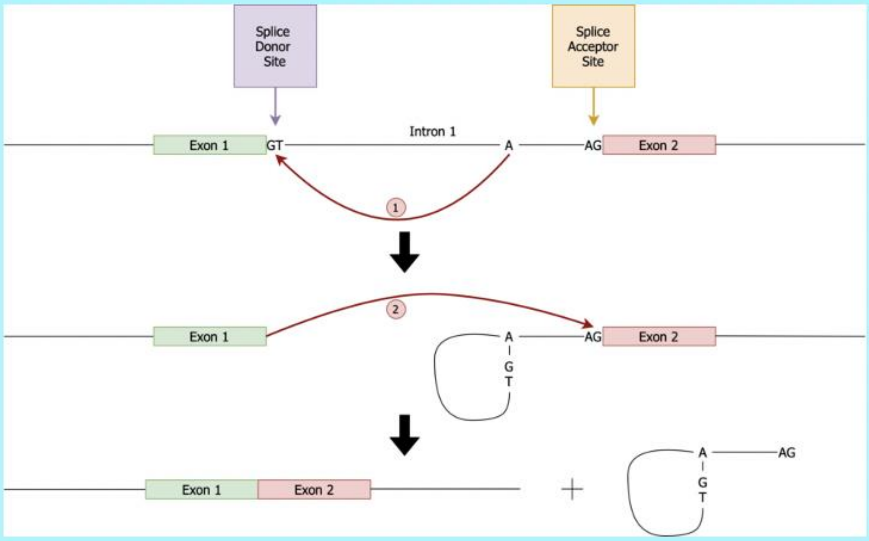

See [Understanding Eukaryotic Genes] Module 4 to further review splicing.

Because the *tblastn* alignment for CDS-1 of *Rheb* terminates at 17,358,713 and the alignment includes the last amino acid of the CDS (Figure 34), we expect to find the splice donor site for CDS-1 at around position 17,358,713.

1. To examine the genomic region surrounding the splice donor site of CDS-1, enter "**chr3R:17,358,713**" into the "enter position or search terms" text box.
2. Click on the "go" button.
3. Zoom out 3x and another 10x to examine the 30 bp surrounding this position.

The GT splice donor site closest to 17,358,713 is located at 17,358,715 – 17,358,716. This splice donor site is in phase 0 relative to frame +1 (Figure 40, top), in phase 2 relative to frame +2 (Figure 40, middle), and in phase 1 relative to frame +3 (Figure 40, bottom). This splice donor site is supported by multiple lines of evidence—*BLAT* Alignments of NCBI RefSeq Genes, *Spaln* alignment of *D. melanogaster* proteins, and the *GeMoMa*, *Geneid*, and *Augustus* gene predictions, as well as the RNA-Seq read coverage from samples of adult females and adult males.

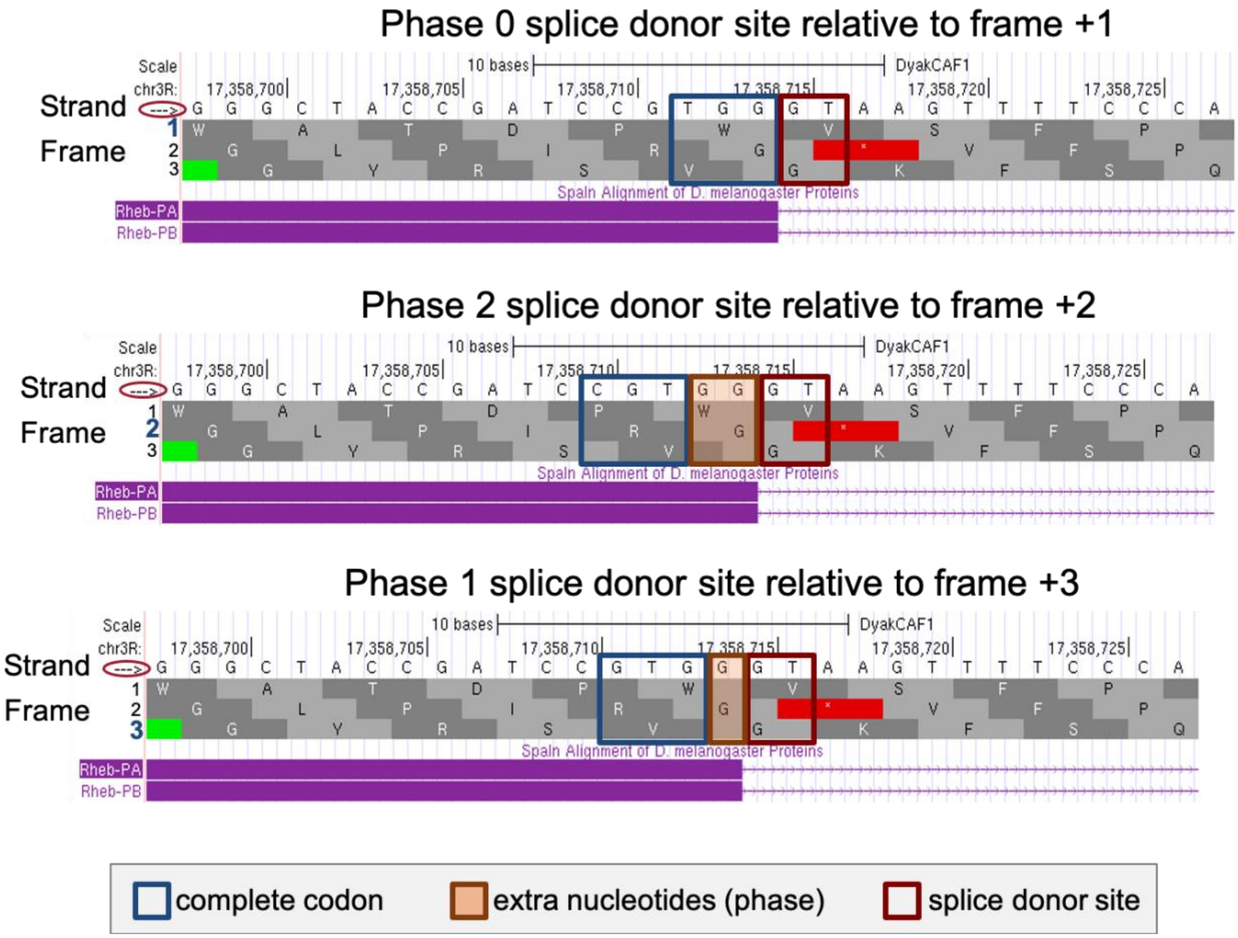

**Figure 40 There are three possible phases for the splice donor site (0, 1, or 2), which depend on the reading frame. This splice donor site (at 17,358,715 – 17,358,716) is supported by multiple lines of evidence. Since CDS-1 is in frame +3 and the last complete codon (GTG which codes for V) is located at 17,358,711 – 17,358,713, there is one nucleotide (G at 17,358,714) between the last complete codon and the splice donor site. Hence, this splice donor site is in phase 1 (bottom).**

Based on our *tblastn* alignment in Part 5, CDS-1 is in frame +3.

1. Zoom out far enough to see the entire length of CDS-1 and confirm that frame +3 has an open reading frame (ORF) (i.e., no stop codons are shown within frame +3 of CDS-1).

Since CDS-1 is in frame +3, the splice donor site at 17,358,715 – 17,358,716 is in phase 1. This means that the splice acceptor site of CDS-2 must be in phase 2 in order to maintain the open reading frame. Based on our *tblastn* alignment in Part 5, we placed CDS-2 at 17,358,844 – 17,358,912 in frame +1 (Figure 35).

2. To examine the genomic region surrounding the splice acceptor site of CDS-2, enter "**chr3R:17,358,844**" into the "enter position or search terms" text box.
3. Click on the "go" button.
4. Zoom out 3x and another 10x to examine the 30 bp surrounding this position (Figure 41).

There is only one potential canonical splice acceptor site (AG) in the 30 bp region surrounding the start of the *tblastn* alignment to CDS-2. This potential splice acceptor site, located at 17,358,840 – 17,358,841, is supported by the *BLAT* and *Spaln* alignments, and the *GeMoMa*, *Geneid*, and *Augustus* gene predictions, as well as the RNA-Seq read coverage from samples of adult females and adult males.

Since CDS-2 is in frame +1 and the first complete codon (AAA which codes for K) is located at 17,358,844 – 17,358,846, there are two nucleotides (GC at 17,358,842 – 17,358,843) between the potential splice acceptor site and the first complete codon. Hence, this splice acceptor site is in phase 2.

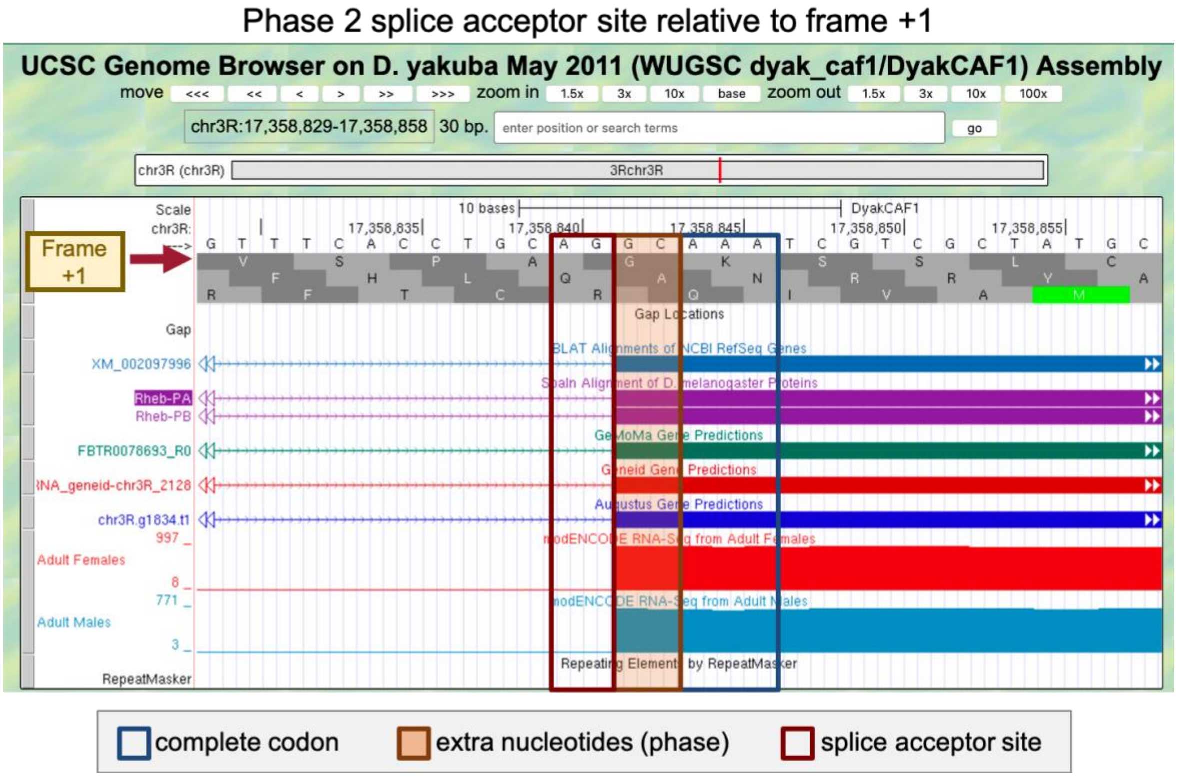

**Figure 41 This splice acceptor site (at 17,358,840-17,358,841) is supported by multiple lines of evidence. Since CDS-2 is in frame +1 and the first complete codon (AAA which codes for K) is located at 17,358,844-17,358,846, there are two nucleotides (GC at 17,358,842-17,358,843) between the splice acceptor site and the first complete codon. Hence, this splice acceptor site is in phase 2.**

Since the splice donor site for CDS-1 is in phase 1 (relative to frame +3), the splice acceptor site for CDS-2 must be in phase 2 (relative to frame +1) in order to maintain the open reading frame after the intron has been removed. The extra nucleotides near the splice sites (i.e., G + GC) will form an additional amino acid (Glycine/G) after splicing (Figure 42). Collectively, our analysis suggests that CDS-1 ends at 17,358,714 with a phase 1 splice donor site and CDS-2 begins at 17,358,842 with a phase 2 splice acceptor site.

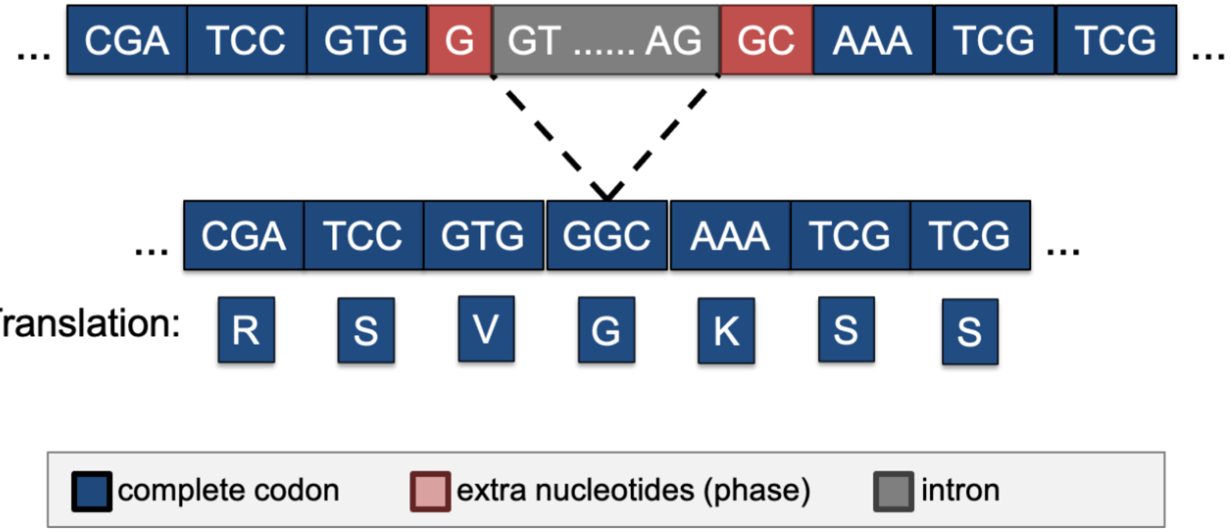

**Figure 42 The phase 1 donor site (G) of CDS-1 combines with the phase 2 acceptor site (GC) of CDS-2 to form the codon GGC, which codes for a Glycine (G).**

The same annotation strategy can be used to determine the phases for the remaining splice donor and splice acceptor sites between CDS-2 and CDS-3, CDS-3 and CDS-4, and CDS-4 and CDS-5, respectively (Figure 43).

| CDS | Frame | Splice Acceptor Phase | Splice Donor Phase |
|---|---|---|---|
| 1 | +3 | | 1 |
| 2 | +1 | 2 | 1 |
| 3 | +2 | 2 | 2 |
| 4 | +1 | 1 | 0 |
| 5 | +3 | 0 | |

**Figure 43 Summary of the phases of each splice donor and acceptor site in Rheb-RA.**

#### Part 6.4: Use spliced RNA-Seq reads to verify the coordinates for the first intron

Since RNA-Seq reads are derived primarily from processed mRNAs (where the introns have been removed), the subset of RNA-Seq reads that span multiple exons (i.e., spliced RNA-Seq reads) can provide us with additional evidence for a splice junction. When we map a spliced RNA-Seq read against the genome, part of the spliced read will map to one CDS while the rest of the read will map to another CDS. The region between these two alignment blocks would correspond to the intron. Splice site prediction tools such as *TopHat* or *regtools* can recognize this distinct mapping pattern of spliced RNA-Seq reads in order to infer the possible locations of the splice junction.

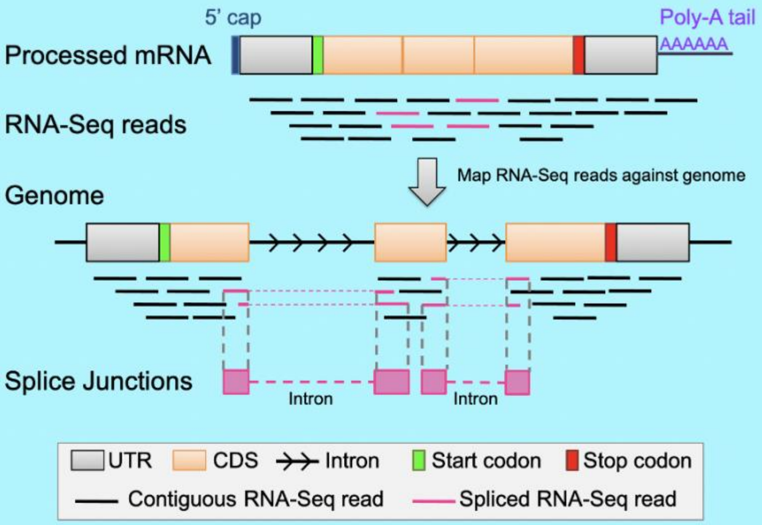

**See Understanding Eukaryotic Genes Module 4 to further review splice junction predictions.**

1. Under the “RNA Seq Tracks,” change the “Splice Junctions” track to “pack.”
   - Note: The “Splice Junctions” track identifies the splice junctions predicted by the *regtools* program.
2. Click on the “refresh” button.
3. To examine the region surrounding Intron-1 (i.e., intron between CDS-1 and CDS-2), enter “**chr3R:17,358,715-17,358,841**” into the “enter position or search term” text box.
   - Note: We found these coordinates in Part 6.3.
4. Click on the “go” button.
5. Zoom out 3x to examine the 381 bp surrounding this position (Figure 44).

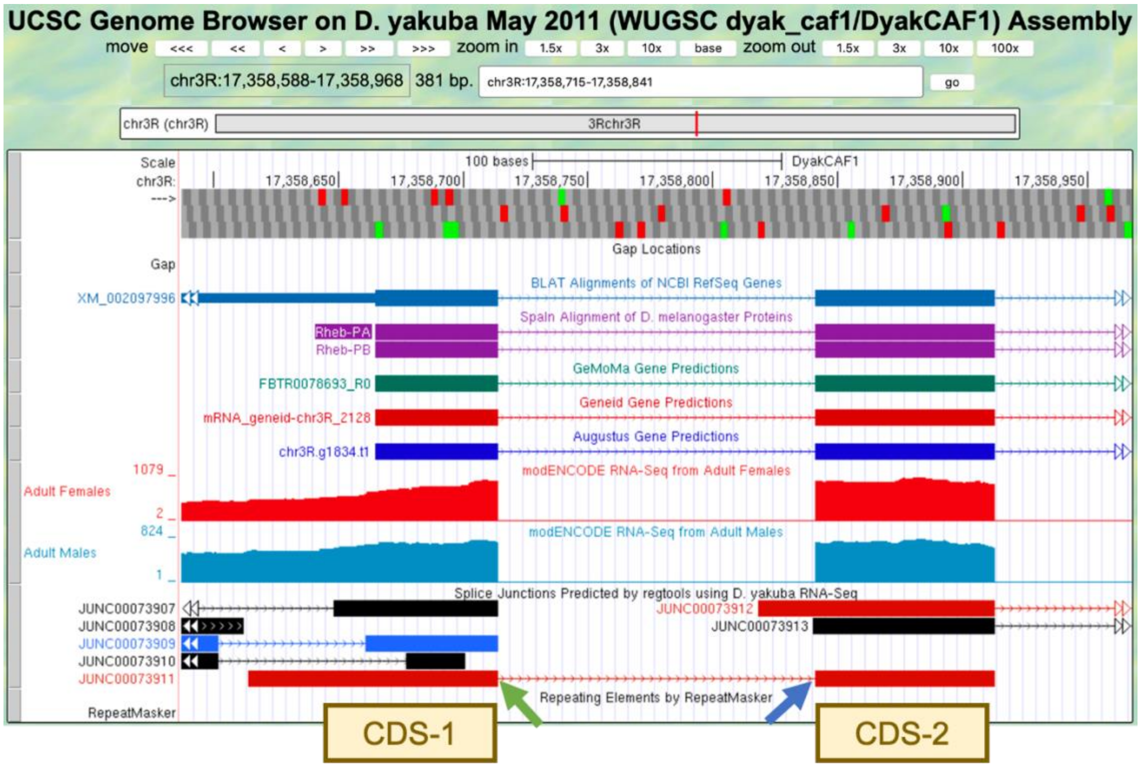

**Figure 44 The splice junction JUNC00073911 predicted by *regtools* connects CDS-1 with CDS-2. The predicted splice donor and acceptor sites are indicated by the green and blue arrows, respectively.**

There is only one splice junction predicted in this region.

6. To examine the splice donor site predicted by the splice junction JUNC00073911, zoom into the region surrounding the beginning of the intron predicted by this junction (Figure 44, green arrow).
7. To examine the splice acceptor site predicted by the splice junction JUNC00073911, zoom into the region surrounding the end of the intron predicted by this junction (Figure 44, blue arrow).

The splice junction between the phase 1 donor site of CDS-1 and the phase 2 acceptor site of CDS-2 is supported by the *BLAT* and *Spaln* alignments, the *GeMoMa*, *Geneid*, and *Augustus* gene predictions, and the RNA-Seq data from samples of adult females and males.

8. We can gather additional evidence to support this splice junction by clicking on "JUNC00073911" and then examining the "Score" field (Figure 45).

The score tells us how many spliced RNA-Seq reads there are that support a predicted splice junction. Since JUNC00073911 has a score of 1694, that splice junction prediction is supported by 1,694 spliced RNA-Seq reads. Splice junction predictions are color-coded based on the number of spliced RNA-Seq reads that support the junction (i.e., their scores). Based on the color-coded table in the "Description" section, JUNC00073911 will be red in the Genome Browser image since greater than 1,000 spliced RNA-Seq reads support the feature (Figure 45).

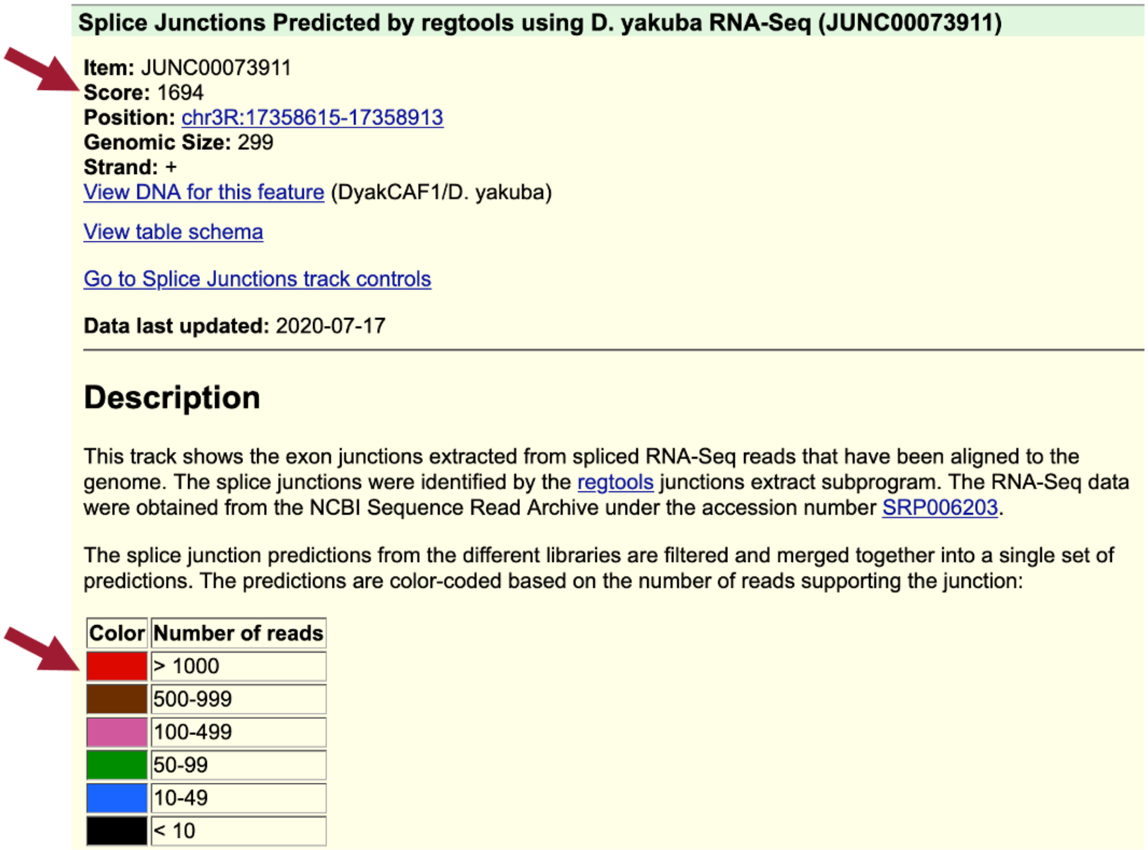

**Splice Junctions Predicted by regtools using D. yakuba RNA-Seq (JUNC00073911)**

**Item:** JUNC00073911
**Score:** 1694
**Position:** chr3R:17358615-17358913
**Genomic Size:** 299
**Strand:** +
View DNA for this feature (DyakCAF1/D. yakuba)

View table schema

Go to Splice Junctions track controls

**Data last updated:** 2020-07-17

**Description**

This track shows the exon junctions extracted from spliced RNA-Seq reads that have been aligned to the genome. The splice junctions were identified by the regtools junctions extract subprogram. The RNA-Seq data were obtained from the NCBI Sequence Read Archive under the accession number SRP006203.

The splice junction predictions from the different libraries are filtered and merged together into a single set of predictions. The predictions are color-coded based on the number of reads supporting the junction:

| Color | Number of reads |
|---|---|
| | > 1000 |
| | 500-999 |
| | 100-499 |
| | 50-99 |
| | 10-49 |
| | < 10 |

**Figure 45 The splice junction JUNC00073911 has a score of 1694 indicating that it is supported by 1,694 spliced RNA-Seq reads; therefore, this feature will be red in the Genome Browser image.**

9. Click on the back button of the web browser to return to the Genome Browser image.

Examination of the features in the “Splice Junctions Predicted by *regtools* using *D. yakuba* RNA-Seq” track shows that the splice junction prediction JUNC00073911 is consistent with our splice donor site for CDS-1 at 17,358,715 – 17,358,716 and our splice acceptor site for CDS-2 at 17,358,840-17,358,841 that we annotated in Part 6.3.

#### Part 6.5: Use splice junction predictions to verify the coordinates for the second intron

The same annotation strategy can be used to determine the coordinates for Intron-2 between CDS-2 and CDS-3.

1. To examine the region surrounding Intron-2, enter “**chr3R:17,358,912-17,359,013**” into the “enter position or search term” text box.
    - Note: These coordinates can be found in the table in Figure 35.
2. Click on the “go” button.
3. Zoom out 3x to examine the 306 bp surrounding this position (Figure 46).

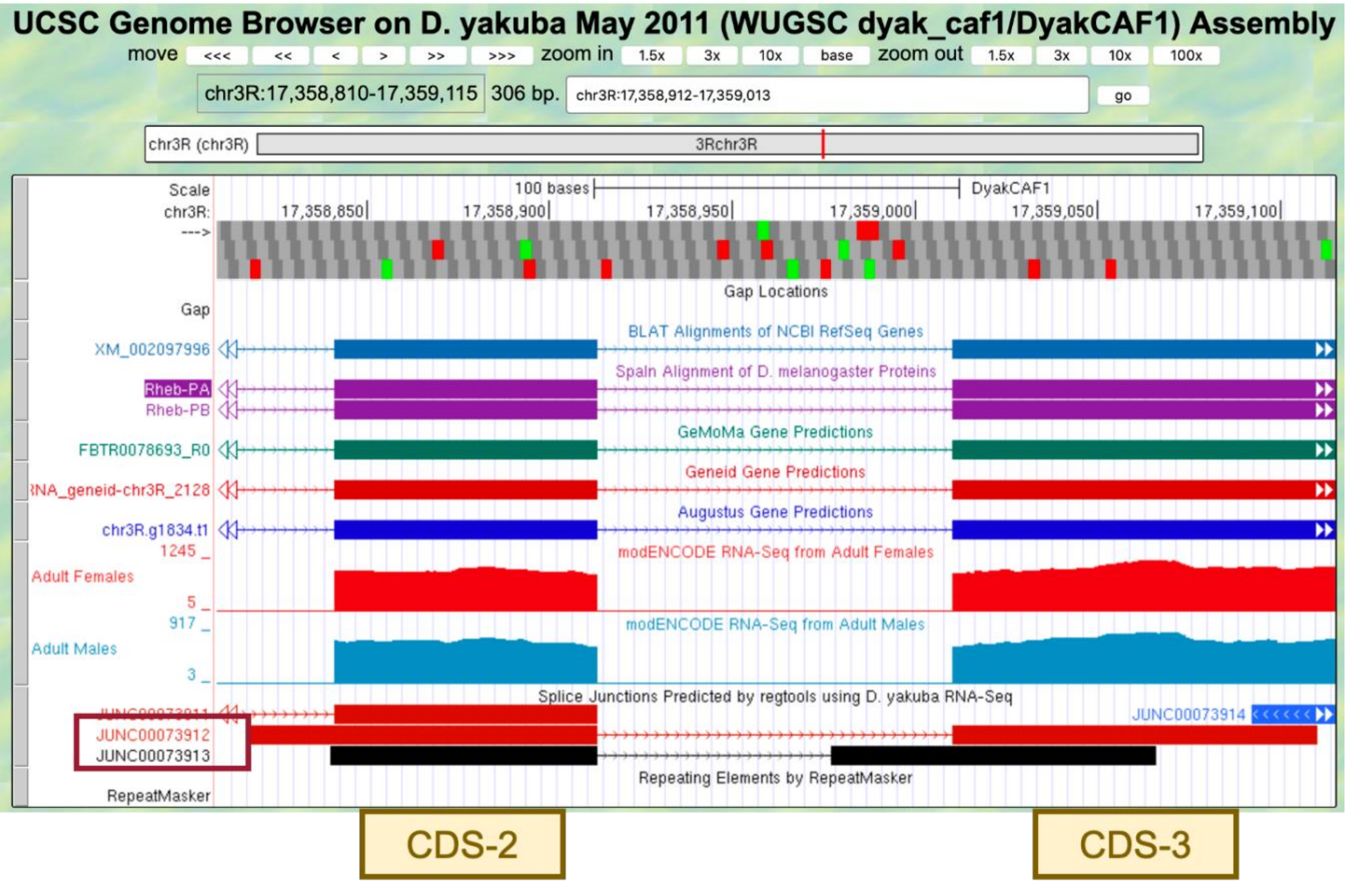

**Figure 46 The splice junctions JUNC00073912 and JUNC00073913 predicted by *regtools* connect CDS-2 with CDS-3.**

There are two splice junctions predicted in this region, JUNC00073912 and JUNC00073913.

The *tblastn* alignment for CDS-2 ends at 17,358,912 (Figure 35). The potential splice donor site at 17,358,914-17,358,915 for CDS-2 is supported by the splice junction predictions JUNC00073912 and JUNC00073913, the *BLAT* and *Spaln* alignments, the *GeMoMa*, *Geneid*, and *Augustus* gene predictions, and the RNA-Seq data from samples of adult females and males. There is one nucleotide (A at 17,358,913) between the last complete codon (AAC) and the potential splice donor site. Hence, this splice donor site is in phase 1 relative to frame +1.

The *tblastn* alignment for CDS-3 spans from 17,359,013-17,359,216 in frame +2 (Figure 35). The potential splice acceptor site at 17,359,009-17,359,010 for CDS-3 is supported by the splice junction prediction JUNC00073912, the *BLAT* and *Spaln* alignments, the *GeMoMa*, *Geneid*, and *Augustus* gene predictions, and the RNA-Seq data from samples of adult females and males. Note that this potential splice acceptor site is not supported by JUNC00073913. There are two nucleotides (CC at 17,359,011-17,359,012) between the first complete codon (TTC) and the potential splice acceptor site. Hence, this splice acceptor site is in phase 2 relative to frame +2. This phase 2 splice acceptor site is compatible with the phase 1 splice donor site for CDS-2.

4. Click on the “JUNC00073912” feature to determine the number of spliced RNA-Seq reads that support this splice junction prediction (Figure 47).
5. Click on the back button of the web browser to return to the Genome Browser image.

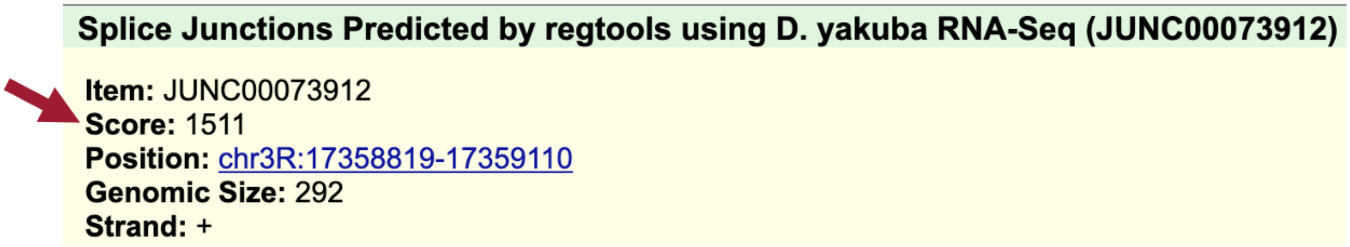
**Splice Junctions Predicted by regtools using D. yakuba RNA-Seq (JUNC00073912)**

**Item:** JUNC00073912
**Score:** 1511
**Position:** chr3R:17358819-17359110
**Genomic Size:** 292
**Strand:** +

**Figure 47 The score for JUNC00073912 shows that this junction is supported by 1,511 spliced RNA-Seq reads.**

In addition to the splice junction JUNC00073912, which supports the proposed splice acceptor site for CDS-3 at 17,359,009-17,359,010, there is another splice junction which suggests a different splice acceptor site (JUNC00073913) (Figure 48).

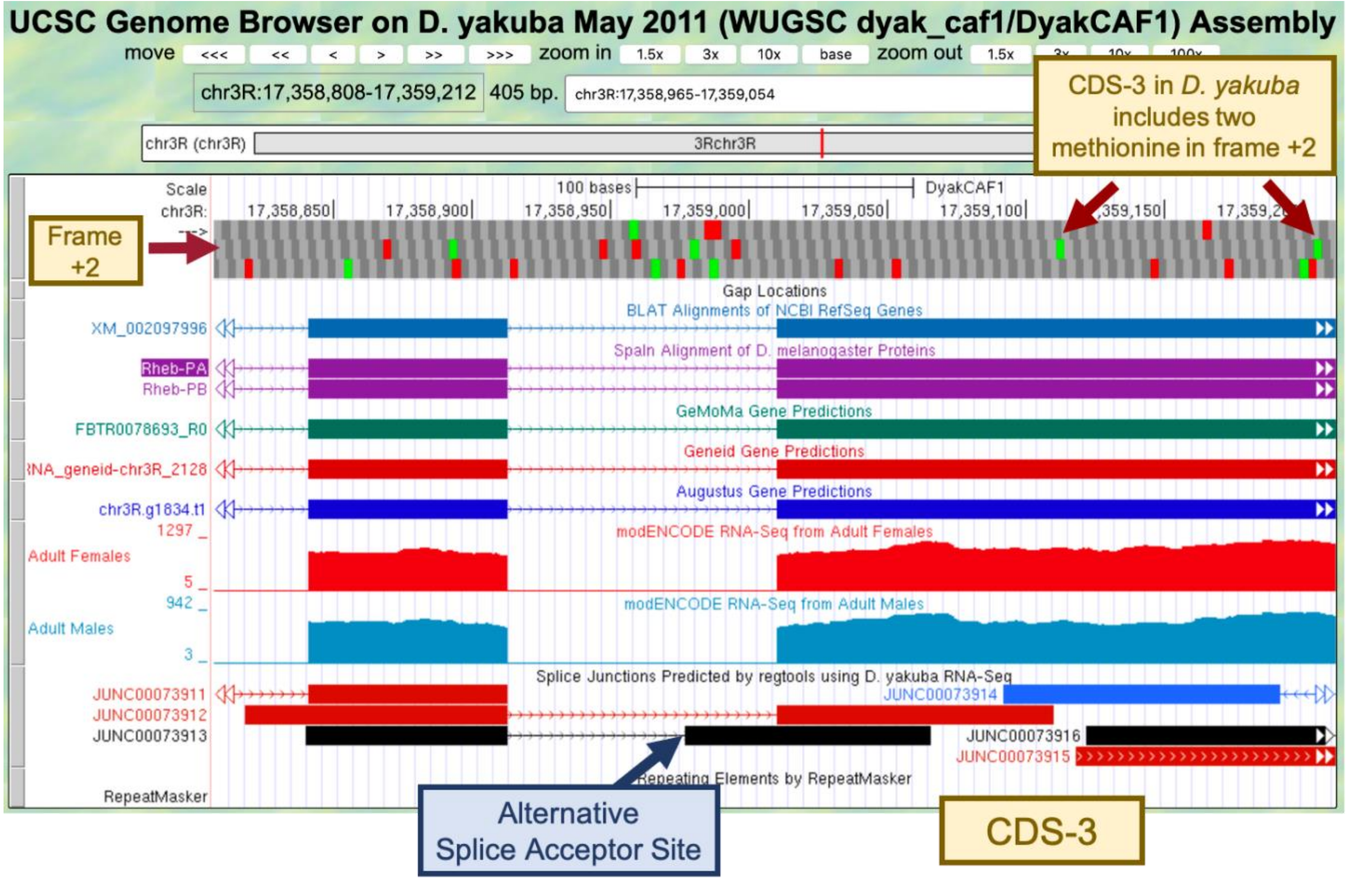

**Figure 48 Splice junction prediction JUNC00073913 proposes an alternative splice acceptor site at 17,358,976-17,358,977. Since CDS-3 in *D. yakuba* includes two methionine in frame +2, JUNC00073913 could indicate the presence of a novel isoform of *Rheb* in *D. yakuba* that is not found in *D. melanogaster*.**

CDS-3 in *D. yakuba* includes two methionine in frame +2 (at 17,359,112-17,359,114 and 17,359,205-17,359,207). Hence, the splice junction JUNC00073913 could indicate the presence of a novel isoform of *Rheb* in *D. yakuba* where the translation start site is located within CDS-3, and the region from the end of the splice junction (i.e., 17,358,978) to the start codon (e.g., 17,359,111) would be part of the 5' UTR.

In order to assess whether there is sufficient evidence to propose a novel isoform of *Rheb* in *D. yakuba*, we will need to determine the number of spliced RNA-Seq reads that support the splice junction JUNC00073913.

6. Click on the splice junction "JUNC00073913" and then examine the "Score" field (Figure 49).
7. Click on the back button of the web browser to return to the Genome Browser image.

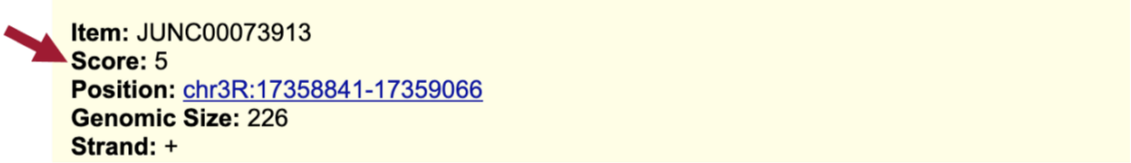

**Figure 49 The score for JUNC00073913 shows that this junction is only supported by 5 spliced RNA-Seq reads.**

Since the splice junction JUNC00073913 is only weakly supported by the available RNA-Seq data, there is insufficient evidence to postulate a novel isoform of *Rheb* in *D. yakuba* based on this splice junction prediction. A score filter can be applied to the "Splice Junctions" evidence track to hide the splice junction predictions that are supported by a small number of spliced RNA-Seq reads.

8. Scroll down to the "RNA Seq Tracks" section and then click on the "Splice Junctions" link.
9. Change the "Show only items with score at or above" to "10."
10. Click on the "Submit" button (Figure 50).

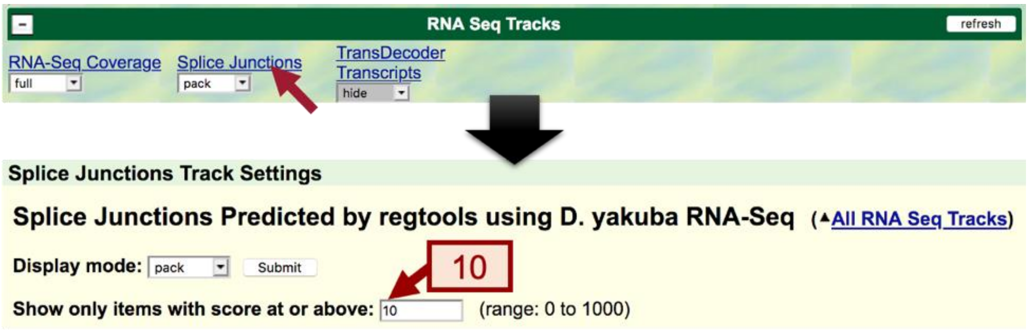

**Figure 50 Configure the Splice Junctions track to only display splice junctions that are supported by at least 10 spliced RNA-Seq reads.**

11. To examine the intron between CDS-2 and CDS-3 after applying the filter, enter "**chr3R:17,358,914-17,359,010**" into the "enter position or search terms."
12. Click on the "go" button.
13. Zoom out 1.5x.

After applying to filter, we no longer see splice junction prediction JUNC00073913 in the Genome Browser image.

#### Part 6.6: Use splice junction predictions to verify the coordinates for the third intron

The *tblastn* alignment for CDS-3 spans 17,359,013-17,359,216 in frame +2 (Figure 35).

1. To examine the region surrounding the splice donor site for CDS-3, enter "**chr3R:17,359,216**" into the "enter position or search terms" text box.
2. Click on the "go" button.
3. Zoom out 3x and another 10x (Figure 51).

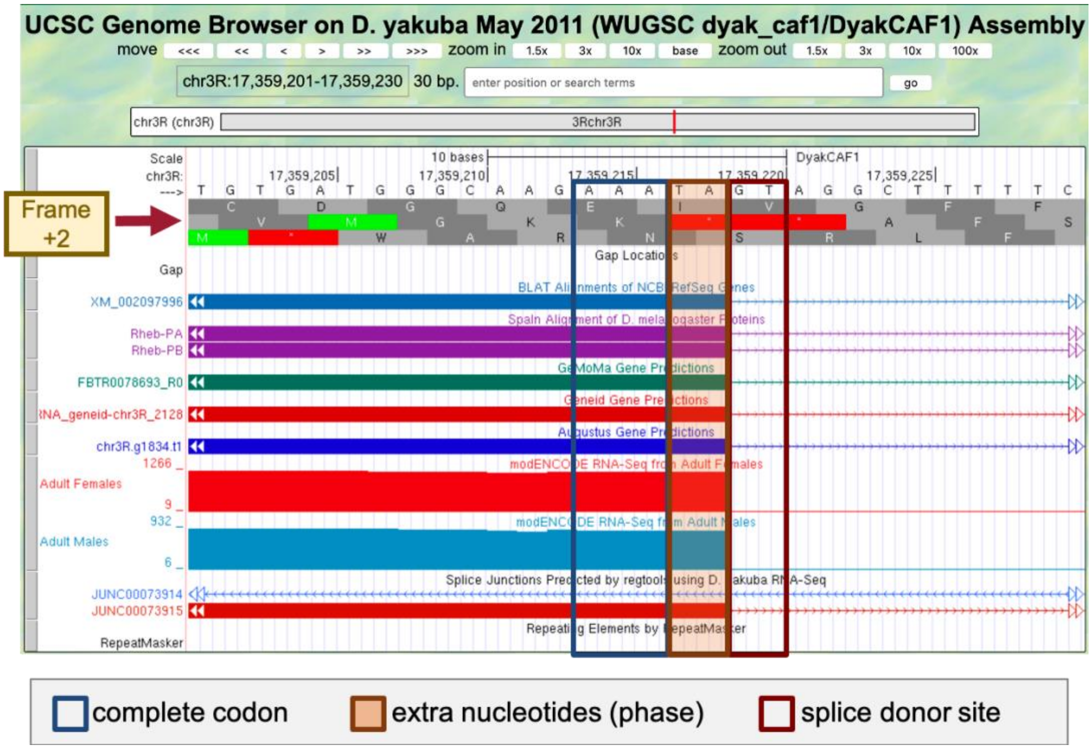

**Figure 51 The potential splice donor site at 17,359,219-17,359,220 for CDS-3 is supported by the *BLAT* and *Spaln* alignments, the *GeMoMa*, *Geneid*, and *Augustus* gene predictions, the RNA-Seq data from samples of adult females and males, and the splice junction prediction JUNC00073915. There are two nucleotides (TA at 17,359,217-17,359,218) between the last complete codon (AAA) and the potential splice donor site. Hence, this splice donor site is in phase 2 relative to frame +2.**

4. To examine the genomic region surrounding the start of the *tblastn* alignment to CDS-4 (Figure 35), enter "**chr3R:17,359,279**" into the "enter position or search terms" text box.
5. Click on the "go" button.
6. Zoom out 3x and another 10x (Figure 52).

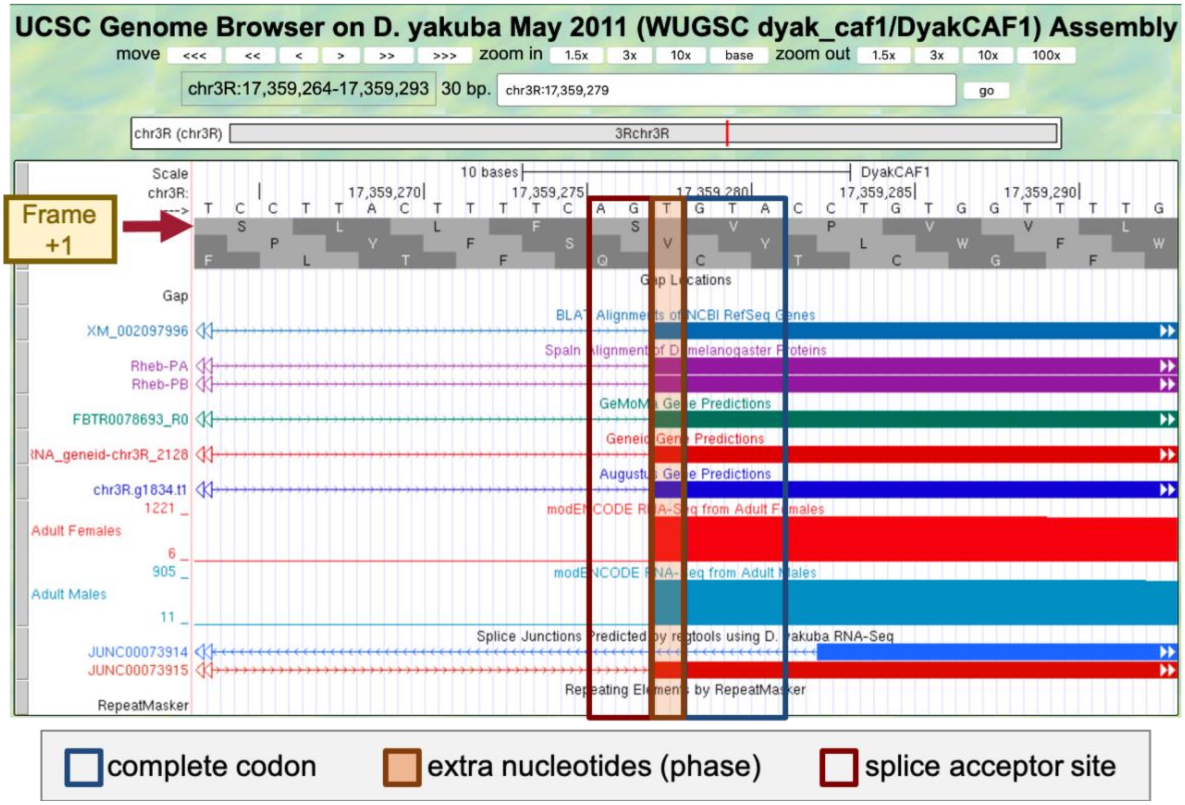

**Figure 52 The potential splice acceptor site at 17,359,276-17,359,277 for CDS-4 is supported by the *BLAT* and *Spaln* alignments, the *GeMoMa*, *Geneid*, and *Augustus* gene predictions, the RNA-Seq data from samples of adult females and males, and the splice junction prediction JUNC00073915. There is one nucleotide (T at 17,359,278) between the first complete codon (GTA) and the potential splice acceptor site. Hence, this splice acceptor site is in phase 1 relative to frame +1. This phase 1 splice acceptor site is compatible with the phase 2 splice donor site for CDS-3.**

The genomic region also contains an additional splice junction prediction JUNC00073914. Since this feature is in blue, this junction is supported by 10–49 spliced RNA-Seq reads. By contrast, the splice junction prediction JUNC00073915 is in red, indicating that the splice junction is supported by more than 1,000 spliced RNA-Seq reads (Figure 53).

7. Click on the “JUNC00073914” feature to learn more about this splice junction prediction.

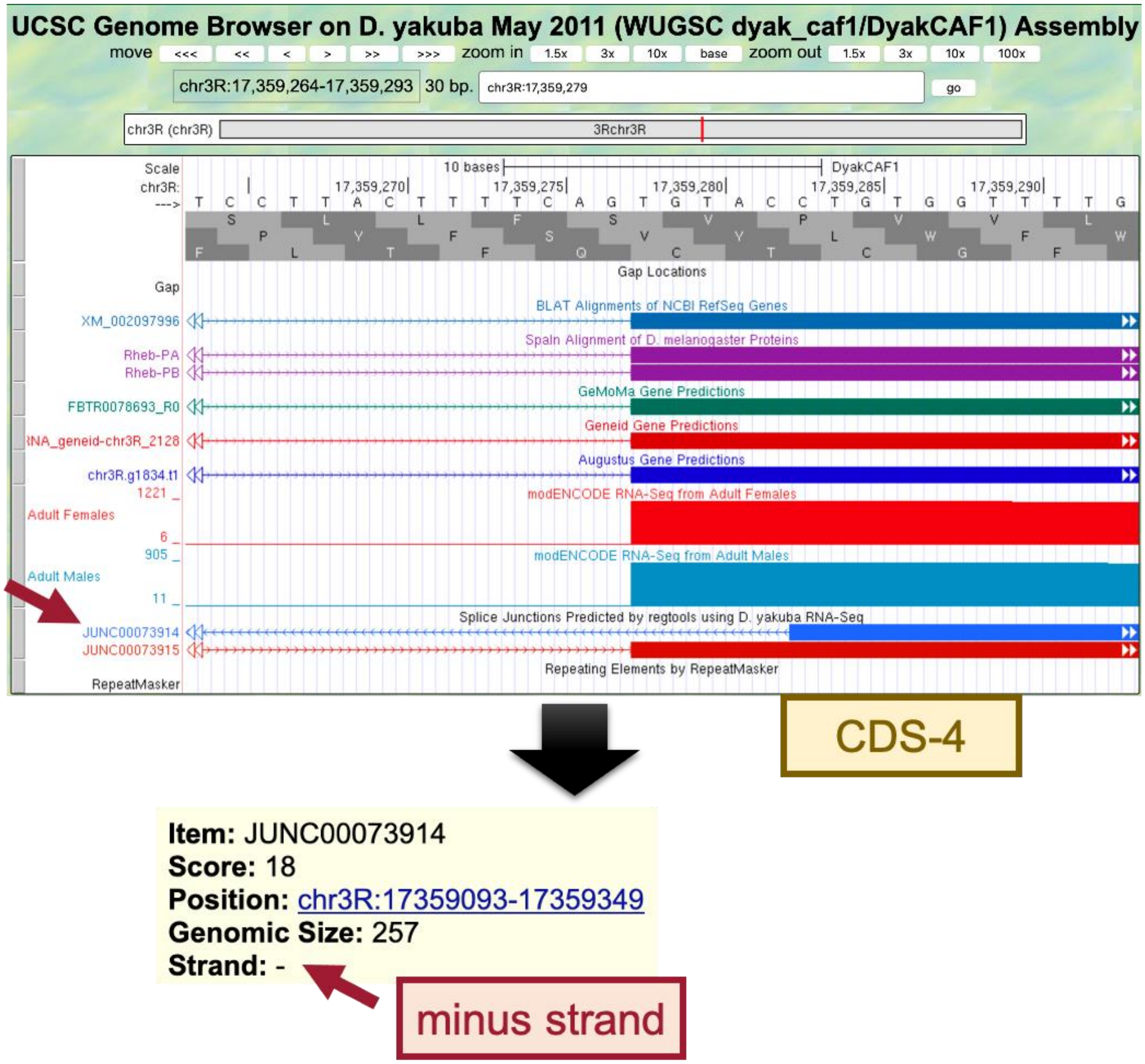

**Figure 53 The splice junction prediction JUNC00073914 is supported by 18 spliced RNA-Seq reads. However, the splice junction is in the minus strand whereas the *Rheb* gene is on the plus strand relative to *D. yakuba* chr3R.**

Because the *Rheb* ortholog is in the plus strand relative to *D. yakuba* chr3R and the splice junction JUNC00073914 is in the minus strand, this splice junction is unlikely to be an intron for the *Rheb* gene (Figure 53). This splice junction could be caused by RNA-Seq reads that have been misplaced, or it could have been derived from a non-coding RNA that overlaps with *Rheb*.

Based on the low score (18) of the JUNC00073914 splice junction and it being in the minus strand, the splice acceptor site for CDS-4 is assigned to 17,359,276-17,359,277.

#### Part 6.7: Use splice junction predictions to verify the change in donor site sequence for the fourth intron

Go back to the web browser tab with the Gene Record Finder record for the *D. melanogaster Rheb* gene.

1. Open a new tab and navigate to the Gene Record Finder.
2. Enter "**Rheb**" into the text box.
3. Click on the "Find Record" button.

The "Introns with Non-canonical Splice Sites" panel shows that the intron with the FlyBase ID "intron_Rheb:6_Rheb:7" has a GC splice donor in *D. melanogaster*. The FlyBase ID for an intron begins with the prefix "intron_", followed by the names of the two transcribed exons that flanked the intron. Hence, the FlyBase ID "intron_Rheb:6_Rheb:7" indicates that the splice donor site between the transcribed exons Rheb:6 and Rheb:7 has the non-canonical sequence GC (Figure 54).

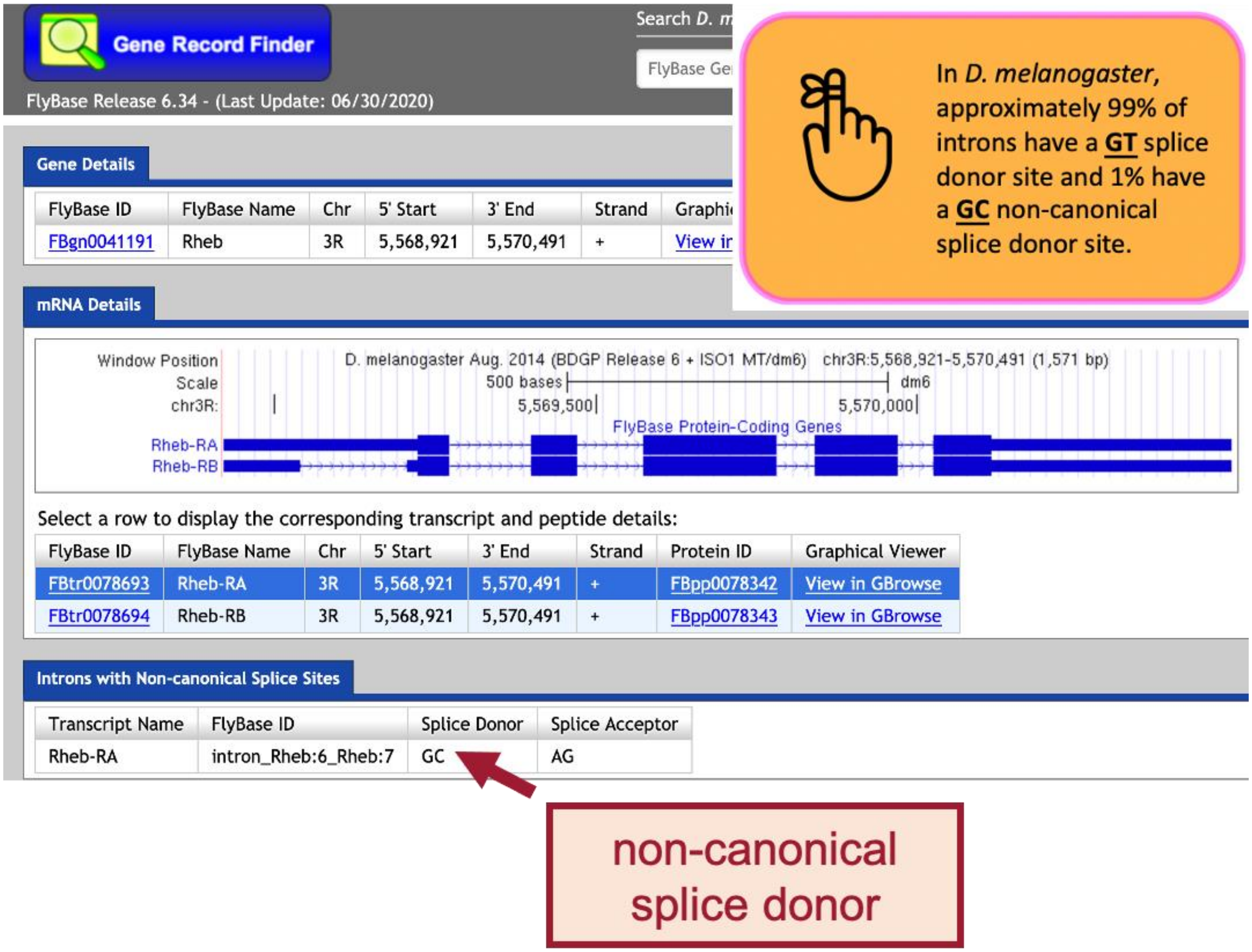

**Figure 54 The "Introns with Non-canonical Splice Sites" panel of the Gene Record Finder shows that the intron between the transcribed exons Rheb:6 and Rheb:7 begins with a non-canonical GC splice donor site in *D. melanogaster*.**

4. To determine the IDs of the CDS's that overlap with the transcribed exons Rheb:6 and Rheb:7, click on the Genome Browser image in the "mRNA Details" panel of the Gene Record Finder.

Comparison of the "FlyBase Transcribed Exons" and "FlyBase Coding Exons" tracks in the Genome Browser for *D. melanogaster* shows that the coding exon CDS-4 (Rheb:4_9855_1) overlaps with the transcribed exon Rheb:6, and coding exon CDS-5 (Rheb:5_9855_0) overlaps with the transcribed exon Rheb:7.

5. Zoom in to the end of CDS-4 to verify that it has a non-canonical GC donor site (Figure 55).

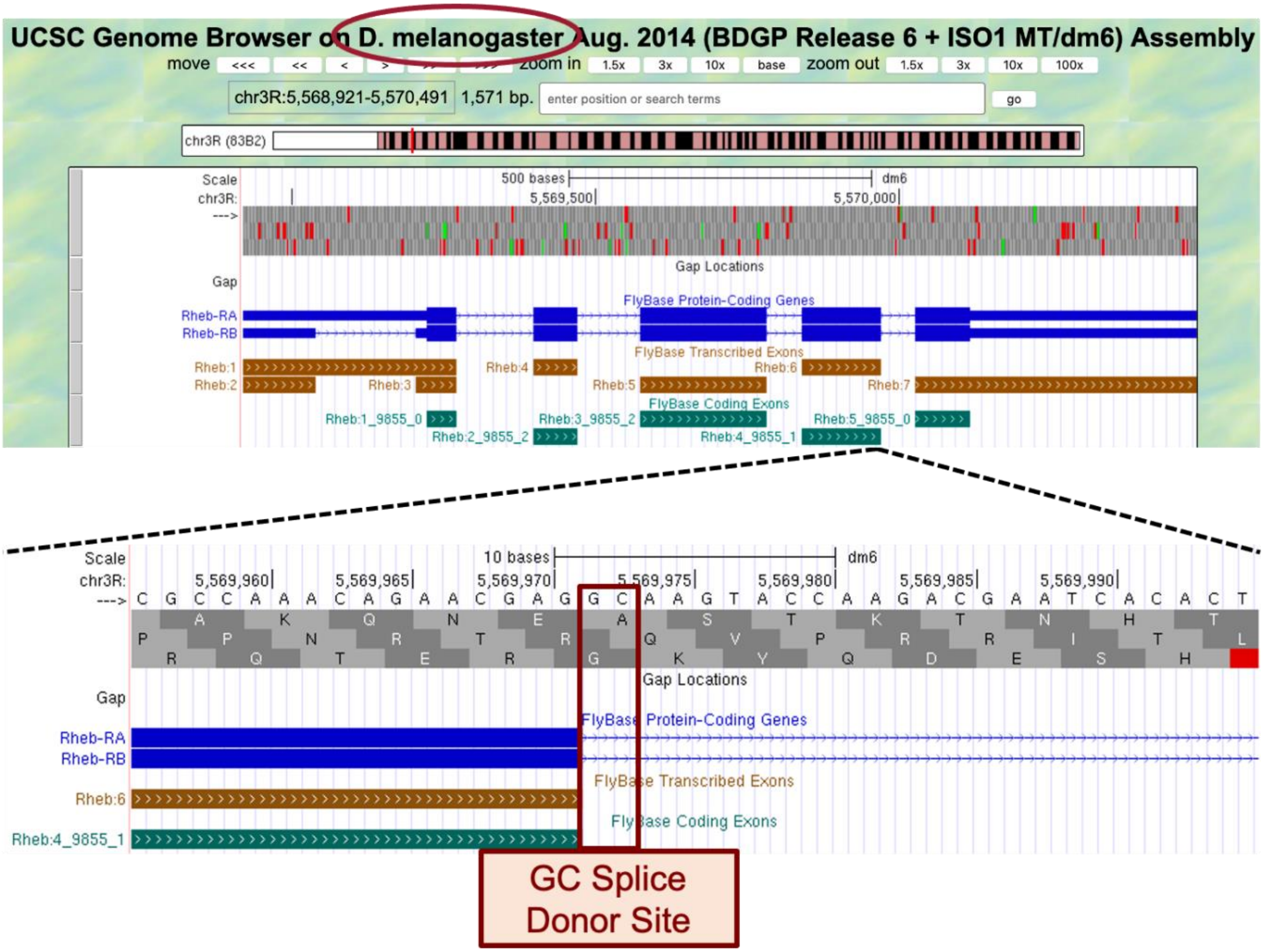

**Figure 55 Examination of the region surrounding the end of CDS-4 in *D. melanogaster* shows that it has a non-canonical GC splice donor site.**

Consequently, if the splice sites are conserved between *D. melanogaster* and *D. yakuba*, then the splice donor site for CDS-4 would also use a non-canonical GC donor site in *D. yakuba*.

6. Go back to the web browser tab with the Genome Browser for *D. yakuba*.

The *tblastn* alignment for CDS-4 ends at 17,359,407 in frame +1 (Figure 35).

7. To examine the genomic region surrounding the end of the *tblastn* alignment to CDS-4, enter "**chr3R:17,359,407**" into the "enter position or search terms" text box.
8. Click on the "go" button.
9. Zoom out 3x and another 10x (Figure 56).

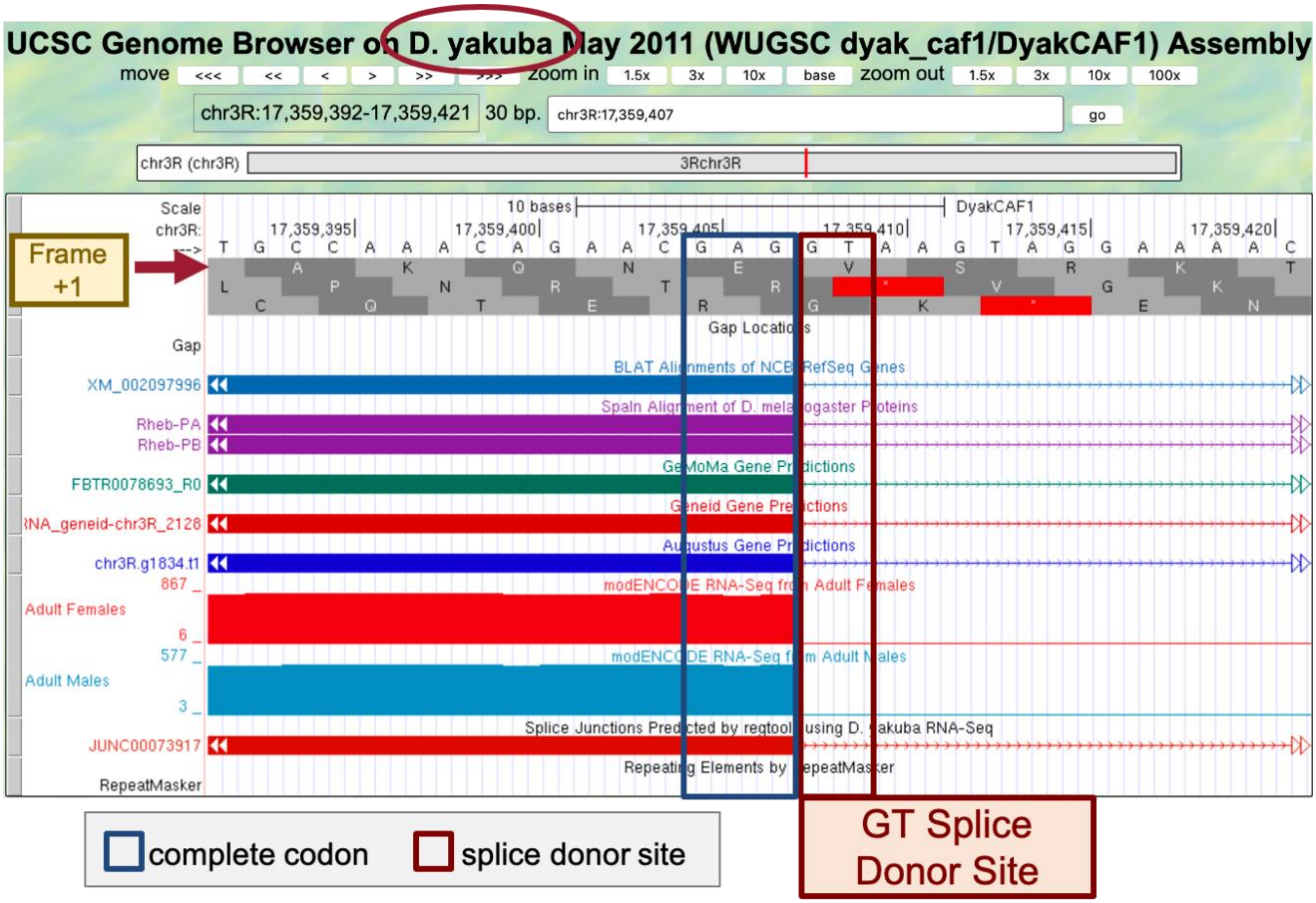

**Figure 56 The potential GT splice donor site at 17,359,408-17,359,409 for CDS-4 is supported by the *BLAT* and *Spaln* alignments, the *GeMoMa*, *Geneid*, and *Augustus* gene predictions, the RNA-Seq data from samples of adult females and males, and the splice junction prediction JUNC00073917; therefore, the available evidence indicates that the splice donor site for CDS-4 in *D. yakuba* has a canonical GT donor site (instead of the non-canonical GC donor site as seen in *D. melanogaster*). There are no nucleotides between the last complete codon (GAG) and the potential splice donor site. Hence, this splice donor site is in phase 0 relative to frame +1.**

The *tblastn* alignment for CDS-5 spans 17,359,470 – 17,359,559 in frame +3 (Figure 35).

10. To examine the genomic region surrounding the start of the *tblastn* alignment to CDS-5, enter "**chr3R:17,359,470**" into the "enter position or search terms" text box.
11. Click on the "go" button.
12. Zoom out 3x and another 10x (Figure 57).

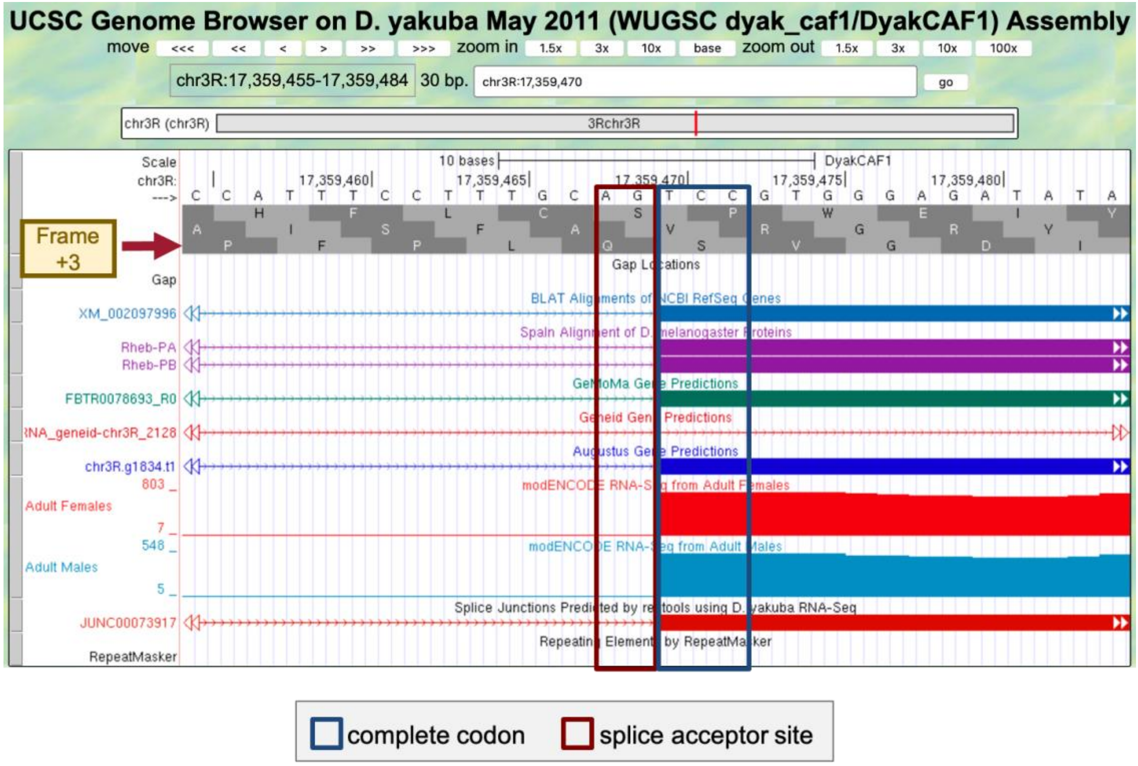

**Figure 57 The potential splice acceptor site at 17,359,468-17,359,469 for CDS-5 is supported by the *BLAT* and *Spaln* alignments, the *GeMoMa* and *Augustus* gene predictions, the RNA-Seq data from samples of adult females and males, and the splice junction prediction JUNC00073917. The *Geneid* prediction mRNA_*geneid*-chr3R_2128 is inconsistent with the potential splice acceptor site. However, per the previous analysis described in Part 6.2, this *Geneid* prediction might have combined two adjacent genes (*Rheb* and *CRMP*) into a single prediction. There are no nucleotides between the potential splice acceptor site and the first complete codon (TCC). Hence, this splice acceptor site is in phase 0 relative to frame +3. This phase 0 splice acceptor site is compatible with the phase 0 splice donor site for CDS-4.**

Taken together, the analysis described above would produce the CDS coordinates for Rheb-PA in *D. yakuba* chr3R summarized below (Figure 58):

**Gene Model for Rheb-PA in *D. yakuba***

| CDS | Frame | Splice Acceptor Phase | Coordinates | | Splice Donor Phase |
|---|---|---|---|---|---|
| | | | Start | End | |
| 1 | +3 | | 17,358,666 | 17,358,714 | 1 |
| 2 | +1 | 2 | 17,358,842 | 17,358,913 | 1 |
| 3 | +2 | 2 | 17,359,011 | 17,359,218 | 2 |
| 4 | +1 | 1 | 17,359,278 | 17,359,407 | 0 |
| 5 | +3 | 0 | 17,359,470 | 17,359,556 | |

**Figure 58 Summary of the five CDS's in Rheb-PA. Stop codon is located at 17,359,557-17,359,559.**

### Part 7: Verify and submit gene model

Now that we've completed the annotation of Rheb-PA in *D. yakuba*, we need to verify our proposed gene model and prepare the corresponding sequence files for submission.

#### Part 7.1: Verify gene model of Rheb-PA

Our analysis of the CDS-by-CDS *tblastn* alignments and the evidence tracks on the Genome Browser allow us to precisely define the start and end positions of each of the five coding exons (CDS's) of Rheb-PA. To verify that our proposed gene model satisfies the basic biological constraints (e.g., begins with a start codon, has compatible splice sites, ends with a stop codon), we will check our gene model coordinates using the Gene Model Checker.

1. Open a new web browser tab and navigate to the Gene Model Checker (Figure 59).
2. In the "Project Details" section:
   - Species Name: select "D. yakuba"
   - Genome Assembly: select "May 2011 (WUGSC dyak_caf1/DyakCAF1)"
   - Scaffold Name: enter "**chr3R**"
3. In the "Ortholog Details" section:
   - Ortholog in D. melanogaster: enter "**Rheb-PA**"
4. In the "Model Details" section:
   - Errors in Consensus Sequence?: select "No"
   - Coding Exon Coordinates: enter a comma-delimited list of coordinates for the five CDS's "**17358666-17358714, 17358842-17358913, 17359011-17359218, 17359278-17359407, 17359470-17359556**"
   - Annotated Untranslated Regions?: select "No"
   - Orientation of Gene Relative to Query Sequence: select "Plus" since Rheb-PA is on the positive strand relative to chr3R scaffold
   - Completeness of Gene Model Translation: select "Complete"
   - Stop Codon Coordinates: click within the textbox and the coordinates will automatically populate
5. Click on the "Verify Gene Model" button to run the Gene Model Checker.

> ⚠ Note that the coordinates for the "Coding Exon Coordinates" field **do not include the stop codon**. We will enter the stop codon coordinates separately in the "Stop Codon Coordinates" field.

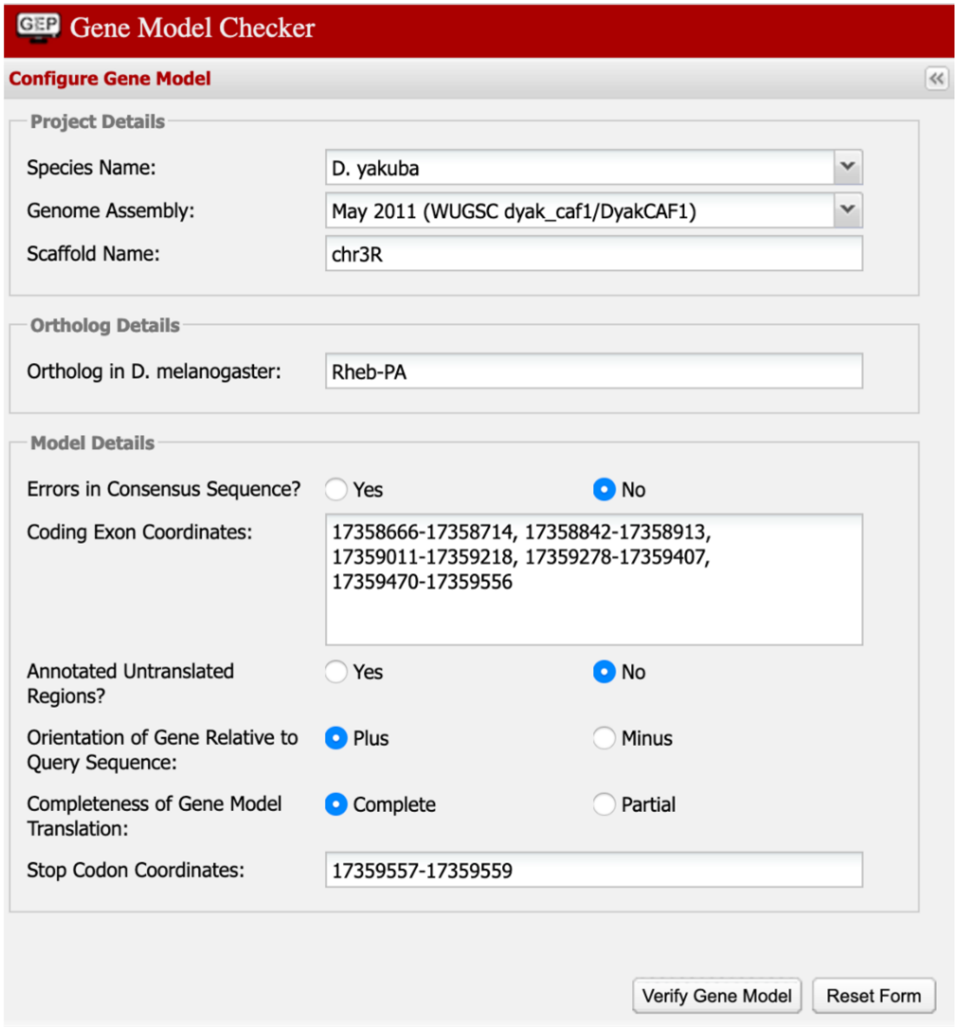

**Figure 59 Verify the D. *yakuba* gene model for Rheb-PA using the Gene Model Checker.**

Once the analysis is complete, the right panel of the Gene Model Checker contains the results. The "Checklist" tab enumerates the list of criteria that have been checked by the Gene Model Checker (Figure 60). For example, the Gene Model Checker verifies that our proposed gene model begins with a start codon and ends with a stop codon. It also verifies that the splice junctions contain the standard splice donor and acceptor sites. Some of the items on the checklist have been skipped because they do not apply to a complete gene (e.g., Acceptor for CDS 1).

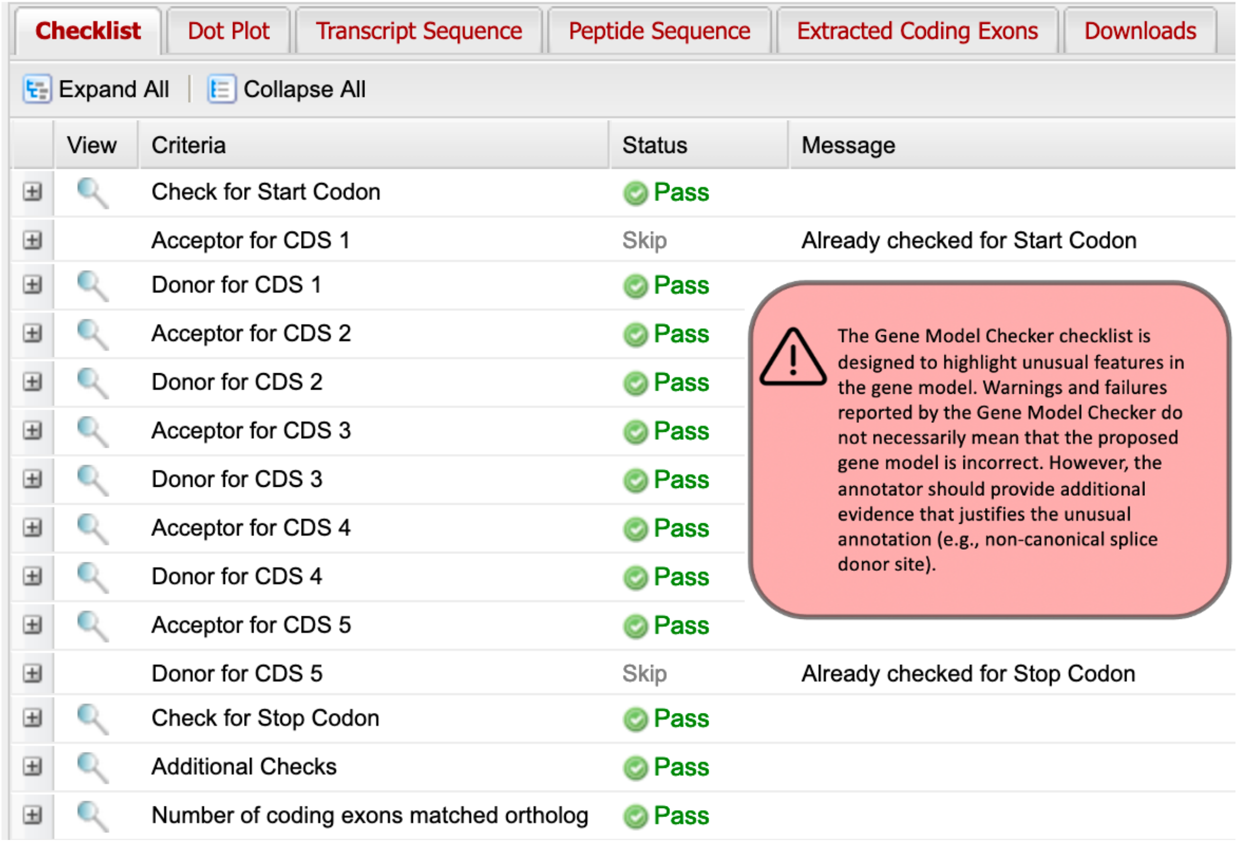

**Figure 60 The Gene Model Checker checklist shows that the proposed gene model for Rheb-PA satisfies the biological constraints of most protein-coding genes (e.g., canonical start codon, stop codon, splice sites).**

In addition to verifying the basic gene structure, the Gene Model Checker also compares the proposed gene model against the putative *D. melanogaster* ortholog using a protein alignment and a dot plot.

6. Click on the "**Dot Plot**" tab to examine the dot plot between the *D. melanogaster* protein (x-axis) and the protein sequence for the submitted model in *D. yakuba* (y-axis) (Figure 61).

The alternating color boxes in the dot plot correspond to the different CDS's in the two sequences. Dots in the dot plot correspond to regions of similarity between the *D. melanogaster* protein and the submitted *D. yakuba* gene model. If the submitted sequence is identical to the *D. melanogaster* ortholog, then the dot plot will show a straight diagonal line with a slope of 1. Changes in the size of the submitted model compared to the *D. melanogaster* ortholog will alter the slope of this line.

In this case, the dot plot shows that the five CDS's of Rheb-PA in *D. melanogaster* and *D. yakuba* have similar lengths (compare the length shown on the x-axis to the length shown on the y-axis for each CDS). However, within a small region of CDS-4, the dot plot did not detect sequence similarity between the submitted model for *D. yakuba* and the *D. melanogaster* ortholog (Figure 61).

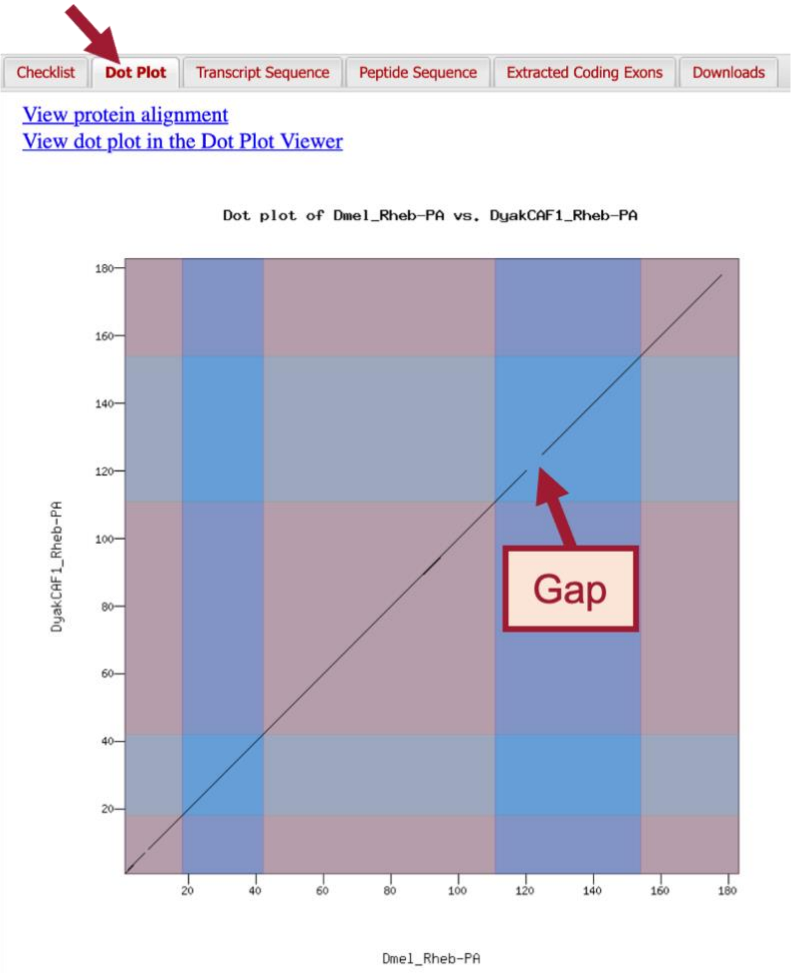

**Figure 61 The dot plot comparing the *D. melanogaster* Rheb-PA (x-axis) with the submitted gene model in *D. yakuba* (y-axis) shows that the main differences between the two protein sequences are located in CDS-4 and CDS-5.**

To further investigate the dot plot, we will examine the protein alignment between the two sequences.

7. Click on the "View protein alignment" link above the dot plot (Figure 62).

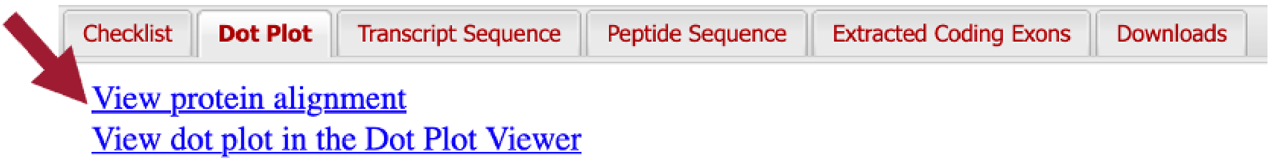

**Figure 62 Click on the "View protein alignment" link under the "Dot Plot" tab.**

The alignment shows the comparison of the *D. melanogaster* protein against the conceptual translation for the submitted *D. yakuba* gene model. Similar to the dot plot, the alternating colors correspond to the different CDS's (Figure 63).

The protein alignment shows that the five CDS's have high levels of sequence similarity between the *D. melanogaster* ortholog and the *D. yakuba* gene model. The symbols in the match line denote the level of similarity ("*****" indicates conserved amino acids, "**:**" denotes amino acids with highly similar chemical properties). Hence, the tiny gap in the dot plot within CDS-1 can be attributed to similar, but not identical, amino acids near its center.

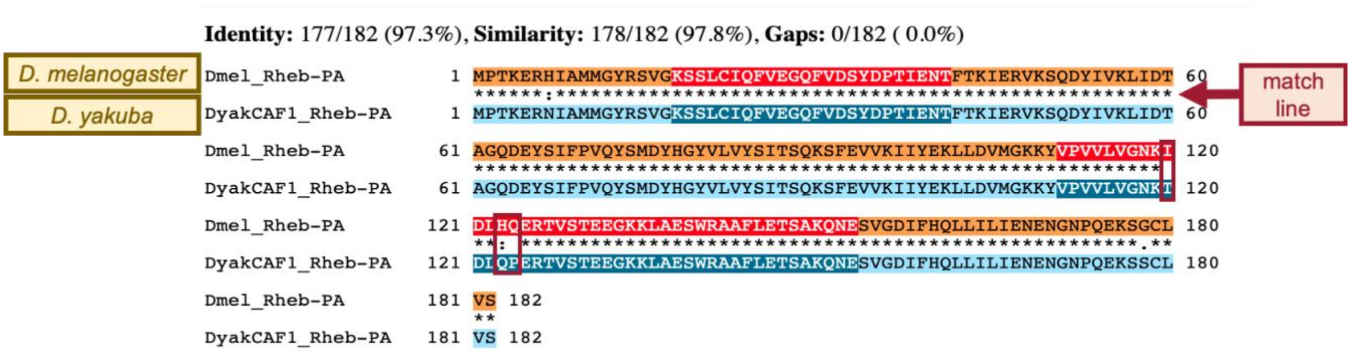

**Figure 63 The protein alignment between *D. melanogaster* Rheb-PA versus the submitted gene model in *D. yakuba* shows that the tiny gap within CDS-4 in the dot plot can be attributed to three amino acid residues that differ between these two species (red boxes).**

The protein alignment between the *D. melanogaster* Rheb-PA and the submitted gene model in *D. yakuba* shows that that the gap within CDS-4 in the dot plot can be attributed to differences in three amino acid residues. We need to verify the placement of CDS-4.

8. Click on the "Checklist" tab.
9. Click on the magnifying glass icon next to the "Acceptor for CDS 4" entry (Figure 64).
   - Note: A new window containing the Genome Browser will appear with our submitted gene model shown in the red "Custom Gene Model" track.

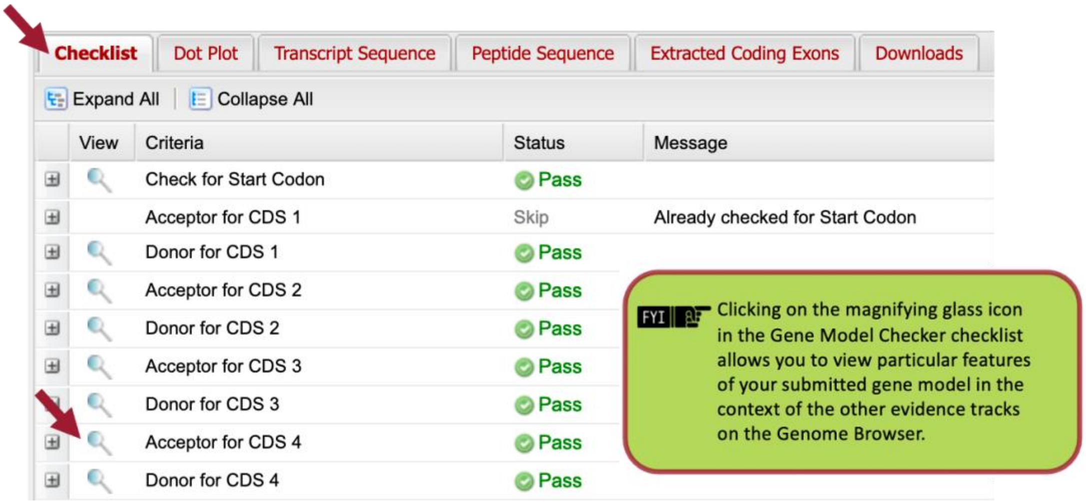

**Figure 64 To verify the placement of CDS-4, click on the magnifying glass icon next to the "Acceptor for CDS 4" entry.**

10. Zoom out 1.5x and another 3x so that we can examine the genomic region surrounding CDS-4 of Rheb-PA in frame +1.

The placement of CDS-4 in the Custom Gene Model for Rheb-PA is in congruence with the *BLAT* and *Spaln* alignments, the *GeMoMa*, *Geneid*, and *Augustus* gene predictions, and the RNA-Seq data from samples of adult females and males (Figure 65). The red arrows in Figure 65 demarcate the three amino acid residues that differ between CDS-4 in *D. yakuba* and *D. melanogaster*.

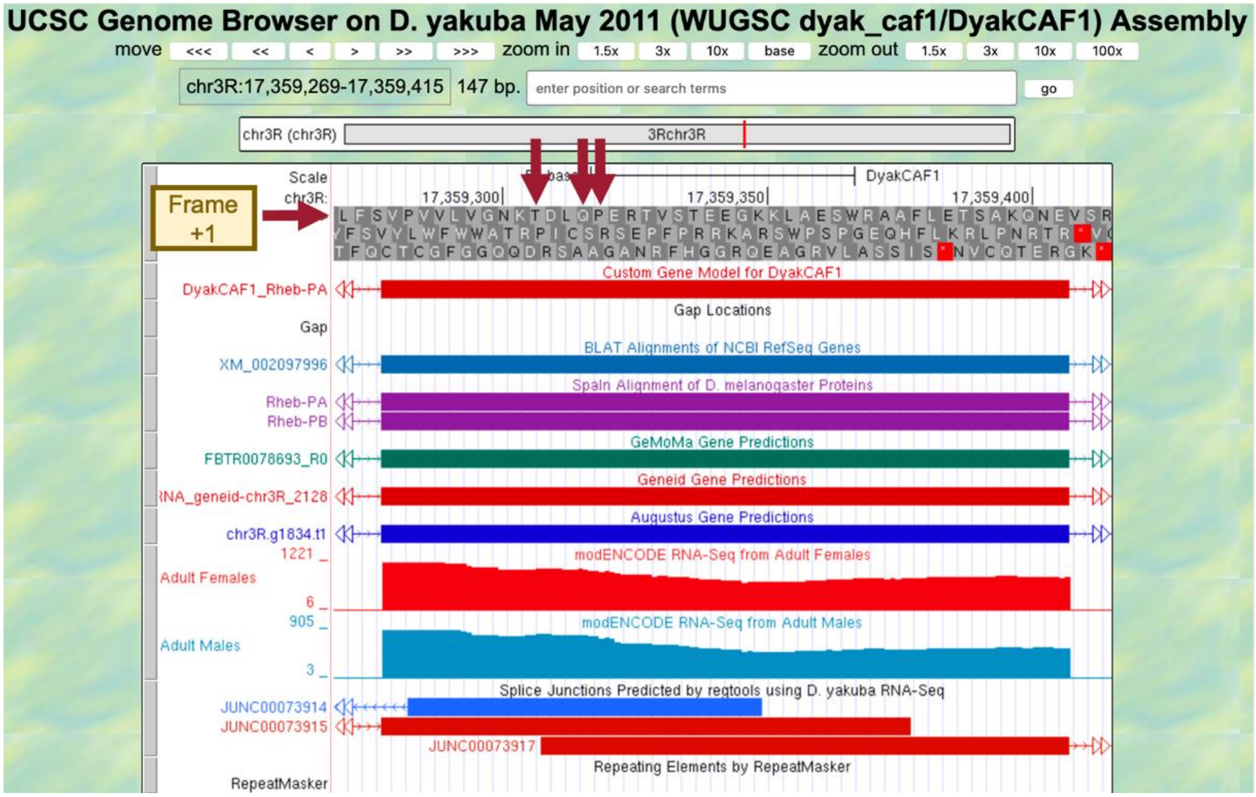

**Figure 65 The placement of CDS-4 in the Custom Gene Model for Rheb-PA is in congruence with multiple lines of evidence. The red arrows demarcate the three amino acid residues that differ between CDS-4 in *D. yakuba* and *D. melanogaster*.**

To verify our previous observations, we can view the submitted gene model within the context of the other evidence tracks in the Genome Browser.

11. Zoom out until you can see our entire submitted gene model for Rheb-PA in *D. yakuba* (Figure 66).

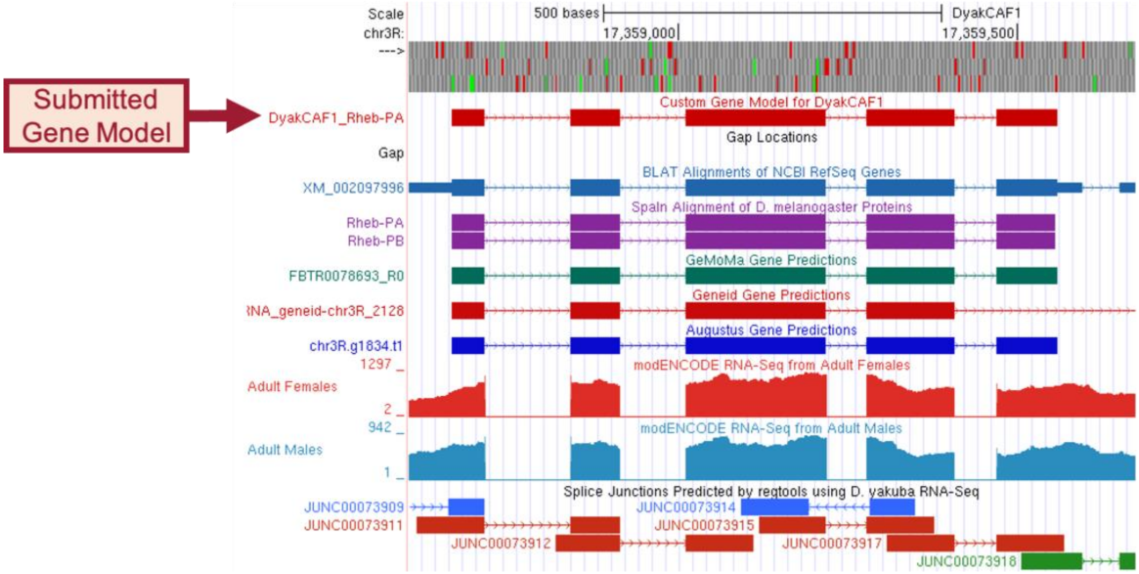

**Figure 66 Our submitted gene model for Rheb-PA in *D. yakuba* is shown under the track title "Custom Gene Model for DyakCAF1."**

#### Part 7.2 Download the files required for project submission

In addition to the Pathways Project Annotation Report Form (automatic download), you must prepare three additional data files to submit a project to the GEP – a General Feature Format File (GFF), a Transcript Sequence File (fasta), and a Peptide Sequence File (pep). The Gene Model Checker automatically creates these three files for a specific isoform (e.g., Rheb-PA) when you verify a gene model.

1. You can download these files by clicking on the "Downloads" tab and then clicking on each of the links to save each file to your computer (Figure 67).
   - Note: You don't have to rename these files, you'll only need to rename the merged file.

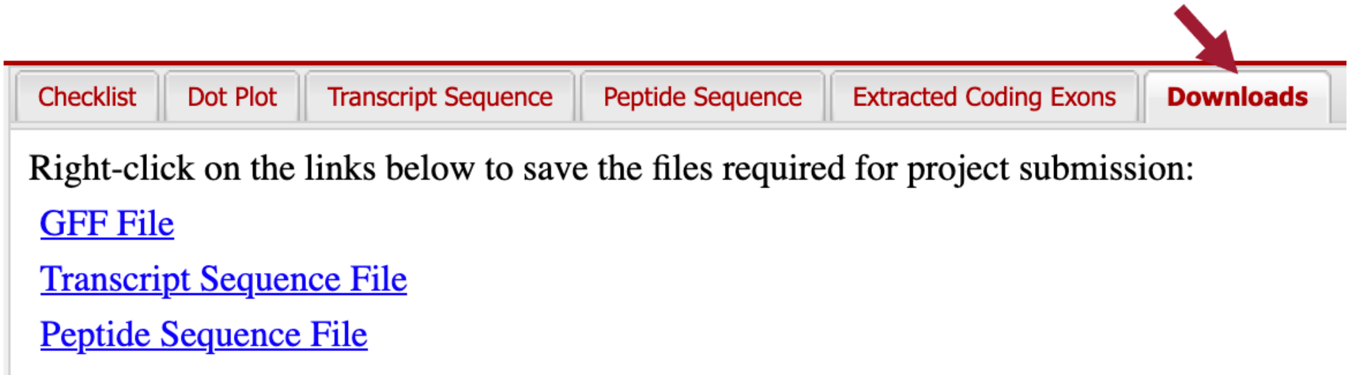

**Figure 67 In preparation for project submission, click on the "Downloads" tab and save the GFF, transcript sequence, and peptide sequence files for the gene model to your computer.**

Recall that *Rheb* in *D. yakuba* has two isoforms; the files we just downloaded were for the Rheb-PA isoform. Since the coding exons (CDS's) for both of our isoforms are identical, we only need to change the name of the ortholog in the Gene Model Checker and then verify our second isoform.

2. Change the name of the ortholog in the "Ortholog in D. melanogaster" field to "**Rheb-PB**" (Figure 68).
    - **Leave everything else the same as we had it for Rheb-PA.**
3. Click on "Verify Gene Model" to create a new set of files for Rheb-PB.

We can check that the Rheb-PB isoform was verified by looking at the axes titles under the "Dot Plot" tab.

4. Click on the "Downloads" tab and then click on each of the links to save each file to your computer.

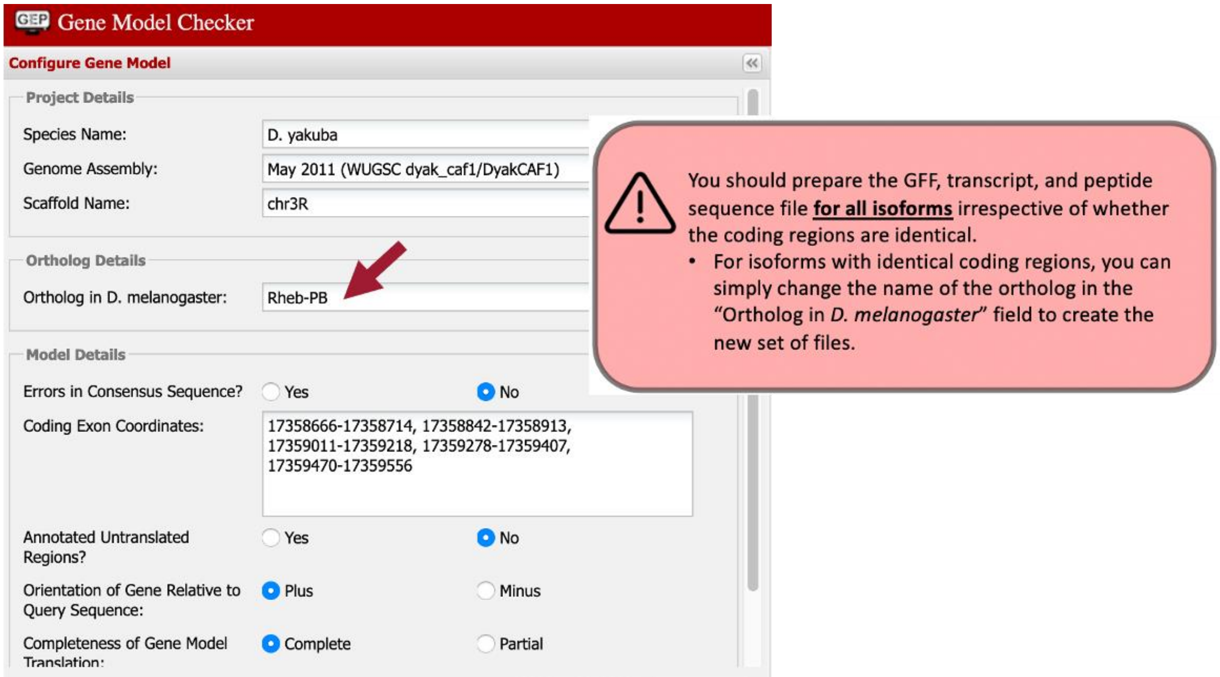

**Figure 68 Change the name of the ortholog in the "Ortholog in D. melanogaster" field to "Rheb-PB" and then click on "Verify Gene Model."**

Now we need to combine the GFF, transcript sequence, and peptide sequence files for all of our isoforms into a single file prior to project submission (we'll submit one merged file for each file type).

5. Open a new web browser tab and navigate to the [Annotation Files Merger.]

Let's merge our two isoform GFF files first.

6. Change the "File Type:" to "GFF Files (.gff)."
7. Drag the two GFF files we downloaded for Rheb-PA and Rheb-PB to the "Drag and drop the files you want to merge here" section.
8. Click on the "Merge Files" button (Figure 69).

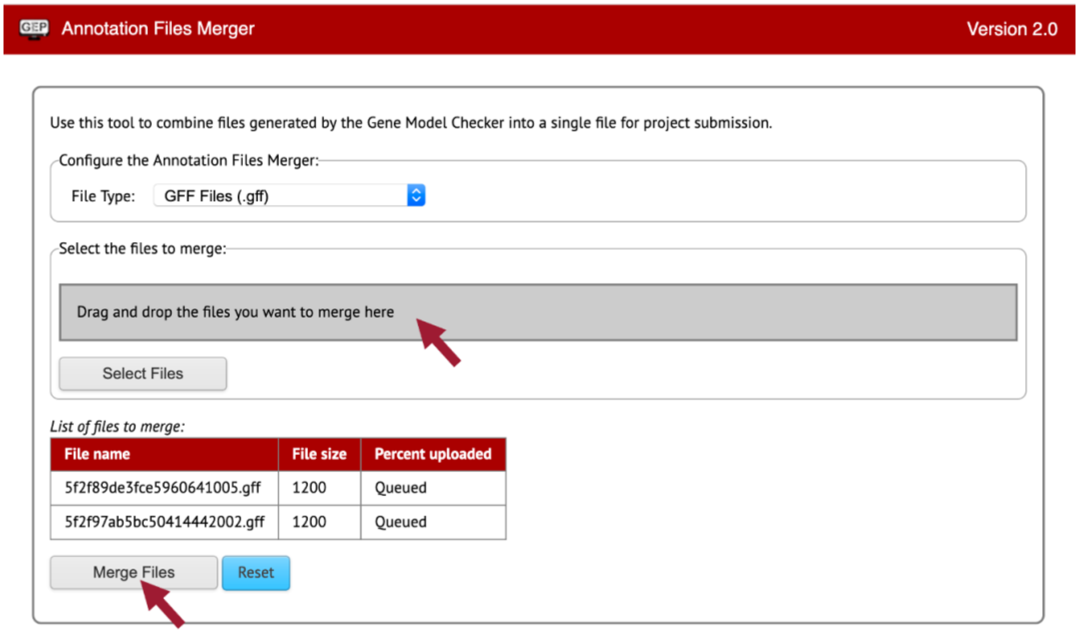

**Figure 69 Merge the GFF files for Rheb-PA and Rheb-PB.**

9. Download the merged GFF file by right-clicking (control click on macOS) on the "Merged File Link" (Figure 70).
10. Click on "Save Link As…"
11. Enter "**dyak_Rheb.gff**" as the file name.
12. Once you click on the "Save" button, the merged GFF file should download onto your computer.

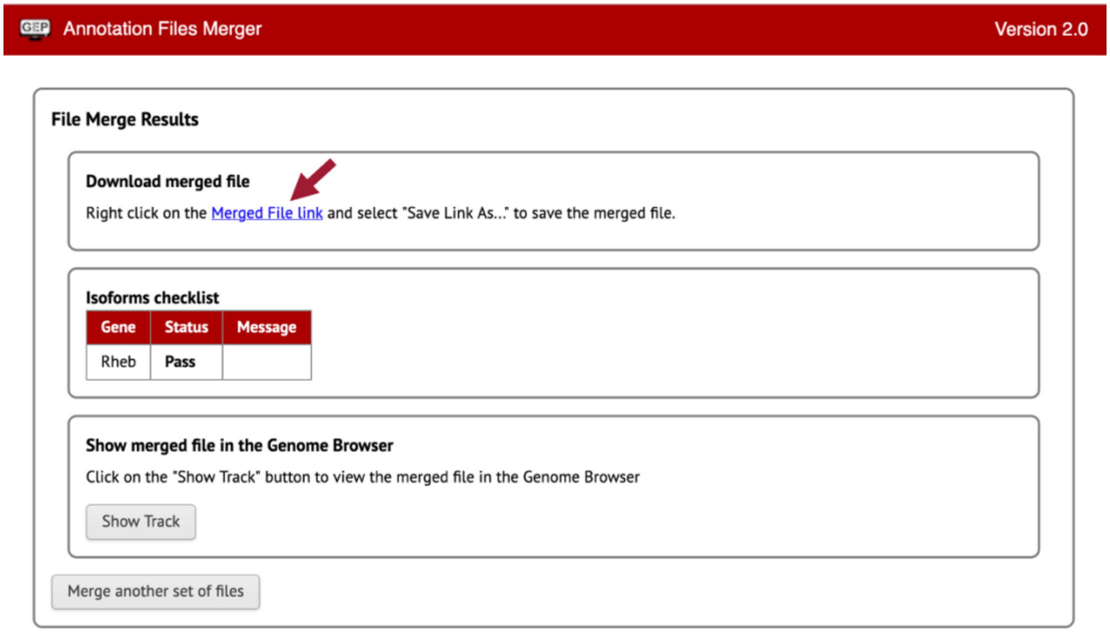

**Figure 70 Download the merged GFF files for Rheb-PA and Rheb-PB.**

Now we need to merge our two isoform Transcript Sequence Files (.fasta).

13. Click on the “Merge another set of files” button.
14. Change the “File Type:” to “Transcript Sequence Files (.fasta).”
15. Drag the two fasta files we downloaded for Rheb-PA and Rheb-PB to the “Drag and drop the files you want to merge here” section.
16. Repeat steps 8-10.
17. Enter “**dyak_Rheb.fasta**” as the file name.
18. Once you click on the “Save” button, the merged fasta file should download onto your computer.

Lastly, we need to merge our two isoform Peptide Sequence Files (.pep).

19. Click on the “Merge another set of files” button.
20. Change the “File Type:” to “Peptide Sequence Files (.pep).”
21. Drag the two pep files we downloaded for Rheb-PA and Rheb-PB to the “Drag and drop the files you want to merge here” section.
22. Repeat steps 8-10.
23. Enter “**dyak_Rheb.pep**” as the file name.
24. Once you click on the “Save” button, the merged pep file should download onto your computer.

### Appendix

#### A. Investigate the other *tblastn* alignments to *D. yakuba* chr3R

**General Information**

| | | | |
|---|---|---|---|
| **Symbol** | Dmel\Rheb-PA | **Species** | *D. melanogaster* |
| **Annotation Symbol** | CG1081-PA | **FlyBase ID** | FBpp0078342 |
| **Associated gene** | Dmel\Rheb | | |
| **Length (aa)** | 182 ← 182aa | **Theoretical pI** | 5.32 |
| **Predicted MW (kDa)** | 20.7 | | |

...

**Protein Domains**

**Protein Domains (via Pfam)**

Isoform displayed: Rheb-PA

Small_GTPase 182

Pfam protein domains

| InterPro name | classification | start | end |
|---|---|---|---|
| Small GTPase (Small_GTPase) | Family | 8 | 164 |

Small_GTPase domain: 8–164aa

**The "Protein Domains" section of the FlyBase Polypeptide Report for Rheb-PA (FBpp0078342) shows a conserved Small_GTPase domain at residues 8-164 aa of the protein.**

**Range 1: 5727469 to 5727792** GenBank Graphics ▼ Next Match ▲ Previous Match

| Score | Expect | Method | Identities | Positives | Gaps | Frame |
|---|---|---|---|---|---|---|
| 60.5 bits(145) | 2e-10 | Compositional matrix adjust. | 39/109(36%) | 57/109(52%) | 1/109(0%) | +1 |

```
Query  55       VKLIDTAGQDEYSIFPVQYSMDYHGYVLVLVYSITSQKSFEVVKIIYEKLLDVMGKKYVPVV  114
                 ++ DTAGQ+ Y         Y       G +LVY I      ++E V+      +L D     +    V ++
Sbjct  5727469  AQIWDTAGQERYRAITSAYYRGAVGALLVYDIAKHLTYENVERWLRELRDHADQNIV-IM  5727645

Query  115      LVGNKIDLHQERTVSTEEGKKLAESWRAAFLETSAKQNESVGDIFHQLL  163
                LVGNK DL    R+V T+E K  AE      +F+ETSA  + +V    F  +L
Sbjct  5727646  LVGNKSDLRHLRSVPTDEAKLFAERNGLSFIETSALDSTNVETAFQNIL  5727792
```

**Examination of the query coordinates of the additional *tblastn* alignments to chr3R shows that they overlap with the Small_GTPase conserved domain. For example, the region at 5,727,469-5,727,792 aligned with amino acid residues 55-163 of the Rheb-PA protein when it is translated on the positive strand in the first reading frame (+1).**

**Range 3: 9373062 to 9373502** GenBank Graphics ▼ Next Match ▲ Previous Match ▲ First Match

| Score | Expect | Method | Identities | Positives | Gaps | Frame |
|---|---|---|---|---|---|---|
| 58.9 bits(141) | 1e-09 | Compositional matrix adjust. | 45/148(30%) | 75/148(50%) | 25/148(16%) | -2 |

```
Query  8        IAMMGYRSVGKSSLCIQFVEGQFVDSYDPTIENTFTKIERVKSQDYIVKLIDTAGQDEY-  66
                I ++G    VGK+ + I + + +F + Y PT+ +         V  +DY + L DTAGQ++Y
Sbjct  9373502  ITVVGDGMVGKTCMLITYTQNEFPEEYVPTVFDNHACNIAVDDRDYNLTLWDTAGQEDYE  9373323

Query  67       SIFPVQY--------------------SMDYHGYVLVLVYSITSQKSFEVVKIIYEKLLDVM  106
                 + P+ Y                    +   + ++L YSI+S+ SFE VK  +    +
Sbjct  9373322  RLRPLSYPNVSTIALQRDDVIPKLESIAP*TNCFLLCYSISSRTSFENVKSKWWPEIRHF  9373143

Query  107      GKKYVPVVLVGNKIDL---HQERTVSTE  131
                   +VPVVLVG K+DL    + E+ V+T+
Sbjct  9373142  SA-HVPVVLVGTKLDLRIPNSEKFVTTQ  9373062
```

In-frame stop codon

**Examination of the *tblastn* alignment blocks also show three cases where the alignments contain in-frame stop codons (i.e., at 9,373,502-9,373,062, 9,658,240-9,658,575, and 26,681,803-26,681,411). The in-frame stop codons can likely be attributed to homologous overextension of the alignments into the introns.**
