## Supplement 3 for "Manual Annotation of Genes within *Drosophila* Species: the Genomics Education Partnership protocol"

### Key Steps in Ortholog Assignment

Step 1: Examine the genomic neighborhood surrounding the target gene (e.g., *Ilp2*) in the informant species (e.g., *D. melanogaster*) using the GEP's mirror of the UCSC Genome Browser.

- Determine the two genes upstream of the target gene (relative to the 5' end).
- Determine the two genes downstream of the target gene (relative to the 3' end).
- Identify any nested genes in the genomic neighborhood of the target gene.

Step 2: Identify the location of the putative ortholog of the target gene in the target species (e.g., *D. yakuba*) based on sequence similarity.

- Obtain the protein sequence for the target gene in the informant species from the *Gene Record Finder* (or from FlyBase)
  - For genes with multiple isoforms, use the isoform with the largest coding region in this analysis
- Perform a *tblastn* search of the protein sequence for the target gene from the informant species against the genome assembly of the target species
- Interpret the *tblastn* search result:
  - If the *tblastn* search reports no significant (E-value < 0.05) matches, change the *tblastn* search parameters to increase the sensitivity of the search (e.g., decrease the Word Size from 6 to 3, turn off the low complexity filter, increase the Expect threshold to 10).
  - If the *tblastn* search reports multiple matches, compare the E-value for the best match with the E-value for the second-best match. If these matches have similar E-values, use the Reciprocal Best Hit strategy (Ward N et al., 2014) to support the ortholog assignment.

Step 3: Examine the genomic neighborhood surrounding the target gene (e.g., *Ilp2*) in the target species (e.g., *D. yakuba*)

- Navigate to the genomic region surrounding the putative ortholog of the target gene in the target species using the GEP UCSC Genome Browser
- Obtain the predicted (RefSeq) protein sequence for the two genes upstream and the two genes downstream of the putative ortholog
  - Also, include any nested gene predictions that are found within the introns of the putative ortholog of the target gene or that the putative ortholog may be nested within in this analysis
- For each gene prediction, perform a *blastp* search of the predicted protein sequence in the target species against the database of annotated proteins in *D. melanogaster*.

    - Report the best *D. melanogaster* protein match and the second best match (based on E-value and percent identity) for the most upstream, nearest upstream, nearest downstream, and the most downstream genes in the target species.
- Compare the relative gene order and orientations of the genes surrounding the target gene in the informant species and in the target species (Figure 5).

Step 4: If local synteny is not conserved between the informant and target species, examine the genomic region surrounding the next best match in the *tblastn* search using the protocol in Step 3

- After evaluating all of the potential candidate regions from the *tblastn* search, define the approximate location of the ortholog of the target gene based on sequence similarity (E-value, percent identity) and conservation of local synteny.
