## Supplement 4 for "Manual Annotation of Genes within *Drosophila* Species: the Genomics Education Partnership protocol"

### Track Information for Tracks used in Coordinate Refinement

| Track Set | Track | Description |
|---|---|---|
| Genes and Gene Predictions Tracks | RefSeq genes | Transcripts from the NCBI RefSeq database of that species aligned against the assembly using *BLAT* |
| | D. mel Transcripts | *D. melanogaster* transcripts annotated by FlyBase aligned against the genome assembly using a translated *BLAT* search |
| | GeMoMa | *D. melanogaster* protein sequences from FlyBase aligned against each scaffold in the assembly (the predicted gene models were constructed using *GeMoMa*. |
| | Drosophila RefSeq Transcripts | *Drosophila* transcripts from the NCBI RefSeq database were aligned against the assembly using BLAT. |
| | Genscan | The predictions are based on transcriptional, translational and donor/acceptor splicing signals as well as the length and compositional distributions of exons, introns and intergenic regions. |
| | GeneID | Geneid is a program to predict genes in anonymous genomic sequences designed with a hierarchical structure. |
| | Augustus | The predictions are based on transcriptional, translational, and donor/acceptor splicing signals, as well as the length and compositional distributions of exons, introns and intergenic regions. |
| | SNAP | The predictions are based on transcriptional, translational, and donor/acceptor splicing signals, as well as the length and compositional distributions of exons, introns and intergenic regions. |
| | GlimmerHMM | A new gene finder based on a Generalized Hidden Markov Model (GHMM). The predictions are based on transcriptional, translational, and donor/acceptor splicing signals, as well as the length and compositional distributions of exons, introns and intergenic regions. |
| RNA-Seq Tracks | RNA-Seq Coverage | For embryos, females, males, and cumulative. |
| | Splice Junctions | Generated using regtools |
| | TransDecoder Transcripts | TransDecoder was run against the transcripts assembled by StringTie using default parameters. |
| Comparative Genomics | Drosophila Conservation (28 species) | measurements of evolutionary conservation using two methods (phastCons and phyloP) from the PHAST package |
