## Supplement 6 for "Manual Annotation of Genes within *Drosophila* Species: the Genomics Education Partnership protocol"

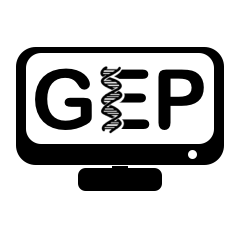
Pathways Project: Annotation Report

**Note:** You should also prepare the corresponding **GFF, transcript, and peptide sequence files** as part of your submission.

| Student Name(s): | Click or tap here to enter text. |
| --- | --- |
| Student Email(s): | Click or tap here to enter text. |
| Faculty Instructor: | Click or tap here to enter text. |
| Course Name/Number: | Click or tap here to enter text. |
| College/University: | Click or tap here to enter text. |

### Project Details

| Project Species (e.g., *D. yakuba*) | Click or tap here to enter text. |
| --- | --- |
| NCBI Taxonomy ID (e.g., 7245) | Click or tap here to enter text. |
| NCBI Assembly ID (e.g., dyak_caf1) | Click or tap here to enter text. |
| Assembly Accession (e.g., GCA_000005975.1) | Click or tap here to enter text. |
| Genome Assembly (e.g., May 2011 (WUGSC dyak_caf1/DyakCAF1)) | Click or tap here to enter text. |
| Scaffold Name (e.g., chr3R) | Click or tap here to enter text. |
| Scaffold Accession (e.g., CM000160.2) | Click or tap here to enter text. |
| Gene ID in target species (e.g., dyak_Rheb) | Click or tap here to enter text. |

| Gene ID in *D. melanogaster* (e.g. dmel_Rheb) | Click or tap here to enter text. |
| --- | --- |
| Accession Number of Ortholog in *D. melanogaster* (e.g., NT_033777) | Click or tap here to enter text. |
| Chromosome of Ortholog in *D. melanogaster* | Click or tap here to enter text. |

| Date of Submission (YYYY-MM-DD) | Click or tap here to enter text. |
| --- | --- |

The data from this document will eventually be published, so we will need some contact information from you as well as permissions:

| Permanent email address (e.g., one you will probably use 5 years from now): | Click or tap here to enter text. |
| --- | --- |
| Alternative email address (optional): | Click or tap here to enter text. |
| Cell phone number (optional): | Click or tap here to enter text. |

If you choose to be a co-author(s), you will have to respond promptly to requests to read and approve the manuscript, and, as part of that review, you will also be required to validate some specific data within the manuscript (full instructions will be provided). We estimate that you would contribute 3-5 hours of your time to the manuscript preparation process.

If you want to be a co-author(s) on a publication, and we cannot reach you at the time the publication is ready for your review, you will no longer be a co-author(s) on the publication that arises from this data because you are not able to read and approve the manuscript.

If you would like to be a co-author(s), please enter your name(s) in the format you want it/them to be displayed as in publications in the table below. Note: If more than three students contribute to an individual gene annotation report as a group project, none of those students are eligible for co-authorship but the class as a whole will be acknowledged.

| Names you want on the publication (s) | Click or tap here to enter text. |
| --- | --- |
|  | Click or tap here to enter text. |
|  | Click or tap here to enter text. |

By submitting this report to the GEP, you acknowledge that you are allowing the data presented here to be published.

| ☐ | By checking this box, I/we understand that my instructor may submit my/our Annotation Report and supporting documentation to the Genomics Education Partnership (GEP). The Report contains my/our name(s) and contact information. The Partnership may use my/our work in a publication. To be a co-author on any possible publications, I need to reply promptly to an email from the GEP when the manuscript is ready for review; if I do not, there is a chance my name will not be included as a co-author. |
| --- | --- |

Navigation Pane

You will find it helpful to use the *Navigation* *Pane* function of Word while editing this document. It will be much easier to move around sections this way. Since Word versions vary in their layout, please follow the [Use the Navigation pane in Word](https://support.office.com/en-us/article/use-the-navigation-pane-in-word-394787be-bca7-459b-894e-3f8511515e55?ui=en-US&rs=en-US&ad=US) webpage to enable viewing of the navigation pane in your version of Word. Make sure to select the “Headings” tab within the pane to see the different sections in the document. Note that your version of MS Word might be different than what the website is portraying.

Entering Text Data

To enter text data, please click on the textbox with this font, and type your data. The prompt text should be cleared automatically when you start typing, but if it is not, please delete it and enter your data.

When entering coordinates: do not use commas to separate place values – this will make it easier to move data to the [Gene Model Checker](http://gander.wustl.edu/~wilson/genechecker/index.html). For example, use 10000 instead of 10,000.

Adding Images

To add images within the blue boxes, click the Picture icon (
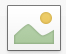
). This opens a file browser from which you can select your image. For more information, please refer to [this video tutorial](https://youtu.be/Up5z1mRsOqE?t=243). If you need to change the picture, right click and select “Change Picture,” and then click “From a File.”

Screenshot in GEP UCSC Genome Browser

Avoid taking a screenshot of the GEP UCSC Genome Browser. We have incorporated a way to export the image. Right-click within the genome browser, and select “View Image”. This should automatically take you to a new tab. If it doesn’t make sure that popups are enabled for the GEP UCSC Genome Browser from within your internet browser.

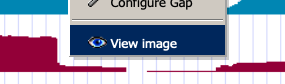

### A. Gene Report Form

**Inspect the region around your gene in *D. melanogaster.* Record the names of the nearest two protein coding genes upstream and the two downstream of your gene in *D. melanogaster* and in your target species including the *blastp* results that support your target species Gene Symbol. Note if your gene is nested within another gene, or another gene is nested within your gene, and indicate whether the genes are orthologs between the two species. If you do not have a nested/nesting gene, leave the “Nested Gene” column blank.**

|  |  | Most Upstream | Nearest Upstream | Nested Gene | Target Gene | Nearest Downstream | Most Downstream |
| --- | --- | --- | --- | --- | --- | --- | --- |
|  | *D. melanogaster* Gene Symbol | Symbol | Symbol | Symbol | Symbol | Symbol | Symbol |
|  | Strand in *D. melanogaster* | +/- | +/- | +/- | +/- | +/- | +/- |
|  | Target NCBI RefSeq Accession | ID | ID | ID | ID | ID | ID |
|  | Strand in  Target Species | +/- | +/- | +/- | +/- | +/- | +/- |
| BEST HIT | Accession | Accession | Accession | Accession | Accession | Accession | Accession |
|  | *D. melanogaster* Gene Symbol | ID | ID | ID | ID | ID | ID |
|  | E-value | e-value | e-value | e-value | e-value | e-value | e-value |
|  | Percent Identity | % | % | % | % | % | % |
| 2^nd^ BEST HIT^[1]^ | *D. melanogaster* Gene Symbol | Symbol | Symbol | Symbol | Symbol | Symbol | Symbol |
|  | E-value | e-value | e-value | e-value | e-value | e-value | e-value |
|  | Percent Identity | % | % | % | % | % | % |
|  | Orthologs? | YES/NO | YES/NO | YES/NO | YES/NO | YES/NO | YES/NO |

**Paste below an image of the genomic neighborhood for *D. melanogaster* from the GEP UCSC Genome Browser including both of the two upstream and two downstream genes.** Select the “default tracks” for the region, and then set a comparative genomics track (e.g., Drosophila Chain/Net) (if available) to “pack”.

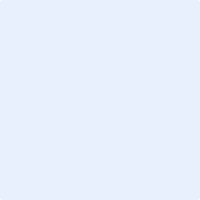

**Paste below an image of the genomic neighborhood of your target gene in the target species from the GEP UCSC Genome Browser. Be sure to include including both of the two nearest upstream and both of the two nearest downstream genes.** Select the “default tracks” for the region, and then set a comparative genomics track (e.g., Drosophila Chain/Net) (if available) to “pack”.

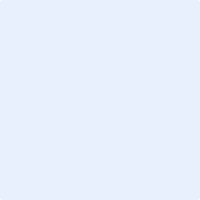

**Paste below a screenshot of the “Descriptions” panel of your *tblastn* results of the amino acid sequence of the *D. melanogaster* protein coding isoform for your gene against the genome assembly in your target species.**

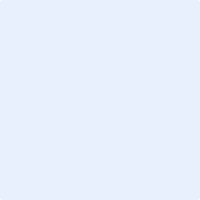

**Explain what evidence supports your hypothesis that you have located the correct genomic neighborhood in the target species, and you are, therefore, annotating the ortholog to the *D. melanogaster* gene.** Be sure to also explain any discrepancies found in the *blast* results or genomic neighborhood described above (e.g., regions of the protein are missing, synteny, number of exons).

Click or tap here to enter text.

### Consensus Sequence Errors Report Form

Only complete this section if you have identified errors in the project consensus sequence, otherwise move to the Coding Sequence (CDS) Report Form section (but do NOT delete this section).

**All of the coordinates reported in this section should be relative to the coordinates of the original genomic sequence.**

Location(s) in the project sequence with consensus errors:

| *locus* | *locus* | *locus* | *locus* |
| --- | --- | --- | --- |
| *locus* | *locus* | *locus* | *locus* |
| *locus* | *locus* | *locus* | *locus* |
| *locus* | *locus* | *locus* | *locus* |

##### 1. Evidence that supports the consensus errors postulated above

**Note:** Evidence that could be used to support the hypothesis of errors within the consensus sequence include CDS alignment with frame shifts or in-frame stop codons, multiple RNA-Seq reads with discrepant alignments compared to the project sequence, and multiple high-quality discrepancies in the Consed assembly.

Click or tap here to enter text.

If you need to use images to explain evidence of consensus errors, please paste screenshots below.

##### 2. Generate a VCF file which describes the changes to the consensus sequence

If your target species is available in the [Sequencer Updater tool](http://gander.wustl.edu/~wilson/sequence_updater/index.html), create a Variant Call Format (VCF) file that describes the changes to the consensus sequence you have identified above. **Paste a screenshot with the list of sequence changes below:**

Note: If your target species is not available in the Sequence Updater tool, see the VCF work-around instructions attached at the end of “How to annotate genes in other Drosophila species” instructions.

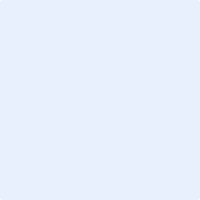

### B. Coding Sequence (CDS) Report Form

Number of isoforms in *D. melanogaster:* Click or tap here to enter text.

Number of isoforms in this project: Click or tap here to enter text.

**Note:** If more isoforms exist than there is space for in the table, please add more rows by going to the bottom-right-most cell and pressing tab.

**Complete the following table for all of the *D. melanogaster* isoforms in this project:**

| **Name(s) of unique isoform(s) in *D. melanogaster* based on coding sequence** | **List of isoforms with identical coding sequences in *D. melanogaster*** | **Select if coding isoform is likely present in target species.^[2]^** |
| --- | --- | --- |
| Isoform | Identical isoforms | ☐ |
| Isoform | Identical isoforms | ☐ |
| Isoform | Identical isoforms | ☐ |
| Isoform | Identical isoforms | ☐ |
| Isoform | Identical isoforms | ☐ |
| Isoform | Identical isoforms | ☐ |
| Isoform | Identical isoforms | ☐ |
| Isoform | Identical isoforms | ☐ |
| Isoform | Identical isoforms | ☐ |
| Isoform | Identical isoforms | ☐ |
| Isoform | Identical isoforms | ☐ |

Is there strong evidence (e.g., extended RNA-Seq, high quality splice sites, conservation across multiple species) for distinct protein coding isoforms present in your species that are not found in *D. melanogaster*?	☐ Yes	☐ No

If yes, how many?	Enter number

If yes, create additional isoform reports for those coding sequences and name the isoforms “-PAA,” “-PAB,” etc. (e.g., dyak_Rheb-PAA).

**Note:** For isoforms with identical coding sequences, you only need to complete the Isoform Report Form for one of these isoforms (i.e., using the name of the isoform listed in the left column of the table above). However, you should **generate GFF, transcript, and peptide sequence files for ALL isoforms**, regardless of whether they have identical coding sequences as other isoforms. If an isoform is not present in your target species, do not complete an isoform report for that isoform.

When copying sections for additional isoforms, copy the entire section first, and then populate it. (Copying individual pages will alter the Navigation Pane.)

#### Missing CDS Isoforms

Missing Isoforms in your target species: ☐ Yes	☐ No

If you think all CDS isoforms within the D. melanogaster genome exist within your target species, please select “No”. However, if you believe there are CDS isoforms within the D. melanogaster genome that do not exist within your target species, please select “Yes” and provide explanations with screenshots and text below.

Please do not try to create page-breaks or section breaks within your explanation.

#### [TYPE GENE ISOFORM NAME HERE] – Coding Isoform Report Form

Complete this report form for each unique isoform listed in the table above (copy and paste to create as many copies of this Isoform Report Form as needed; ***when copying sections for additional isoforms, copy the entire blank section first, and then populate it.***)***^[3]^***:

##### Gene Model Checker Checklist

Enter the coordinates for the coding exons below (e.g., 100-200). Add more rows if needed. All rows do not need to be populated. Remember to leave out commas when typing the coordinates. If more CDSs exist, please add more rows by going to the bottom-right-most cell and pressing tab.

| **CDS Number** | **Acceptor Phase** | **CDS** **Coordinates** | **Donor Phase** |
| --- | --- | --- | --- |
| CDS 1 | 0 | Start - Stop |  |
| CDS 2 |  | Start - Stop |  |
| CDS 3 |  | Start - Stop |  |
| CDS 4 |  | Start - Stop |  |
| CDS 5 |  | Start - Stop |  |
| CDS 6 |  | Start - Stop |  |
| CDS 7 |  | Start - Stop |  |
| CDS 8 |  | Start - Stop |  |
| CDS 9 |  | Start - Stop |  |
| CDS 10 |  | Start - Stop |  |
| CDS 11 |  | Start - Stop |  |
| CDS 12 |  | Start - Stop |  |
| CDS 13 |  | Start - Stop |  |
| CDS 14 |  | Start - Stop |  |
| CDS 15 |  | Start - Stop |  |
| CDS 16 |  | Start - Stop |  |
| CDS 17 |  | Start - Stop |  |
| CDS 18 |  | Start - Stop |  |

Stop Codon Coordinates (e.g., 201-203): Start – Stop

Enter the coordinates of your final gene model for this isoform into the [Gene Model Checker](http://gander.wustl.edu/~wilson/genechecker/index.html) and **paste a screenshot of the checklist results and data entry section below:**

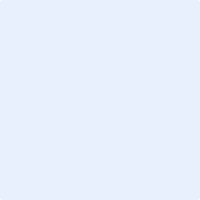

**Note:** For projects with consensus sequence errors, report the exon coordinates relative to the **original project sequence**. Include the VCF file you have generated above when you submit the gene model to the Gene Model Checker. The Gene Model Checker will use this VCF file to automatically revise the submitted exon coordinates.

##### View the gene model on the GEP UCSC Genome Browser

Use the custom track feature from the Gene Model Checker to capture an image of your gene model shown on the Genome Browser for your project.

In the checklist results of the Gene Model Checker, click on the
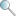
 icon next to “Number of coding exons matched ortholog,” and a new window will open showing the Genome Browser view of this region. Your gene model will be shown under the track title “Custom Gene Model.”

Select the “default tracks” for the region, and then set the following evidence tracks (if available) to “pack.”

1. at least one transcript prediction track (e.g., TransDecoder Transcripts, modENCODE Cufflinks Transcripts)
2. at least one splice-site prediction track (e.g., Splice Junctions, modENCODE TopHat Junctions)
3. a comparative genomics track (e.g., Drosophila Chain/Net)

**Paste an image of your gene model as shown on the Genome Browser (including the above listed tracks) below:**

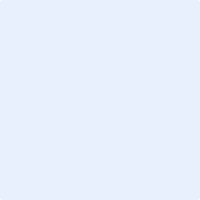

##### Alignment between the submitted model and the *D. melanogaster ortholog*

**Paste an image of the protein alignment** generated by the Gene Model Checker (available through the “**View protein alignment**” link under the “Dot Plot” tab) **below:**

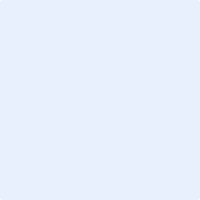

##### Dot plot between the submitted model and the *D. melanogaster ortholog*

**Paste an image of the dot plot** of your submitted model against the putative *D. melanogaster* ortholog (generated by the Gene Model Checker). **Provide an explanation for any anomalies** on the dot plot (e.g., large gaps, regions with no sequence similarity) in the textbox below the image. Also propose why you think they might be valid. You can include screen shots to illustrate the logic of your explanation below the text box.

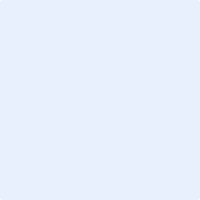

Click or tap here to enter text.

**Note: Large vertical and horizontal gaps** near exon boundaries in the dot plot often indicate that an incorrect splice site might have been picked. Please re-examine these regions and provide a detailed justification as to why you have selected this particular set of donor and acceptor sites.

If you need to use images to explain anomalies within your dot plot, please paste screenshots here.

### C. Transcription Start Site (TSS) Report Form

**Note:** Complete this section if you have annotated the Coding Isoforms for the gene above. This section is **OPTIONAL** based on your instructor’s expectations. If more isoforms exist, please add more rows by going to the bottom-right-most cell and pressing tab.

| **Name(s) of transcript isoform(s) with unique TSS in *D. melanogaster*** | **List of transcript isoforms with identical TSS in *D. melanogaster*** |
| --- | --- |
| Isoform | Transcript Isoforms |
| Isoform | Transcript Isoforms |
| Isoform | Transcript Isoforms |
| Isoform | Transcript Isoforms |
| Isoform | Transcript Isoforms |
| Isoform | Transcript Isoforms |
| Isoform | Transcript Isoforms |
| Isoform | Transcript Isoforms |
| Isoform | Transcript Isoforms |
| Isoform | Transcript Isoforms |
| Isoform | Transcript Isoforms |

#### Missing Transcription Start Sites

Missing TSS in your target species: ☐ Yes	☐ No

If you think all TSS within the D. melanogaster genome exist within your target species, please select “No”. However, if you believe there are TSS within the D. melanogaster genome that do not exist within your target species, please select “Yes” and provide explanations with screenshots and text below.

Please do not try to create page-breaks or section breaks within your explanation.

#### **[TYPE TRANSCRIPT ISOFORM NAME HERE] – TSS Annotation** *Report Form*

Complete this report form for each unique isoform listed in the table above (copy and paste to create as many copies of this Isoform Report Form as needed; ***when copying sections for additional isoforms, copy the entire section first, and then populate it.***):

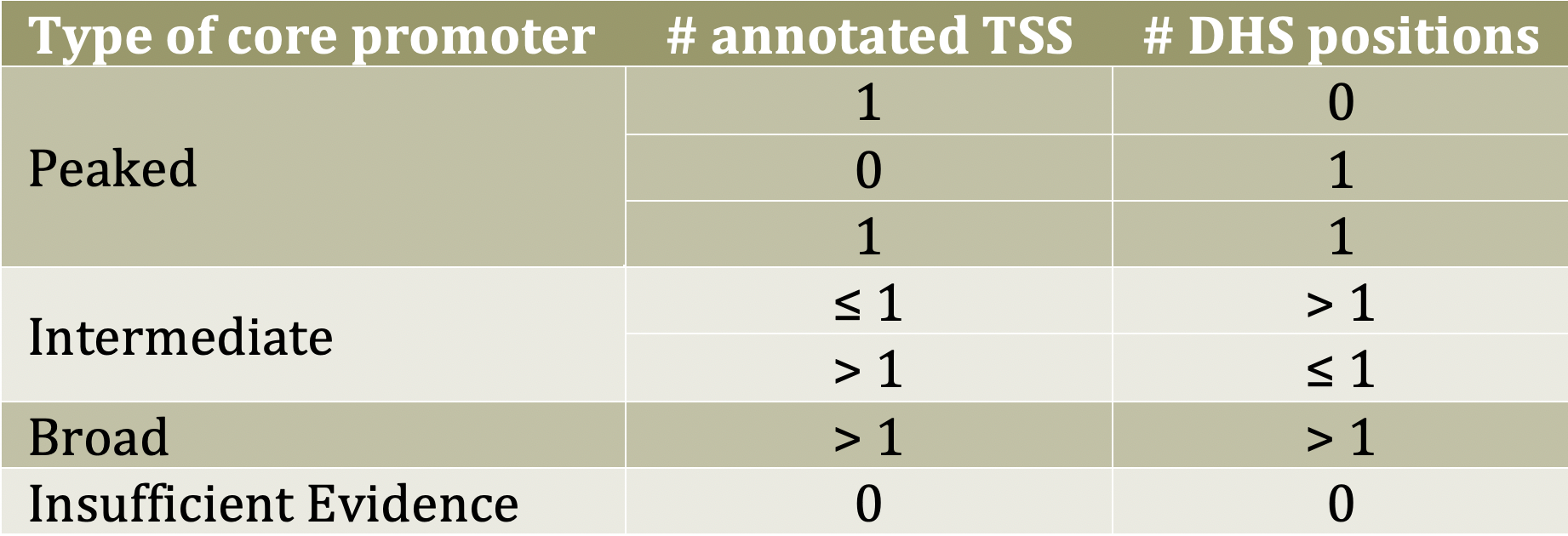

Type of core promoter in *D. melanogaster*:

☐ Peaked

☐ Intermediate

☐ Broad

☐ Insufficient Evidence

Coordinate(s) of the TSS position(s) or likely region(s) for the TSS (enter “NA” if data not applicable). Also, enter the reason you think these are the best coordinates in the bottom cell of the table:

| Based on blastn alignment | Proposed coordinate(s) |
| --- | --- |
| Based on core promoter motifs (e.g., Inr) | Proposed coordinate(s) |
| Based on RNA-seq, TopHat, Cufflinks tracks | Proposed coordinate(s) |
| Based on Conservation tracks | Proposed coordinate(s) |
| Based on other evidence (please specify) | Proposed coordinate(s) |
| Reason for proposed coordinate(s). | |

**Note:** If the *blastn* alignment for the initial transcribed exon is a partial alignment, you can **extrapolate the TSS position** based on the number of nucleotides that are missing from the beginning of the exon. (Enter “Insufficient evidence” if you cannot determine the TSS position based on the available evidence.)

What do you believe is/are the best coordinate(s) or coordinate range for the TSS position(s) for this isoform given all the evidence?

Best proposed coordinate(s)

Provide an explanation if the best TSS coordinate(s) or coordinate range is inconsistent with at least one of the evidence types specified above. You can add screen shots to further explain your logic below the textbox:

Click or tap here to enter text.

Perform a *blastn* alignment of the initial transcribed exon in *D. melanogaster* against the genomic sequence of your target species. Remember to increase sensitivity by changing the “Word size” parameter to **7**, the “Match/Mismatch Scores” to “**1, -1**”, the “Gap Costs” to “**Existence: 2 Extension: 1**”, and **uncheck the filter for “Low complexity regions.”** **Paste a screenshot of the *blastn* alignment below:**

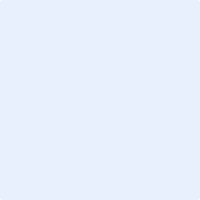

**Paste an image of the Genome Browser region surrounding the putative TSS (±300 bp) with the following evidence tracks:**

1. RNA-Seq Alignment Summary
2. Splice Junctions
3. Short Match results for the Inr motif (TCAKTY)
4. A comparative genomics track (e.g., Drosophila Chain/Net)

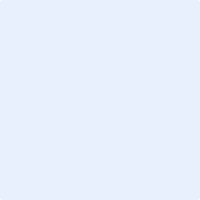

##### *Search for core promoter motifs*

The consensus sequences for the *Drosophila* core promoter motifs are available at <http://gander.wustl.edu/~wilson/core_promoter_motifs.html>

Use the "Short Match" functionality in the GEP UCSC Genome Browser to search for each of the core promoter motifs listed below **in the region surrounding the TSS (±300 bp) in your project and in the *D. melanogaster* ortholog**. For TSS annotations where you can only define a TSS search region, you should report all motif instances ±300 bp of either edge of your search region.

###### Coordinates of the motif search region

Your project (e.g., scaffold_1234:1000-1600): Start - Stop

Orthologous region in *D. melanogaster*: Start - Stop

Record the **orientation and the start coordinate** (e.g., +10000) of each motif match below. (Enter "**NA**" if there are no motif instances within the search region.)

**Note:** Highlight (in yellow) the motif instances that support the TSS annotation(s) above.

| Core promoter motif | Your project | *D. melanogaster* |
| --- | --- | --- |
| BRE^u^ | Start coordinate | Start coordinate |
| TATA Box | Start coordinate | Start coordinate |
| BRE^d^ | Start coordinate | Start coordinate |
| Inr | Start coordinate | Start coordinate |
| MTE | Start coordinate | Start coordinate |
| DPE | Start coordinate | Start coordinate |
| Ohler_motif1 | Start coordinate | Start coordinate |
| DRE | Start coordinate | Start coordinate |
| Ohler_motif5 | Start coordinate | Start coordinate |
| Ohler_motif6 | Start coordinate | Start coordinate |
| Ohler_motif7 | Start coordinate | Start coordinate |
| Ohler_motif8 | Start coordinate | Start coordinate |

### D. Transcript Report Form

**Note:** Complete this section if you have annotated the transcript for the gene above. This section is **OPTIONAL** based on your instructor’s expectations. If more isoforms exist, please add more rows by going to the bottom-right-most cell and pressing tab.

| **Name(s) of transcript isoform(s) in *D. melanogaster*** | **Do you think this isoform is present in your target species (select if Yes)?** |
| --- | --- |
| Transcript isoform | ☐ |
| Transcript isoform | ☐ |
| Transcript isoform | ☐ |
| Transcript isoform | ☐ |
| Transcript isoform | ☐ |
| Transcript isoform | ☐ |
| Transcript isoform | ☐ |
| Transcript isoform | ☐ |

Is there strong evidence for distinct transcript isoforms present in your species that are not found in *D. melanogaster* (yes/no)?	☐ Yes	☐ No

If yes, how many?	*Enter number*

*Note that if you have a novel coding region you should also have at least one novel transcript.

If yes, create additional transcript isoform reports for those transcript sequences and name the isoforms “-RAA,” “-RAB,” etc. (e.g. dyak_Rheb-RAA).

*If these new transcripts include novel CDS’s, then make sure to match the isoform naming used for the coding region above.

#### Missing Transcript Isoforms

Missing Isoforms in your target species: ☐ Yes	☐ No

If you think all transcript isoforms within the D. melanogaster genome exist within your target species, please select “No”. However, if you believe there are transcript isoforms within the D. melanogaster genome that do not exist within your target species, please select “Yes” and provide explanations with screenshots and text below.

Please do not try to create page-breaks or section breaks within your explanation.

#### [TYPE TRANSCRIPT NAME HERE] - Transcript Isoform Report Form

Complete this report form for each unique isoform listed in the table above (copy and paste to create as many copies of this Isoform Report Form as needed; ***when copying sections for additional isoforms, copy the entire section first, and then populate it.***):

Enter the coordinates for the **transcribed exons** below (e.g., 100-200). Add more rows if needed. Also mark whether the exon contains UTRs (UnTranslated Regions). Make sure to enter the coordinates without commas.

| **Exon #** | **Acceptor Phase** | **Coordinates** | **Donor Phase** | **Check box if contains UTR** |
| --- | --- | --- | --- | --- |
| Exon 1 | 0 | Start - Stop |  | ☐ |
| Exon 2 |  | Start - Stop |  | ☐ |
| Exon 3 |  | Start - Stop |  | ☐ |
| Exon 4 |  | Start - Stop |  | ☐ |
| Exon 5 |  | Start - Stop |  | ☐ |
| Exon 6 |  | Start - Stop |  | ☐ |
| Exon 7 |  | Start - Stop |  | ☐ |
| Exon 8 |  | Start - Stop |  | ☐ |
| Exon 9 |  | Start - Stop |  | ☐ |
| Exon 10 |  | Start - Stop |  | ☐ |
| Exon 11 |  | Start - Stop |  | ☐ |
| Exon 12 |  | Start - Stop |  | ☐ |
| Exon 13 |  | Start - Stop |  | ☐ |
| Exon 14 |  | Start - Stop |  | ☐ |
| Exon 15 |  | Start - Stop |  | ☐ |
| Exon 16 |  | Start - Stop |  | ☐ |
| Exon 17 |  | Start - Stop |  | ☐ |
| Exon 18 |  | Start - Stop |  | ☐ |

Use the [Gene Model Checker](http://gander.wustl.edu/~wilson/genechecker/index.html) and **make sure to click “yes” for “Annotated Untranslated Regions?”.**

Enter the coordinates of your final gene model for this isoform into the [Gene Model Checker](http://gander.wustl.edu/~wilson/genechecker/index.html) and **paste an image of the checklist results and data entry section below:**

**Note:** For projects with consensus sequence errors, report the exon coordinates relative to the **original project sequence**. Include the VCF file you have generated above when you submit the gene model to the Gene Model Checker. The Gene Model Checker will use this VCF file to automatically revise the submitted exon coordinates.

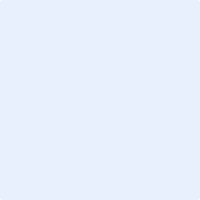

##### View the gene model on the GEP UCSC Genome Browser

Use the custom track feature from the Gene Model Checker to capture an image of your gene model shown on the Genome Browser for your project.

In the checklist results of the Gene Model Checker, click on the
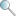
icon next to “Number of coding exons matched ortholog,” and a new window will open showing the Genome Browser view of this region. Your gene model will be shown under the track title “Custom Gene Model.”

Select the “default tracks” for the region, and then set the following evidence tracks (if available) to “pack.”

1. at least one transcript prediction track (e.g., TransDecoder Transcripts, modENCODE Cufflinks Transcripts)
2. at least one splice-site prediction track (e.g., Splice Junctions, modENCODE TopHat Junctions)
3. a comparative genomics track (e.g., Drosophila Chain/Net)

**Paste an image of your gene model as shown on the Genome Browser (including the above listed tracks) below:**

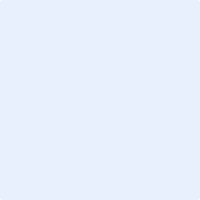

##### *Alignment between the submitted model and the D. melanogaster ortholog*

Show an alignment between the nucleotide sequence for your gene model and the nucleotide sequence from the putative *D. melanogaster* ortholog. You can generate a new alignment using the “Align two or more sequences” feature (*bl2seq*) at the NCBI BLAST web site. Remember to increase sensitivity by changing the “Word size” parameter to **7**, the “Match/Mismatch Scores” to “**1, -1**”, the “Gap Costs” to “**Existence: 2 Extension: 1**”, and **uncheck the filter for “Low complexity regions”.**

If no alignment results, record “NA”. **Paste a screenshot of the nucleotide alignment below:**

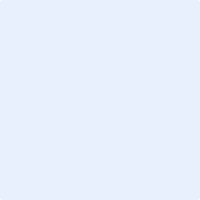

### E. Preparing the Project for Submission

For **each gene**, you should prepare the project GFF, transcript, and peptide sequence files for **ALL** isoforms along with this report. You can combine the individual files of one type (e.g., GFF) for all isoforms for one gene generated by the Gene Model Checker into a single file using the [Annotation Files Merger](http://gander.wustl.edu/~wilson/submissionhelper/index.php). You should have a total of three files, one GFF, one transcript, and one peptide for each gene.

Name the files using **species_gene.filetype**

So, if you have a PEP file for *Rheb* in *D. yakuba*, the filename would be “**dyak_Rheb.pep**”. The same goes for the other file types (FASTA and GFF).

For only projects with multiple errors in the consensus sequence, you should combine all the VCF files into a single project VCF file using the [Annotation Files Merger](http://gander.wustl.edu/~wilson/submissionhelper/index.php). **Paste a screenshot (generated by the Annotation Files Merger) with all the consensus sequence errors you have identified in your project.**

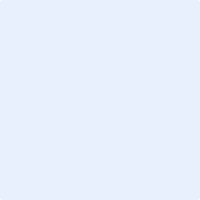

1. Not an isoform of best hit
2. If this isoform is novel, does not exist in *D. melanogaster*, leave this blank.
3. Note: If you are not doing the TSS and full transcript annotations skip to section E. Preparing Project for Submission.
